# Cryptic pathogen-sugar interactions revealed by universal saturation transfer analysis

**DOI:** 10.1101/2021.04.14.439284

**Authors:** Charles J. Buchanan, Ben Gaunt, Peter J. Harrison, Yun Yang, Jiwei Liu, Aziz Khan, Andrew M. Giltrap, Audrey Le Bas, Philip N. Ward, Kapil Gupta, Maud Dumoux, Sergio Daga, Nicola Picchiotti, Margherita Baldassarri, Elisa Benetti, Chiara Fallerini, Francesca Fava, Annarita Giliberti, Panagiotis I. Koukos, Abirami Lakshminarayanan, Xiaochao Xue, Georgios Papadakis, Lachlan P. Deimel, Virgínia Casablancas-Antràs, Timothy D.W. Claridge, Alexandre M.J.J. Bonvin, Quentin J. Sattentau, Simone Furini, Marco Gori, Jiandong Huo, Raymond J. Owens, Christiane Schaffitzel, Imre Berger, Alessandra Renieri, GEN-COVID Multicenter Study, James H. Naismith, Andrew Baldwin, Benjamin G. Davis

## Abstract

Many host pathogen interactions such as human viruses (including non-SARS-coronaviruses) rely on attachment to host cell-surface glycans. There are conflicting reports about whether the Spike protein of SARS-CoV-2 binds to sialic acid commonly found on host cell-surface N-linked glycans. In the absence of a biochemical assay, the ability to analyze the binding of glycans to heavily-modified proteins and resolve this issue is limited. Classical Saturation Transfer Difference (STD) NMR can be confounded by overlapping sugar resonances that compound with known experimental constraints. Here we present ‘universal saturation transfer analysis’ (uSTA), an NMR method that builds on existing approaches to provide a general and automated workflow for studying protein-ligand interactions. uSTA reveals that B-origin-lineage-SARS-CoV-2 spike trimer binds sialoside sugars in an ‘end on’ manner and modelling guided by uSTA localises binding to the spike N-terminal domain (NTD). The sialylated-polylactosamine motif is found on tetraantennary human N-linked-glycoproteins in deeper lung and may have played a role in zoonosis. Provocatively, sialic acid binding is abolished by mutations in some subsequent SARS-CoV-2 variants-of-concern. A very high resolution cryo-EM structure confirms the NTD location and ‘end on’ mode; it rationalises the effect of NTD mutations and the structure-activity relationship of sialic acid analogues. uSTA is demonstrated to be a robust, rapid and quantitative tool for analysis of binding, even in the most demanding systems.

**Extended Abstract:** The surface proteins found on both pathogens and host cells mediate entry (and exit) and influence disease progression and transmission. Both types can bear host-generated post-translational modifications such as glycosylation that are essential for function but can confound biophysical methods used for dissecting key interactions. Several human viruses (including non-SARS-coronaviruses) attach to host cell-surface *N*-linked glycans that include forms of sialic acid (sialosides). There remains, however, conflicting evidence as to if or how SARS-associated coronaviruses might use such a mechanism. Here, we demonstrate quantitative extension of ‘saturation transfer’ protein NMR methods to a complete mathematical model of the magnetization transfer caused by interactions between protein and ligand. The method couples objective resonance-identification via a deconvolution algorithm with Bloch-McConnell analysis to enable a structural, kinetic and thermodynamic analysis of ligand binding beyond previously-perceived limits of exchange rates, concentration or system. Using an automated and openly available workflow this ‘universal saturation transfer’ analysis (uSTA) can be readily-applied in a range of even heavily-modified systems in a general manner to now obtain quantitative binding interaction parameters (K_D_, k_Ex_). uSTA proved critical in mapping direct interactions between natural sialoside sugar ligands and relevant virus-surface attachment glycoproteins – SARS-CoV-2-spike and influenza-H1N1-haemagglutinin variants – by quantitating ligand signal in spectral regions otherwise occluded by resonances from mobile protein glycans (that also include sialosides). In B- origin-lineage-SARS-CoV-2 spike trimer ‘end on’-binding to sialoside sugars was revealed contrasting with ‘extended surface’-binding for heparin sugar ligands; uSTA-derived constraints used in structural modelling suggested sialoside-glycan binding sites in a beta-sheet-rich region of spike N-terminal domain (NTD). Consistent with this, uSTA-glycan binding was minimally- perturbed by antibodies that neutralize the ACE2-binding domain (RBD) but strongly disrupted in spike from the B1.1.7/alpha and B1.351/beta variants-of-concern, which possess hotspot mutations in the NTD. Sialoside binding in B-origin-lineage-NTD was unequivocally pinpointed by cryo-EM to a site that is created from residues that are notably deleted in variants (e.g. H69,V70,Y145 in alpha). An analysis of beneficial genetic variances in cohorts of patients from early 2020 suggests a model in which this site in the NTD of B-origin-lineage-SARS-CoV-2 (but not in alpha/beta-variants) may have exploited a specific sialylated-polylactosamine motif found on tetraantennary human *N*-linked-glycoproteins in deeper lung. Together these confirm a novel binding mode mediated by the unusual NTD of SARS-CoV-2 and suggest how it may drive virulence and/or zoonosis via modulation of glycan attachment. Since cell-surface glycans are widely relevant to biology and pathology, uSTA can now provide ready, quantitative, widespread analysis of complex, host-derived and post-translationally modified proteins with putative ligands relevant to disease even in previously confounding complex systems.

## Introduction

Binding mediated by sialosides present in glycans that are anchored to human cells is central to cell-cell communication in human physiology and is at the heart of many host-pathogen interactions. One of the most well-known is that of influenza virus which binds to sialosides with its haemagglutinin (HA or H) protein and cleaves off sialic acid from the infected cell with its neuraminidase (NA or N) protein; HxNx variants of influenza with different HA or NA protein types have a profound effect on zoonosis and pathogenicity (*1*).

The MERS(*2*) virus, which is related to SARS-CoV-1/2 has been shown to exploit cell-surface sugar sialosides(*2–6*) as part of an attachment strategy. Both SARS-CoV-1(*7–9*) and SARS-CoV-2(*10, 11*) are known to gain entry to host cells through the use of receptor-binding domains (RBDs) of their respective spike proteins that bind human cell-surface protein ACE2, but whether these viruses engage sialosides as part of the infection cycle has, despite predictions(*6, 12*), remained unclear. This is because preliminary reports as to whether complex sialosides are or are not bound are contradictory and format-dependent (*13–15*). Glycosaminoglycans on proteoglycans, such as heparin, have been identified as a primary cooperative glycan attachment point.(*14, 16*) All studies reporting sugar binding have so far implicated binding sites in or close to the RBD of the spike protein. Surprisingly, the N-terminal domain (NTD, **Supplementary Figure S1**), which has a putative glycan binding fold (*10, 17, 18*) and binds sialosides in other non-SARS coronaviruses (including MERS), has been overlooked. The NTD has no confirmed function in SARS-CoV-2. The unresolved role of host cell surface sialosides for this pathogen has been noted as an important open question.(*16*) The hypothesized roles for such binding(*19*) in both virulence(*20*) and zoonosis(*1*) means that there is an urgent need for precise, quantitative and robust methods for analysis.

In principle, magnetization transfer in protein NMR could meet this need, as it can measure ligand binding in native state without the need for additional labelling or modification of either ligand or protein (e.g. attachment to surface or sensor) (**Supplementary Figure S2,3** and see **Supplementary Note 1** for more details). ‘Saturation transfer difference’ (STD)(*21*), which has been used widely to gauge qualitative ligand•protein interactions (*22*), detects the transfer of magnetization while they are bound via ‘cross relaxation’, a widely-exploited mechanism for molecular and biomolecular characterization via NMR nuclear Overhauser effect (NOE) experiments. In reality, complex, highly-modified protein systems have proven difficult to analyze in a quantitative manner by using current methods for several reasons. First, mammalian proteins (or those derived by pathogens from expression in infected mammalian hosts) often bear large, highly-mobile glycans. Critically, in the case of glycoproteins, such as SARS-CoV-2-spike that may themselves bind glycans, this leads to contributions to protein NMR spectra that overlap with putative ligand resonances thereby obscuring needed signal. Second, the NMR spectra of glycan ligands are themselves complex, comprising many overlapped resonances as multiplets. This can limit the accurate determination of signal intensities. Finally, STD is commonly described as limited to specific kinetic regimes and/or ligand-to-protein binding equilibrium position.(*23*) As a result, many regimes and systems have therefore been considered inaccessible to STD.

Using a rigorous theoretical description, coupled with a computational approach based on a Bayesian deconvolution algorithm to objectively and accurately extract signal from all observed resonances, we have undertaken a complete reformulation of magnetization/‘saturation’ transfer (**Supplementary Figures S4, S5, S6, S8**). This approach reliably and quantitatively determines precise binding rates (k_on_, k_off_, k_ex_), constants (K_D_) and interaction ‘maps’ across a full range of regimes (**Supplementary Figure S4**) including previously intractable systems. We demonstrate universal saturation transfer analysis (uSTA) on characterized model ligand-protein systems (see **Supplementary Note 2** for details). We show that it can identify cryptic sialic acid binding in the hemagglutinin HA/H protein of the H1N1 strain of influenza. We have then applied uSTA to resolve the issue of sugar binding by SARS-CoV-2, determining that the B-origin-lineage SARS-CoV-2- spike protein does indeed bind sialosides and does so in an ‘end-on’ mode. Modelling guided by constraints from uSTA identified spike NTD domain as the binding region. Subsequent analysis by high resolution cryo-EM confirms these findings and identifies the key elements of recognition. The binding site is part of a mutational hotspot and is clearly subject to evolutionary modulation in emerging variants. Our results now suggest that recognition of host cell-surface glycans and the modulation of binding by spike-to-sialosides plays a role in zoonosis, infection, disease progression and disease evolution.

## Results

### Design of uSTA based on a Comprehensive Treatment of Ligand•Protein Magnetization Transfer

While using existing STD methodology to study the interaction between the SARS-CoV-2 SPIKE protein and sialosides, we noted several challenges that resulted in the development of uSTA (see **Supplementary Note 3** for more details). Our theoretical analyses (**Supplementary Figure S4**) suggested that many common assumptions or limits that are thought to govern the applicability of magnetization transfer might in fact be circumvented and we set out to devise a complete treatment that might accomplish this (**Supplementary Figures S5,S6,S8**). This resulted in five specific methodological changes that resulted in a more sensitive, accurate, quantitative and general method for studying the interactions between biomolecules and ligands (summarized in **Supplementary Figure S5,S8** and discussed in detail in **Supplementary Note 4**).

First, we noted the discrepancy between *K*_D_s determined using existing STD methods, and those obtained using other biophysical methods.(*24*) We performed a theoretical analysis using the Bloch-McConnell equations, a rigorous formulation for studying the evolution of magnetization in exchanging systems that has been widely used to analyze CEST(*25, 26*), DEST(*27*) and CPMG (*28*) NMR data to describe protein motion. This analysis allowed us to not only explain this discrepancy, but also enabled fitting of data to give accurate k_on_, k_off_, *K*_D_ values for protein-ligand interactions that were in excellent accord with alternative measurements (**Figures 1G, 4A,C** and **Supplementary Figure 13**), and demonstrate that the range of k_on_, k_off_ in which the experiment is applicable is far wider than previously recognized (**Supplementary Figure 4**). Second, in mammalian proteins, contributions from glycans on the surface of the protein could not be removed from the spectrum using relaxation filters used in ‘epitope mapping’(*29*) without compromising the sensitivity of the experiment. We addressed this instead by applying baseline subtraction using data obtained from a protein-only sample. Third, the magnetization transfer and hence the sensitivity of the experiment will be higher when the excitation frequency of the saturation pulse is close to a maximum in the protein NMR spectrum. Providing that any ligand resonances are outside of the bandwidth of the pulse, and a ‘ligand only’ subtraction is applied, the magnetization transfer can be maximised. With this, the response for a given protein-ligand system in fact becomes invariant to the excitation frequency used (**Supplementary Figures S9, S10**). Fourth, in complex molecules, such as sialosides, NMR spectra are crowded and overlapped. To reliably obtain magnetization transfer measurements at all points in the ligand, we developed a peak-picking algorithm based on earlier work(*30*) that can automate the process, returning a list of peak locations and a simulated NMR experiment that can be directly compared to the data. The locations of the peaks matches exactly the locations for multiplets determined using standard multi- dimensional approaches used for resonance assignment (see **Figures 1B, 1D, 2D, 3A** for examples – see also overlaps in all subsequent uSTA analyses and **Supplementary Table S7**).

**Figure 1.**
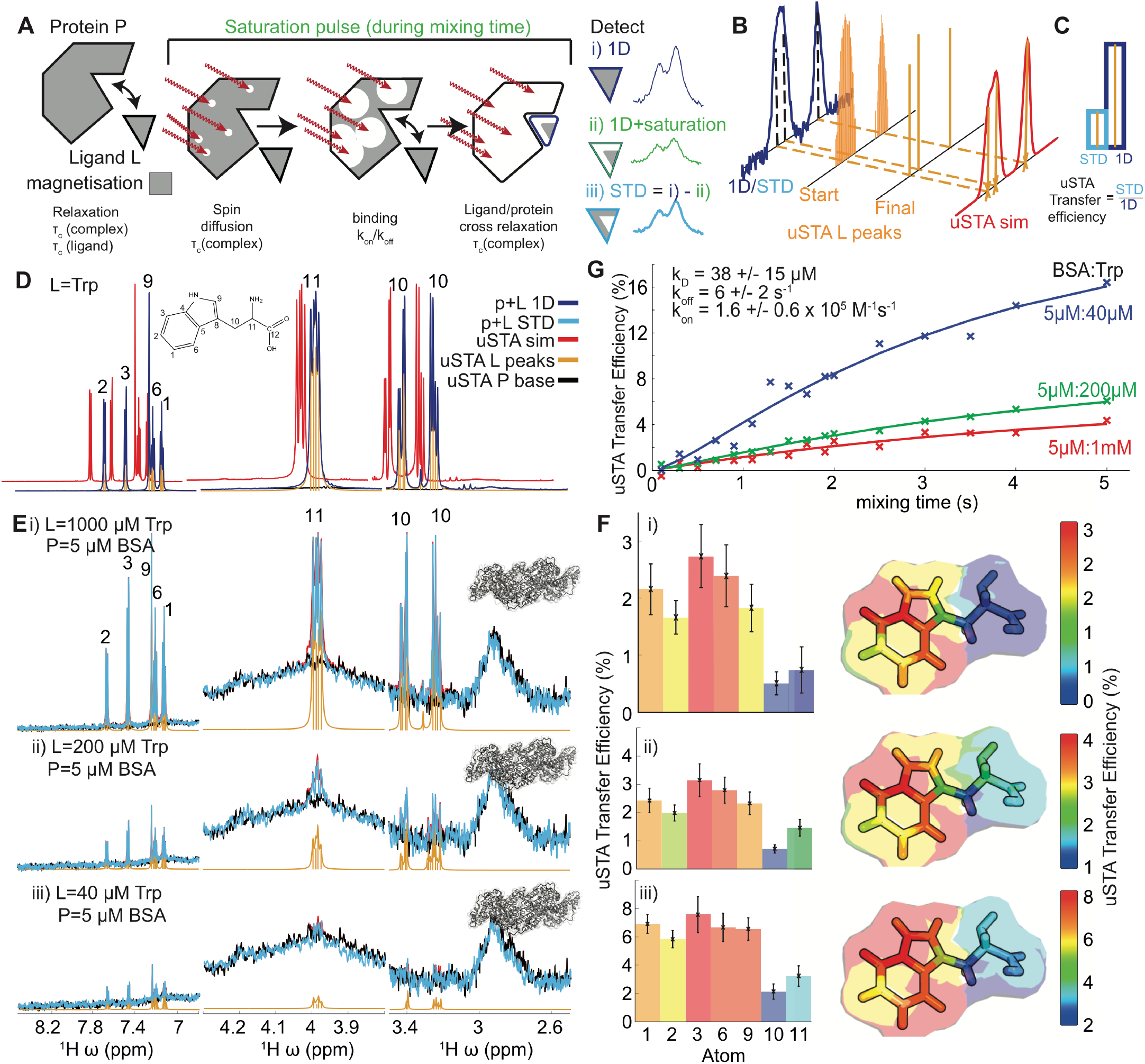
Development of the uSTA Method. **A**),**B**): Schematic of the process for uSTA that exploits comprehensive numerical analysis of multiple cross-relaxation parameters as well as ligand binding kinetics (**A**) using full and automatically-quantified signal intensities in NMR spectra (**B**) and calculates per-resonance transfer efficiencies (**C**). **B**): Signal analysis determined the number of peaks that can give rise to the signal, and returned simulated spectrum by convolving these with peak shape function. The precise peak positions returned are in excellent agreement with the known positions of resonances identified using conventional means. **C**): Magnetization transfer NMR experiments compare two 1D NMR spectra, where the second involves a specific saturation pulse that aims to ‘hit’ the protein but ‘miss’ the ligand in its excitation. To accomplish this the 1D spectrum is acquired with the saturation pulse held off resonance at -35 ppm such that it will not excite protons in either ligand or protein (labelled ‘1D’). The uSTA method requires these two spectra to be analysed as described in (**B**), in pairs, one that contains the raw signal, and the second that is the difference between the two. We define the ‘transfer efficiency’ as the fractional signal that has passed from the ligand to the protein. **D**)**,E),F)**: Application of uSTA to study the interaction between bovine serum albumin (BSA) and L-Tryptophan (Trp). In **D** the 1D ^1^H-NMR spectrum of the mixture at 200 µM Trp and 5 µM BSA (= P+L, **blue**) is dominated by ligand but yet the ligand (L) and protein (P) can still be deconvolved using universal deconvolution, using a reference obtained from a sample containing protein only. This reveals contributions from individual multiplets originating from the ligand (**yellow**) and the protein only baseline (**black**) allowing precise recapitulation of the sum (**red**). In **E** application of universal deconvolution to STD spectra with varying concentrations of tryptophan allows uSTA analysis using ligand resonances identified in **D**. This, in turn allows signal intensity in the STD spectrum (P+L STD, **light blue**) to be determined with high precision. While signal-to-noise in the STD increases considerably with increasing ligand concentration, the measured atom-specific transfer efficiencies as determined by uSTA are consistent (**F, left bar charts** and **right transfer efficiency binding ‘maps’**), showing that the primary contact between protein and ligand occurs on the C4a-C7a bridge of the indole aromatic ring. Application of the same uSTA workflow also allowed precise determination of even weakly binding sugar ligand trehalose (Glc-α1,1α-Glc) to *E. coli* trehalose repressor TreR. Again, the uSTA allows determination of transfer efficiencies with atom-specific precision (see **Supplementary Figure S3F**). **G**): Quantitative analysis of the STD build up curves using a modified set of Bloch-McConnell equations that account for binding and cross relaxation allow us to determine thermodynamic and kinetic parameters that describe the BSA•Trp interaction, *K*_D_, k_on_, and k_off_. The values obtained are indicated and are in excellent accord with those obtained by other methods (*24*). Errors come from a boot-strapping procedure (see **Methods**).

Finally, the uSTA software allowed the combination of intensity from scalar coupled multiplets following a user-inputted assignment, to provide ‘per resonance’ measures of saturation transfer. These are provided as-is, and also as <1/r^6^> interpolated ‘binding maps’ that represent the interaction on nearby heteroatoms allowing ready visual inspection of the binding pose of a molecule.

### Testing of uSTA in Model Systems

The uSTA method (**Figure 1A-C**) was tested first in an archetypal, yet challenging, ligand-to-protein interaction (**Figure 1D-G**). Implementation in an automated manner through software (see **Data Availability**) governing the uSTA workflow reduced artefacts arising from subjective, manual analyses (**Supplementary Figure S6**). The binding of L-tryptophan (Trp) to bovine serum albumin (BSA, **Figure 1D**) is a long-standing benchmark(*24*) due to both the plasticity of this interaction based on supposed hydrophobicity but also due to a lack of corresponding fully-determined, unambiguous 3D (e.g. crystal) structures. This is also a simpler amino acid•protein interaction system (non-modified protein, small ligand) that classical NMR/STD methods are perceived(*23, 31*) to have already delineated well.

As for a standard STD experiment, 1D ^1^H-NMR spectra were determined for both ligand and protein. In addition, mixed spectra containing both protein and excess ligand (P + L) were determined with and without excitation irradiation at frequencies corresponding to prominent resonance within the protein but far from any ligand (pulse ‘on’) or where the centre of the pulse was moved to avoid ligand and protein (pulse ‘off’, labeled ‘1D in Figures). Deconvolved spectra for ligand determined in the presence of protein were matched with high accuracy by uSTA (**Figure 1E**). Moreover, uSTA generated highly consistent binding ‘heat maps’, comprising atom-specific magnetization transfer efficiencies (proton data mapped onto heteroatoms by taking a local 1/r^6^ average to enable visual comparison) that described the pose of ligand bound to protein (**Figure 1F** see also **Supplementary Figures S8**,**S11**). These were determined over a range of ligand concentrations even as low as 40 μM (**Figure 1E(ii),1F(iii)**) where the ability of uSTA to extract accurate signal proved unprecedented and critical to quantitation of binding (see below). Binding maps were strikingly consistent across concentrations, indicating a single, consistent pose driven by strongest interaction of protein with the heteroaromatic indole side-chain of Trp. This not only proved consistent with X-ray crystal structures of BSA with other hydrophobic ligands,(*32*) it also revealed quantitative subtleties of this interaction at high precision – protein ‘grip’ is felt more at the centre of the indole moiety (across bond C4a–C7a) than at its periphery.

Next, with indications of expanded capability of uSTA in a benchmark system, we moved to first analyses of sugar•protein interactions. Sugar ligand trehalose (Tre) binds only weakly to trehalose repressor protein TreR and so proves challenging in ligand-to-protein interaction analysis.(*33*) Nonetheless, the uSTA workflow again successfully and rapidly determined atom-specific transfer efficiencies with high precision and resolution (**Supplementary Figure S3F**).

Atom-precise subtleties were revealed in this case as well: hotspots of binding occur around OH-3 / OH-4 graduate to reduced binding around both sugar rings with only minimal binding of the primary OH-6 hydroxyl (**Supplementary Figure S3F**). Once more, this uSTA-mapped P+L interaction proved consistent with prior X-ray crystal structures (*34*).

### Direct Determination of Ligand•Protein K_D_ using uSTA

The precision of signal determination in uSTA critically allowed variation of ligand/protein concentrations even down to low levels (see above); this valuably enabled direct determination of binding constants in a manner not possible by classical methods. Following measurement of magnetization transfer between ligand and protein, variation with concentration (**Figure 1E**) was quantitatively analyzed using modified Bloch-McConnell equations,(*35*) (see **Methods**). These accounted for intrinsic relaxation, cross-relaxation and protein-ligand binding (**Figure 1A**) to directly provide measurements of equilibrium binding *K*_D_ and associated kinetics (k_ex_). In the

Trp•BSA system this readily revealed a *K*_D_ = 38 ± 15 μM and k_on_ = 160000 ± 60000 M^-1^s^-1^, k_off_ = 6.0± 2.0 s^-1^ (**Figure 1G**), consistent with prior determinations by other solution-phase methods (*K*_D_ = 40 μM by isothermal calorimetry(*31*)). It should be noted that this direct method proved only possible due to the ability of the uSTA method to deconvolute a true signal with sufficient precision, even at the lower concentrations used and consequently lower signal (**Figure 1E**). Thus, uSTA enabled atom-mapping and quantitation for ligand binding improved over previous methods. Critically, these values were fully consistent with all observed NMR data and independently-obtained measures of *K*_D_.

### uSTA Allows Interrogation of Designed Crypticity in Influenza HA Virus Attachment

Having validated the uSTA methodology, we next used it to interrogate sugar binding by viral attachment protein systems that have proved typically intractable to classical methods. The haemagglutinin (HA) trimer of influenza A virus is known to be essential for its exploitation of sialoside-binding;(*36*) H1N1 has emerged as one of the most threatening variants in recent years. We took the H1 HA in both native form and in a form where the introduction of a sequon for adding further *N*-linked glycosylation in the sialoside binding site has previously been designed(*37*) to block sugar interaction *in trans* by creating a competing interaction *in cis*. This designed crypticity in a so-called HA-ΔRBS variant,(*37*) which creates an additional, potentially confounding, glycosylation background, provided another test of uSTA’s ability to delineate relevant sialoside ligand interactions in another important pathogen protein (**Figure 2A**). Notably, despite this intended *in cis* ‘interference’/inhibition, such was the precision of uSTA that residual *in trans* binding of sialo-trisaccharide **2** could still be detected in HA-ΔRBS, albeit at a lower, modulated level (as expected by design(*37*)). In this way, the sensitivity of uSTA to detect cryptic sialoside binding was confirmed.

**Figure 2.**
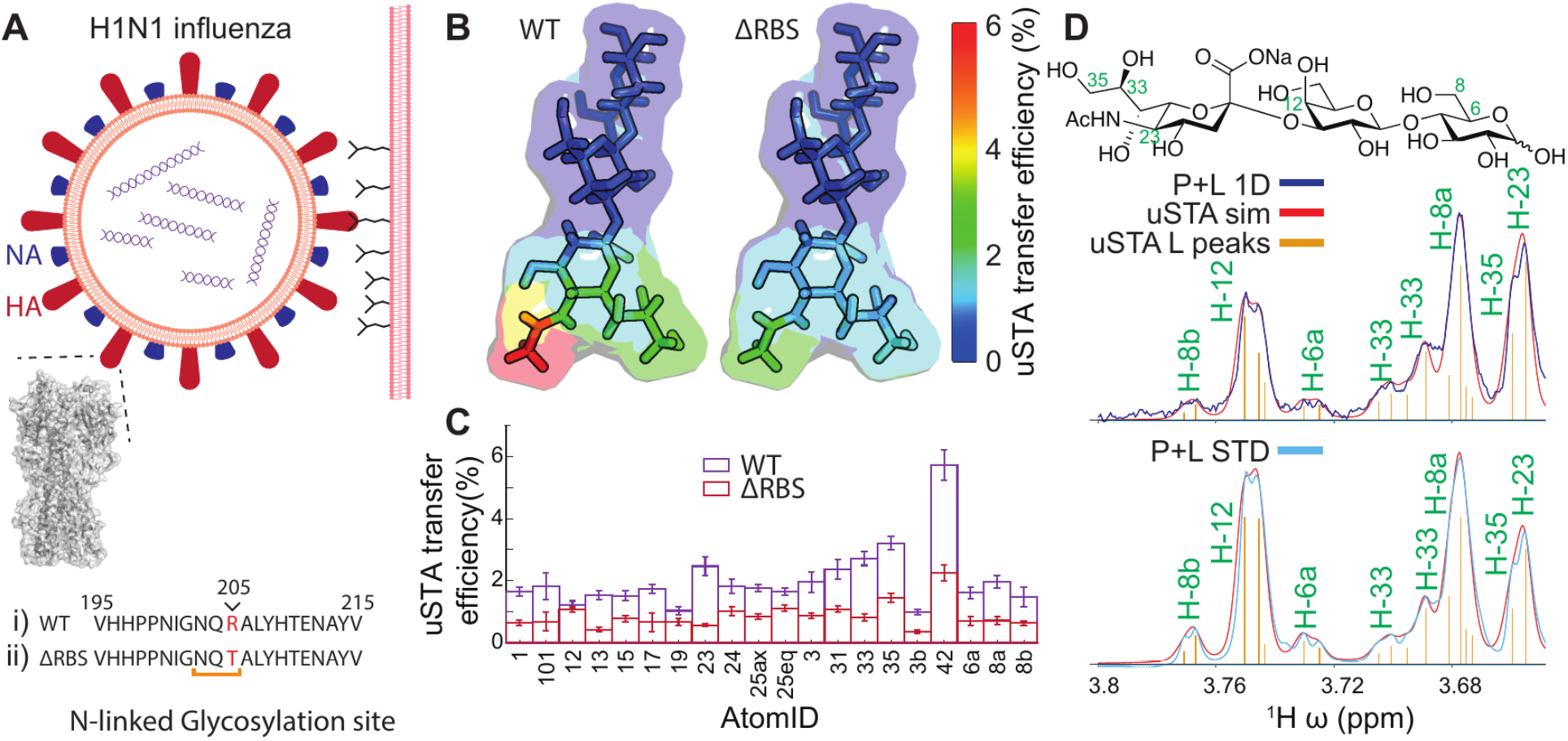
uSTA Allows Mapping of a Designed Cryptic Sugar-Binding Site in H1N1 Influenza Haemagglutinin (HA). **A):** Haemagglutinin (HA) presents on the surface of the viral membrane and has been shown to bind with sialic acid surface glycans to mediate host-cell entry. The validated(*37*) ΔRBS variant is created by the creation of an *N*-linked glycosylation site via the introduction of sequon NQT from wild-type NQR by the R205T point mutation in HA adjacent to the sialic-acid binding site. In this way, disruption of sialic acid binding through designed crypticity in HA-ΔRBS is intended to ablate the binding of HA-wild-type to sialosides. **B),C):** Notably, wild-type and ΔRBS variants of H1N1 haemagglutinin in fact show a remarkably similar overall binding *pattern* (**B**) for 2,3-sialo-trisaccharide **2** but a significant intensity moderation (**C**) for the ΔRBS variant, indicative of a partial (but not complete) loss of binding consistent with design.(*37*) **D):** Raw spectral data demonstrates atom-specific differences in intensity in the 1D vs the difference spectra can be discerned using uSTA. Note that atom numbering shown and used here is that generated automatically by uSTA.

### uSTA Reveals Natural, Cryptic, Sialoside Binding by SARS-CoV-2 Spike

Having validated uSTA in the characterization of a known, designed, intentionally-cryptic sialoside binding site in the attachment protein of one important pathogen virus, we next probed putative naturally-cryptic sialoside binding sites in SARS-CoV-2. Our analysis of the 1D protein ^1^H- NMR spectrum of the purified prefusion-stabilised ectodomain construct(*10*) of intact trimeric SARS-CoV-2-spike attachment protein (**Supplementary Figure S7**) revealed extensive protein glycosylation with sufficient mobility to generate a strong ^1^H-NMR resonance in the region 3.4-4.0 ppm (**Figure 3A**). Whilst lacking detail, these resonances displayed chemical shifts consistent with the described mixed patterns of oligomannose, hybrid and complex *N*-glycosylation found on SARS-CoV-2-spike after expression in human cells.(*38*) As such, these mobile glycans on SARS-CoV-2-spike contain sialoside glycan residues that not only confound analyses by classical NMR methods but they are also potential competing ‘internal’ (*in cis*) ligands for any putative attachment (*in trans*) interactions. Therefore, their presence in the protein NMR analysis presented clear confounding issues for typical classical STD analyses. As such, SARS-CoV-2-spike represented a stringent and important test of the uSTA method.

**Figure 3.**
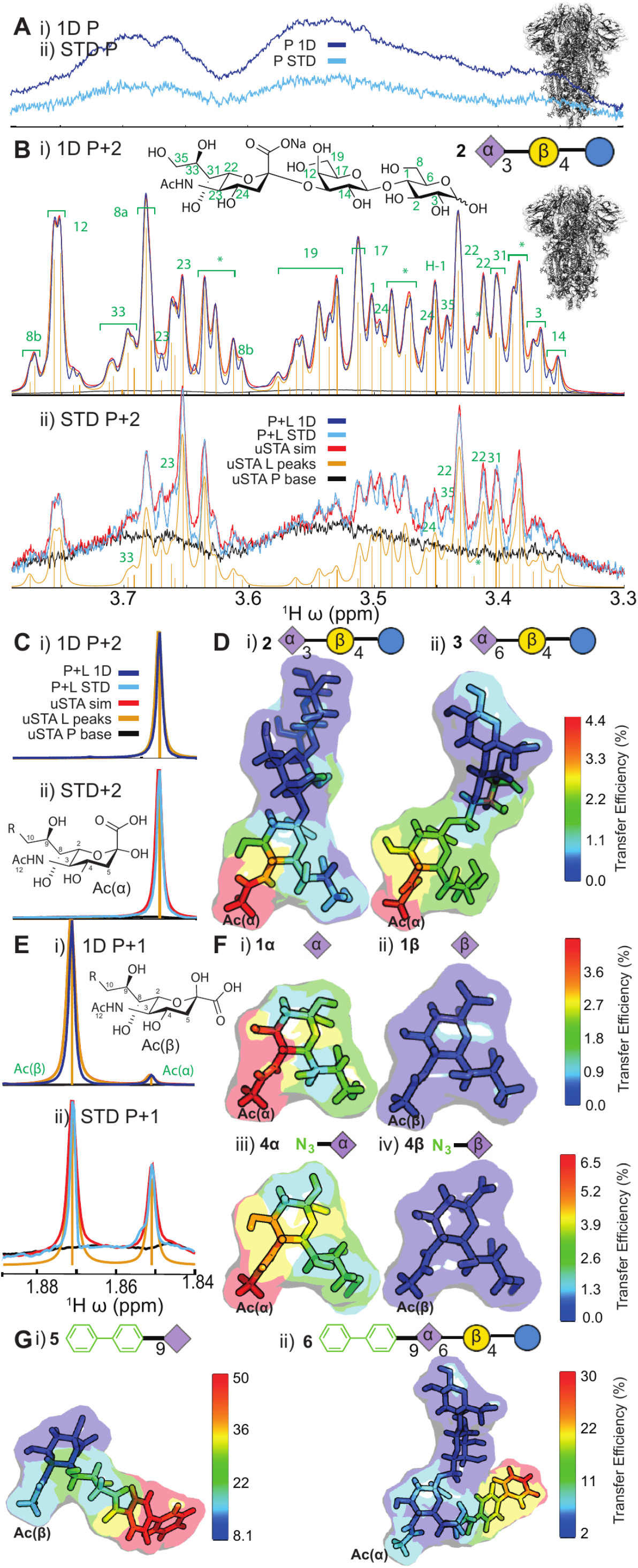
uSTA Reveals Interaction of Sialosides with SARS-CoV-2-spike Protein. A panel of natural, unnatural and hybrid variant sialoside sugars **1**-**6** (see **Supplementary** Figure 12) was used to probe interaction between sialosides and spike. **A**): The 1D ^1^H-NMR of SARS-CoV-2-spike protein shows considerable signal in the glycan- associated region despite protein size, indicative of mobile internal glycans in spike protein. This effectively masks traditional analyses, as without careful subtraction of the protein’s contributions to the spectrum (**Supplementary Figure S8**), the ligand cannot be effectively studied. **B**): Application of the uSTA workflow (**Supplementary Figure S6**) to SARS-CoV-2-spike protein (shown in detail for **2**) The uSTA process of: ligand peak assignment and deconvolution → P + L peak assignment and deconvolution → application to P + L STD yields precise atom-specific transfer efficiencies (**Supplementary Figure S6**). Note how in **ii)** individual multiplet components, have been assigned (**yellow**); the back-calculated deconvolved spectrum (**red**) is an extremely close match for the raw data (**purple**). The spectrum is a complex superposition of the ligand spectrum (and protein only yet uSTA again accurately deconvolves the spectrum revealing the contribution of protein only (**black**) and the ligand peaks (**yellow**). Using these data, uSTA analysis of the STD spectrum pinpoints ligand peaks and signal intensities. Note that spectral atom numbering shown and used here is that generated automatically by uSTA; all other numbering in sugars follows carbohydrate nomenclature convention. **C**): Application of the uSTA workflow (**Supplementary Figure S6**) reveals atom-specific binding modes to spike protein for both natural e.g. sialoside trisaccharides Siaα2,3Galβ1,4Glc (**2**) and Siaα2,6Galβ1,4Glc (**3**). **E,F**): In the spectra of sialic acid (**1**) and 9-N_3_ azido variant **4**, both interconverting α and β anomeric forms could be readily identified. Despite the dominance of the β form (94 %, **E(i)**), application of the uSTA method following assignment of resonances from the two forms allowed determination of binding surfaces simultaneously (**E(ii),F(i,ii)**). Spike shows strong binding preference for the α anomers (**F(i,iii) vs F(ii,iv)**) despite this strong population difference. Binding surfaces were also highly similar to those of extended trisaccharides **2** and **3**. **G**): Using these intensities, atom-specific transfer efficiencies can be determined with high precision, shown here for hybrid sialoside **5**. The details of both the unnatural BPC moiety and the natural sialic-acid moiety can be mapped: although the unnatural aromatic BPC dominates interaction, uSTA nonetheless delineates the subtleties of the associated contributions from the natural sugar moiety in this ligand (see also **Supplementary Figure S5,S6**).

We evaluated a representative panel of both natural and site-specifically-modified unnatural sialosides as possible ligands of spike using uSTA (**Figure 3** and **Supplementary Figure S3,S12**). Strikingly, whilst use of classical methods provided an ambiguous assessment (**Supplementary Figure S8**), use of uSTA immediately revealed binding and non-binding sugar ligands (**Figure 3** and **Supplementary Figures S8**,**S10,S11** and **Supplementary Table S7**). Initially, the simplest sialoside, *N*-acetyl-neuraminic acid (**1**), was tested as a mixture of its mutarotating anomers (**1α ⇔ 1β** ). When analyzed using uSTA, these revealed (**Figure 3E,3F(i,ii)** and **Supplementary Table S7** and **Supplementary Figure S11**) clear ‘end on’ interactions by **1α** as a ligand (**Figure 3F(i)**), mediated primarily by NHAc-5, but no reliably measurable interactions by **1β** (**Figure 3F(ii)**). It should be noted that this detection of selective α-anomer interaction, despite the much greater dominance of the β-anomer in solution, provided yet another further demonstration of the power of the uSTA method, here operating in the background of dominant alternative sugar (**Figure 3E** and **Supplementary Figure S11**). Together these data not only confirmed interaction of SARS-CoV-2- spike with the smallest natural sialoside sugar but also revealed striking ligand stereochemical (α over β) selectivity by the protein. Provocatively, the α selectivity correlates with the near-exclusive occurrence of sialosides on host cell surfaces as their α- but not β-linked conjugates (see also below).

Next, having confirmed simple, selective monosaccharide α-sialoside binding we explored extended, α-sialoside oligosaccharide ligands (compounds **2**, **3**, **Figure 3C,D**) that would give further insight into binding of natural endogenous human cell-surface sugars, as well as unnatural variants (compounds **4-6**, **Figure 3F,G**) that could potentially interrupt such binding. Sialosides are often found appended to galactosyl (Gal/GalNAc) residues in either α2,3- (**2**) linked or α2,6- linked form (**3**); both were tested (**Figure 3C,D**). Both also exhibited ‘end on’ binding consistent with that seen for *N*-acetyl-neuraminic acid (**1**) alone, but with more extended binding surfaces (**Figure 3D**); qualitatively this suggested a stronger binding affinity (see below for quantitative analysis).

Common features of all sialoside binding modes were observed: the NHAc-5 acetamide of the terminal sialic acid (Sia) is a binding hotspot in **1**, **2** and **3** that drives the ‘end on’ binding.

Differences were also observed: the α2,6-trisaccharide (**3**) displayed a more extended binding face yet with less intense binding hotspots (**Figure 3D**) engaging additionally the side-chain glycerol moiety (C7-C9) of the terminal Sia acid as well as the OH-4 C4 hydroxyl of the galactosyl (Gal) residue. The interaction with α2,3-trisaccharide (**2**) was tighter and more specific to NHAc-5 of the Sia.

These interactions of the glycerol C7-C9 side-chain detected by uSTA were probed further through construction (**Supplementary Figure 12**) of unnatural modified variants (**4**-**6**, **Figure 3F,G**). Whilst replacement of the OH-9 hydroxyl group of sialic acid Sia with azide N_3_-9 in **4** (**Figure 3F(iii)** and **Supplementary Table S7**) was well-tolerated, larger changes (replacement with aromatic group biphenyl-carboxamide BPC-9) in **5** and **6** led to an apparently abrupt shift in binding mode that was instead dominated by the unnatural hydrophobic aromatic modification (**Figure 3G, 4C**). As for native sugar **1**, azide-modified sugar **4** also interacted with spike in a stereochemically-specific manner with only the α-anomer displaying interaction (**Figure 3F(iii)**), despite dominance of the β-anomer in solution (**Figure 3F(iv)**. Notably therefore, uSTA allowed precise dissection of interaction contributions in these unusual hybrid (natural-unnatural) sugar ligands that could not have been determined using classical methods (see **Supplementary Note 5**).

**Figure 4.**
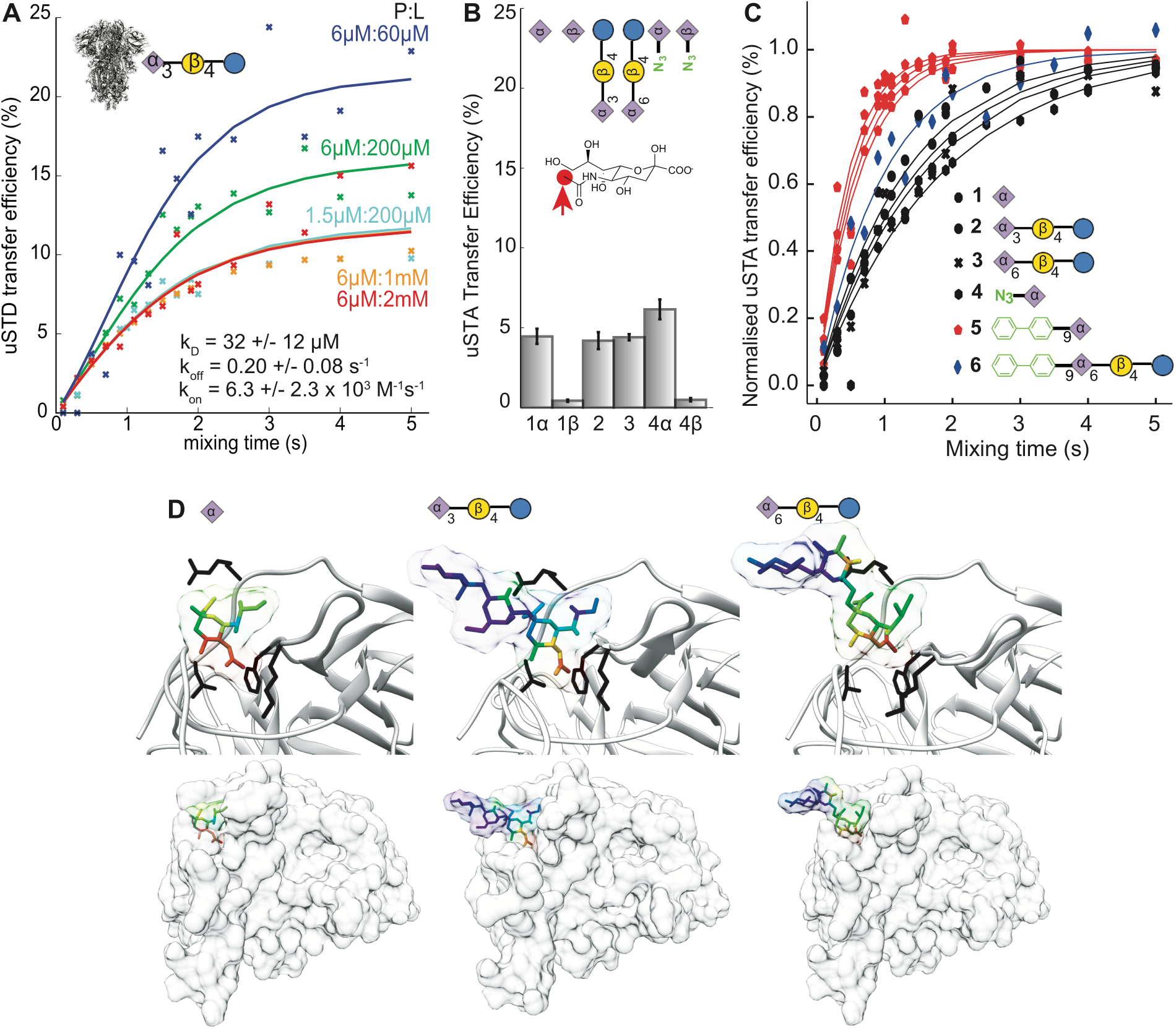
Quantitative uSTA Analyses Allow Comparison and Predictive Constraints for Protein Ligand-Binding Prediction. **A**): Quantitative analysis of the STD build up curves using a modified set of Bloch-McConnell equations that account for binding and cross relaxation allow us to determine thermodynamic and kinetic parameters that describe the SARS-CoV-2•**2** interaction, *K*_D_, k_on_, and k_off_. The values obtained are indicated. Errors come from a boot-strapping procedure (see **Methods**). **B**): Normalized uSTA transfer efficiencies of the NAc-5 methyl protons can be determined for each ligand studied here. This allowed relative contributions to ‘end on’ binding to be assessed via uSTA in a ‘mode-specific’ manner. This confirmed strong alpha over beta sialoside selectivity. Errors were determined through a boot-strapping procedure where mixing times were sampled with replacement, allowing for the construction of histogram of values in the various parameters that robustly reflect their fitting errors. **C**): Normalised build-up curves for the most intense resonances allowed two distinct modes of binding to be identified in natural (**1-4**) and hybrid (**5**, **6**) sugars. Data here is shown at constant protein and ligand concentrations. With the BPC moiety present, the build-up of magnetization occurs significantly faster than when not; various such hybrid ligands give highly similar curves. By contrast, natural ligands have a much slower build-up of magnetization. This, together with the absolute transfer efficiencies being very different, and the overall pattern on the interaction map combine to reveal that the ligands are most likely binding via two different modes and possibly locations on the protein. **D):** Coupling uSTA with an integrative modelling approach such as HADDOCK (High Ambiguity Driven DOCKing)(*39, 40*) allowed generation and, by quantitative scoring against the experimental uSTA data, selection of models that provide atomistic insights into the binding of sugars to the SARS-CoV-2-spike protein, as shown here by superposition of uSTA binding ‘map’ onto modelled poses. uSTA mapping the interaction between SARS-CoV-2-spike (based on RCSB 7c2l (*18*)) with ligands **1a**, **2** and **3** identifies the NHAc-5 methyl group of the tip sialic acid residue making the strongest interaction with the protein. By filtering HADDOCK models against this information, we obtain structural models that that describe the interaction between ligand and protein (**Supplementary Figure S14**). Most strikingly, we see the same pattern of interactions between protein and sialic acid moiety in each case, where the NAc methyl pocket is described by a pocket in the spike NTD. Although sequence and structural homology are low (**Supplementary Figure S1**), MERS-spike protein possesses a corresponding NHAc-binding pocket characterized by an aromatic (F39)–hydrogen-bonding (D36)–hydrophobic (I132) triad.(*5*)

Next, using variable concentrations of the most potent natural ligand α2,3-trisaccharide **2** [6μM spike, **2** at 60μM, 200μM, 1mM and 2mM excitation at 5.3 ppm] and variable concentrations of spike protein we used the uSTA method to directly determine solution-phase affinities (**Figure 4A**): *K*_D_ = 32 ± 12 μM with k_on_ = 6300 ± 2300 M^-1^s^-1^, k_off_ = 0.20 ± 0.08 s^-1^. We also probed binding in a different mode by measuring affinity of spike to **2** when displayed on a modified surface (**Supplementary Figure S13**) using surface plasmon resonance (SPR) analysis; the latter generated a corresponding *K*_D_ = 23 ± 6 μM (k_on_ = 1030 M^-1^s^-1^). Interestingly, these similar determined values for sialoside ligand in solution (by uSTA) or when displayed at a solid-solution interface (by SPR) suggested no significant avidity gain from multiple sugar display on a surface.

### Structural Insights from uSTA Delineate Binding to SARS-CoV-2 Spike

uSTA analyses consistently identified binding hotspots in sugars **1**-**4** providing the highest transfer efficiencies in an atom-specific manner – most notably the ‘end’ NHAc-5 acetamide methyl group of the tip sialic acid residue in all. Combination of uSTA with so-called High Ambiguity Driven DOCKing (HADDOCK) methods(*39, 40*) was then used to probe likely regions in SARS-CoV-2- spike for this ‘end-on’ binding mode via uSTA-data-driven atomistic models. In each case, a cluster of likely poses emerged (**Figure 4D**) for **1α**, **2** , **3** (see **Supplementary Figure S14** for details) consistent with ‘end-on’ binding where the acetamide NHAc-5 methyl group of the sialic acid moiety was notably held by the unusual beta-sheet rich region of the NTD of SARS-CoV-2-spike.

Under the constraints of uSTA and homology, a glycan-binding pocket was suggested, delineated by a triad of residues (F79, T259 and L18) mediating aromatic, carbonyl-hydrogen-bonding and hydrophobic interactions, respectively. However, it should be noted, the sequence and structural homology to prior (i.e. MERS) coronavirus spike proteins in this predicted region was low; the MERS-spike protein uses a corresponding NHAc-binding pocket characterized by an aromatic (F39), hydrogen-bonding (D36) and hydrophobic (I132) triad to bind the modified sugar 9-*O*-acetyl-sialic acid.(*5*)

SARS-CoV-2 glycan attachment mechanisms have to date only identified a role for spike RBD in binding rather than NTD (*15, 16*). First, we used uSTA to compare the relative potency of the sialoside binding identified here to previously identified(*16*) binding of heparin motifs (**Figure 5A,B**). Heparin sugars **7** and **8** of similar size to natural sialosides **3** and **4** were selected so as to allow a near ligand-for-ligand comparison based on similar potential binding surface areas. **7** and **8** also differed from each other only at a single glycan residue (residue 2) site to allow possible dissection of subtle contributions to binding. Notably, unlike the ‘end-on’ binding seen for sialosides **3** and **4**, uSTA revealed an extended, non-localized binding interface for **7** and **8** consistent instead with ‘side-on’ binding (**Figure 5Aii,Bii**).

**Figure 5.**
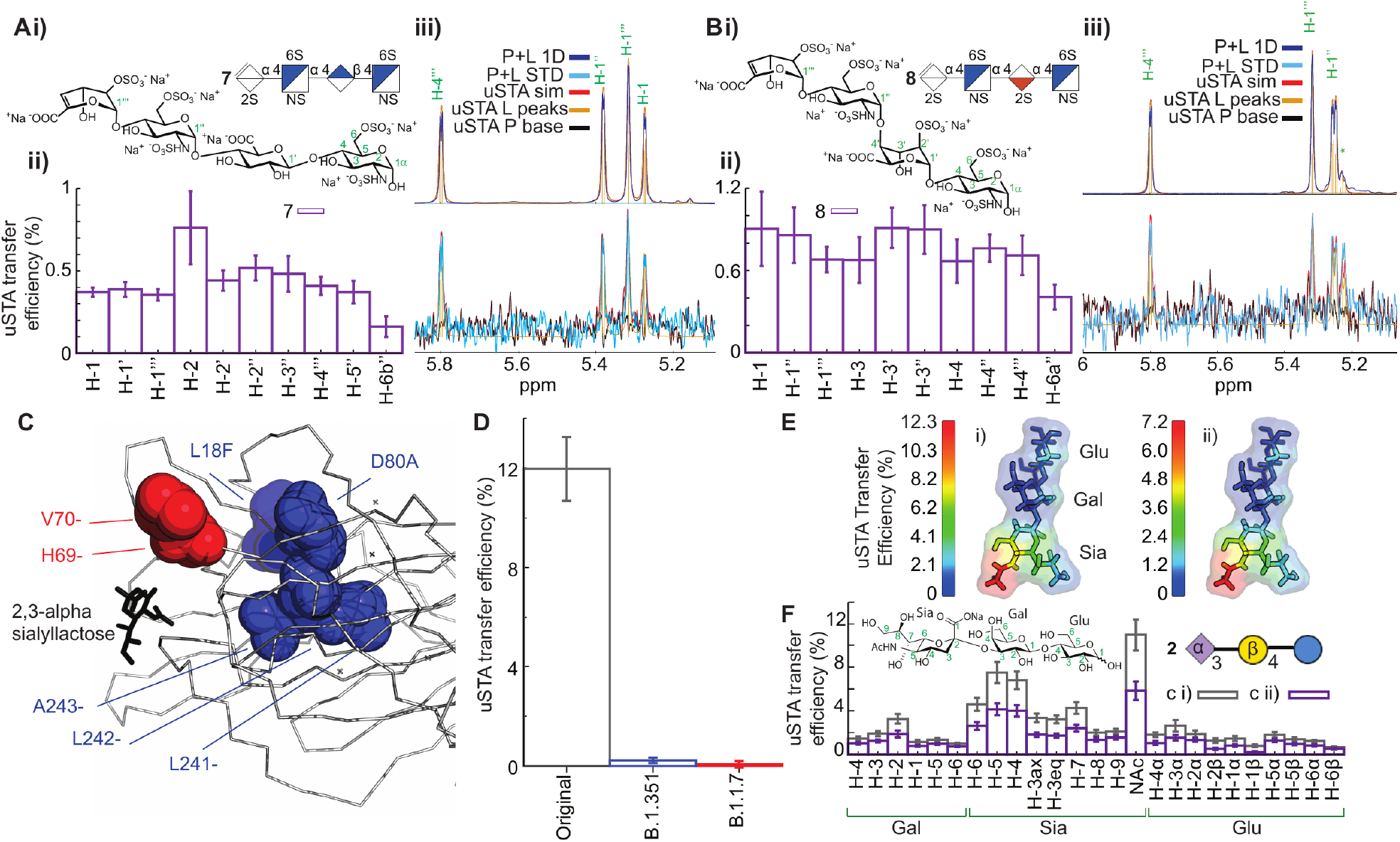
Comparison of SARS-CoV-2 Glycan Attachment Mechanisms and Variant Evolution via uSTA Suggests Binding Away from the RBD that is Lost. **A), B):** Two heparin tetrasaccharides (**7**, **8**) are shown by uSTA (**Aiii,Biii**) to bind B-origin-lineage SARS-CoV-2 spike protein in a ‘side-on’ mode (**Aii,Bii**). Atom specific binding is shown in **A)ii**, **B)ii**. Assignments shown in green utilize conventional glycan numbering. **C**): Mutations to NTD-region identified by uSTA are found in the B.1.351/beta (**blue**) and B.1.1.7/alpha (**red**) lineage variants of SARS-CoV-2-spike. **D**): Ablated binding of the alpha-2,3-sialo-trisaccharide **2**, as measured by the transfer efficiency of the NHAc protons is identified by uSTA in the B.1.351/beta (blue) and B.1.1.7/alpha (red) lineage variants of SARS-CoV-2-spike. This proves consistent with mutations that appear in the sialoside-binding site of NTD identified in this study (see **C** and Figure 6). **E,F**): uSTA of sialoside **2** with B-origin-lineage-SARS-CoV-2 spike in the presence of the potent RBD-neutralizing nanobody C5 (spike (**E i**), nanobody-plus-spike (**E ii**)) shows essentially similar binding patterns with uniformly modulated atomic transfer efficiencies (**F**).

Next, we examined the possible evolution of sialoside binding over lineages of SARS-CoV- 2.(*41*) Two notable ‘variants of concern’ alpha/B1.1.7 (so-called ‘UK or Kent’) and beta/B1.351 (so-called ‘South African’) emerged in secondary phases of the pandemic. When corresponding alpha/B1.1.7 and beta/B.1.351 spike-protein variants were probed by uSTA, these displayed dramatically ablated binding towards sialoside **2** as compared to first phase B-origin-lineage spike (**Figure 5D** and **Supplementary Figure S15**).

Finally, to explore the possible role of sialoside binding in relation to ACE2 binding we also used uSTA to probe the effects upon binding of the addition of a known, potent neutralizing antibody of ACE2•spike binding, C5 (**Figure 5D,E** and **Supplementary Figure S16**).(*42*) Notably, assessment of binding to sialoside **2** in the presence and absence of antibody at a concentration sufficient to saturate the RBD led to only slight reduction in binding. Uniformly modulated atomic transfer efficiencies and near-identical binding maps (**Figure 5E,F**) were consistent with a maintained sialoside-binding pocket with undisrupted topology and mode of binding.

Together these data (‘end-on’ vs ‘side-on’, non-RBD-driven), suggested that the sialoside binding that we observe with uSTA, involves a previously unidentified ‘end-on’ mechanism/mode that operates in addition to and potentially cooperatively with ACE2-binding in SARS-CoV-2. It also suggested that the primary sialoside glycan-binding site SARS-CoV-2-spike is distinct from that of heparin (‘end-on’ vs ‘side-on’), not in the RBD (not neutralized by RBD-binding Ab) and found instead in an unusual NTD region that has become altered in emergent variants (loss of binding in alpha/beta variants of concern).

### Cryo-EM Pinpoints the Sialoside-binding Site in B-origin-lineage-Spike

Structural analysis of the possible binding of sialosides has been hampered to date by the moderate resolution, typically less than 3 Å, of most SARS-CoV-2-spike structures. Moreover, examination of current deposited cryo-EM-derived coulombic maps, show that those for the NTD of spike are often the least strong. In our initial attempts with native protein, large stretches of amino acids within the NTD were not experimentally located (*42*); the most disordered regions occur in the NTD regions that contribute to surface of the spike. Whilst a stabilized closed mutant form (*43*) of the spike was examined and gave improvements, the coulombic map was still too weak and noisy to permit tracing of many of the loops in the NTD. However, with a reported fatty-acid bound form of the spike,(*44*) which has shown prior improved definition of the NTD, we were able to collect a 2.3 Å data set in the presence of the α2,3-sialo-trisaccharide **9**. The map was very clear for almost the entire structure including the previously identified linoleic acid; only 13 N-terminal residues and two loops (residue 618-632 and 676-689) were not located. Consistent with all other structures of spike to date, the density is weaker at the outer surface of the NTD than the core of the structure. However, the map was not only of sufficient quality to model N-glycosylation at site N149, which is in a flexible region, but also even the fucosylation state of *N*-linked glycans at N165 (**Figure 6A**).

**Figure 6:**
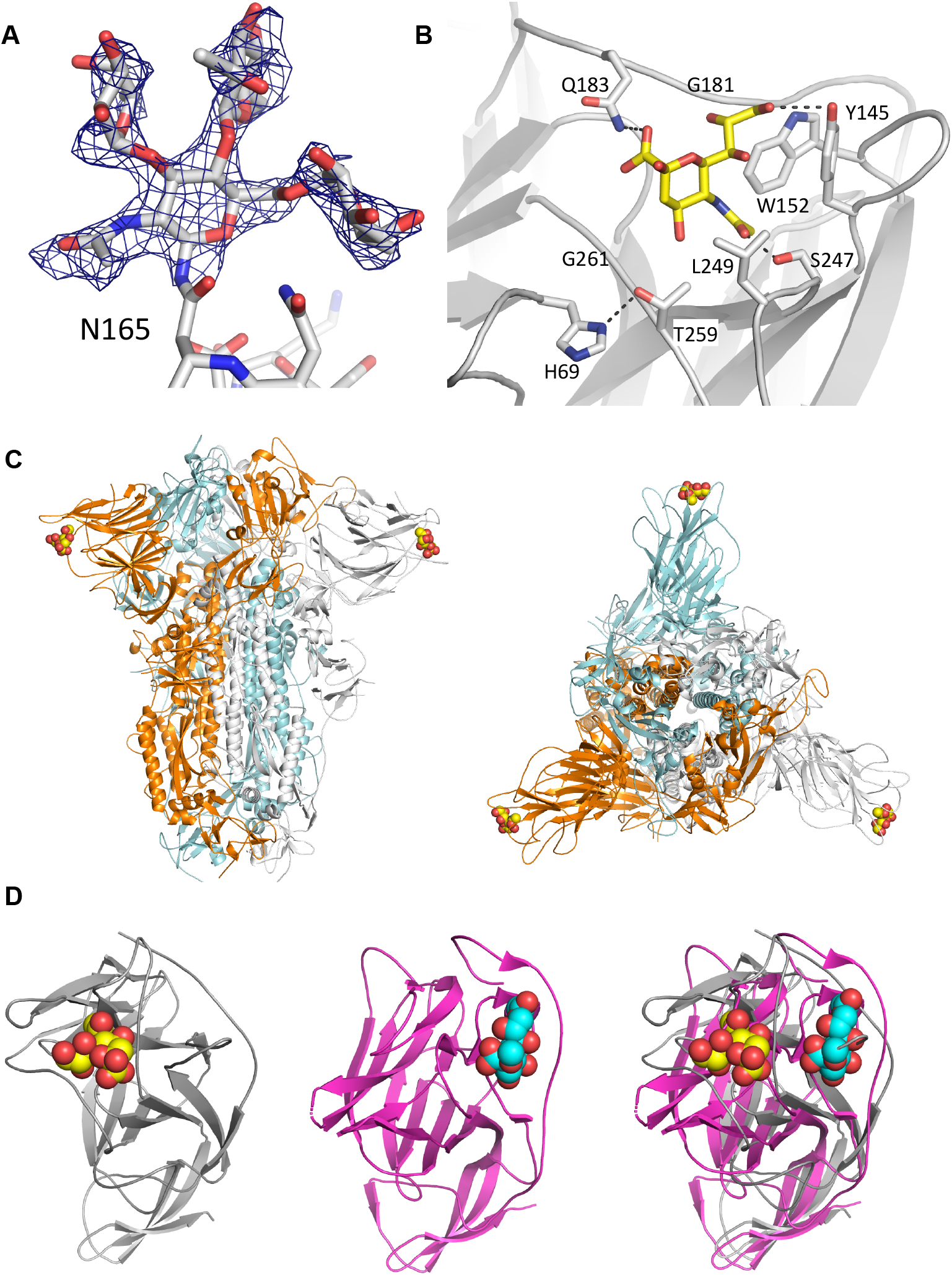
Cryo-EM Analysis of Sialoside Binding in Wuhan-Hu-1-SARS-CoV-2-spike. **A):** In the presence of sialoside **9** (**Supplementary Figure S19**) a well-ordered structure with resolution (2.3 Å) sufficient to identify even glycosylation states of N-linked glycans within spike was obtained. **B):** The sialoside-binding site in the NTD is bounded by H69, Y145, W152, S247 and Q183. Q183 controls stereochemical recognition of α-sialosides by engaging COOH-1. Y145 engages the C7- C9 glycerol sidechain of sialoside. S247 engages the NHAc-5. Notably, Y145 and H69 are deleted in the alpha variant of SARS-CoV-2 that loses its ability to bind sialoside (see Figure 5). **C):** Spike’s sialoside binding site is found in a distinct region of the NTD (**left**, from side, **right** from above). **D):** Superposition (**right**) of SARS-CoV-2-NTD (**left**, grey) with MERS-NTD (**middle**, magenta) shows distinct sites 12 Å apart.

Contouring the map to fit the structure of the protein, we observed density in a pocket at the surface of the NTD lined by residues H69, Y145, W152, Q183, L249 and T259. This density is absent in other spike structures of resolution higher than 2.7 Å resolution (RCSB 7jji, 7a4n, 7dwy, 6x29, 6zge, 6xlu, 7n8h, 6zb5 and 7lxy), even those (such as RCSB 7jji, 6zb5 and 6zge) which have a well-ordered NTD. The density when contoured at 2.1 σ is fitted by an α-sialoside consistent with the terminal residue of **9**, with the distinctive glycerol and N-acetyl groups clear (**Figure 6B** and **Supplementary Figure S17**). The sialoside was included in the refinement and the thermal factors (108 Å^2^) were comparable to those for the adjacent protein residues (95 - 108 Å^2^). In this position, the glycerol moiety makes a hydrogen bond with the side chain of Y145, the N-acetyl group with S247 and the carboxylate with Q183. In addition there are hydrophobic interactions with W152. Lowering the map threshold to 1.6 σ would be consistent with a second pyranoside (e.g. galactoside) residue. At this contour level the map clearly covers the axially configured carboxylate of the sialic acid (**Supplementary Figure S17**). The middle galactoside residue of **9** positioned in this density would make contacts with R248 and L249.

Structural superposition of the NTD with that of the MERS spike (RCSB 6NZK) shows that whilst the sialic acid-binding pockets of both are on the outer surface, these pockets are 12 Å apart (as judged by the C2 atom of respective sialic acids, **Figure 6D**). In MERS spike sialic acid is bound at the edge of the central β-sheet, whereas in SARS-CoV-2 the sugar is bound at the centre of the sheet; thus the pockets use different elements of secondary structure. Due to distinct changes in the structure of the loops connecting the strands, the sialic acid pocket from one protein is not present in the other protein.

We noted several regions of additional density that were not fitted by model. Density consistent with a lipid-like or polyol molecule was visible in the hydrophobic cleft in the NTD formed by L126, Y170, F192, L226 and V227. This location was found to bind polysorbate in a previous report (RCSB 7jji), however the density observed here is not consistent with polysorbate or sialoside **9**. Remote from the NTD there is aromatic ring-like density sandwiched between the side chains of Y660 and Q675 that is absent in well-ordered polysorbate spike complex (**Supplementary Figure S18**). This density is consistent with the binding of the tri-iodo benzamide ring that is attached to the reducing terminus of sialoside **9**. The density is not found in other spike structures we examined (RCSB 7a4n, 7dwy, 6x29, 6zge, 6xlu, 7n8h, 6zb5 and 7lxy). Finally, when spike protein was prepared identically in the absence of ligand **9** we were able to collect a 2.47 Å data set. The resulting map was well ordered, sufficient again to show clearly both fucosylation state at N165 and N-linked glycosylation at N17. Very similar density in the cleft that was previously reported to bind polysorbate was again observed. However, no density for either the sialioside nor iodobenzene moieties of **9** were observed, further confirming our assignments.

### Identification of Sialoside Trisaccharide as a Ligand for B-origin-lineage SARS-CoV-2 Correlates with Clinical Genetic Variation in Early-phase Pandemic

The identification here of a novel sialoside binding mode by spike confirms a potential attachment point for SARS-CoV-2 found commonly on cell surfaces (sialosides are attached both as glycolipid and glycoprotein glyco-conjugates). This, in turn, raised the question of whether glycosylation function in humans impacts upon infection by SARS-CoV-2 and hence in the presentation and pathology of COVID19 disease. Analysis of whole exome sequencing data of an early 2020 cohort of 533 COVID-19-positive patients (see **Supplementary Table S1**) identified two glycan-associated genes within the top five that were most influential upon disease severity.

Specifically, recursive feature elimination applied to a Least Absolute Shrinkage and Selection Operator (LASSO)-based(*45*) logistic regression model identified: *LGALS3BP* (4^th^ out of >18000 analyzed genes) and *B3GNT8* (5^th^ out of >18000) (**Figure 7A** and **Supplementary Figure S19**). Variants in these two genes were beneficially associated with less severe disease outcome (**Figure 7B,C**, see also **Supplementary Tables S2-6** for specific *B3GNT8* and *LGALS3BP* genetic variants, *B3GNT8* chi-squared five categories, *B3GNT8* chi-squared 2x2, *LGALS3BP* chi-squared five categories, *LGALS3BP* chi-squared 2x2, respectively).

**Figure 7.**
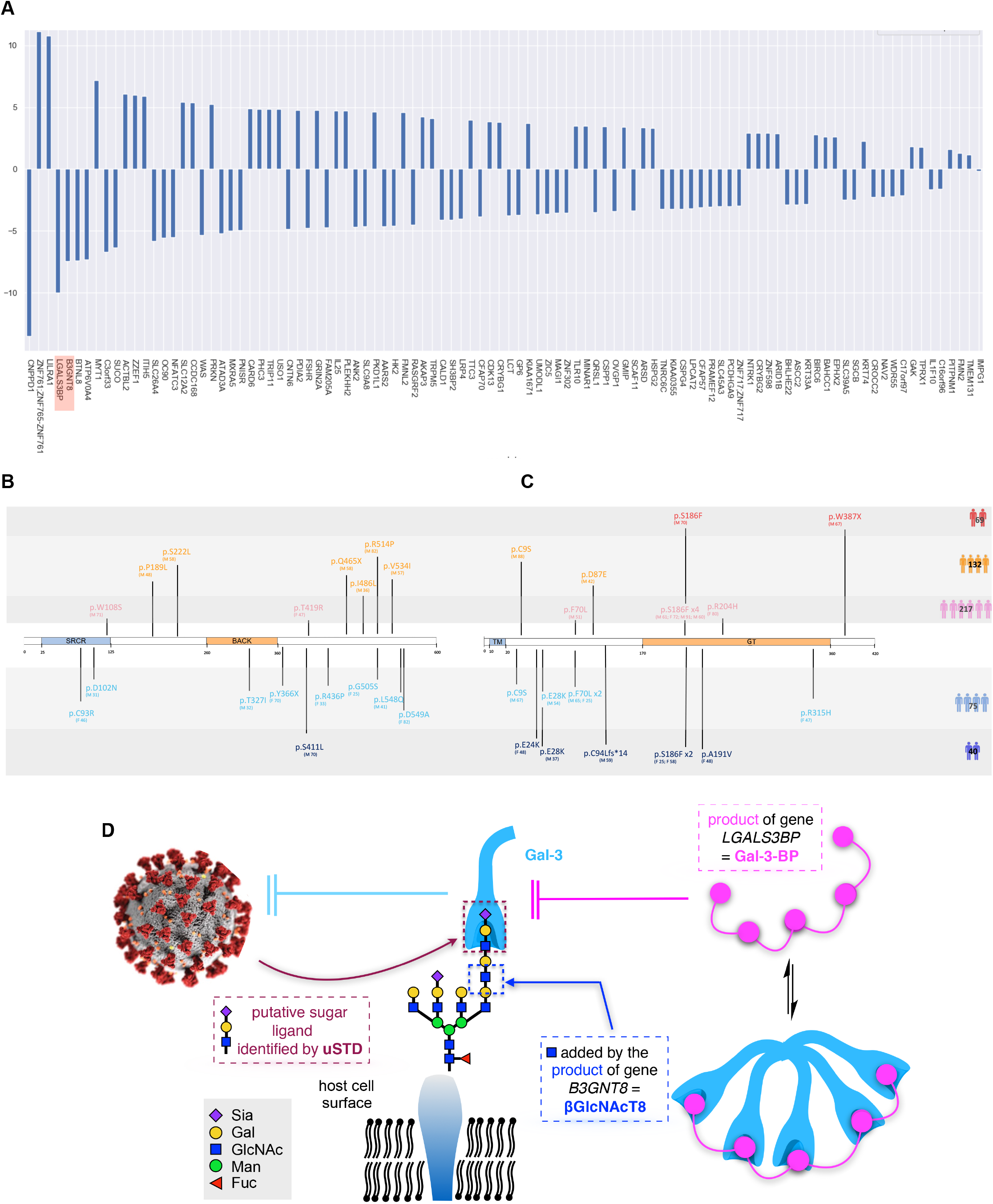
Analyses of Early 2020 SARS-CoV-2-positive Patients Reveals Glycan-Associated Genes Suggesting a Model of Glycan Interaction Consistent with uSTA Observations of Sialoside Binding in B-origin-lineage SARS-CoV-2. **A**): Histogram of the LASSO-based Logistic Regression weightings after Recursive Feature Elimination analysis of 533 SARS-CoV-2-positive patients. Positive weights score susceptible response of gene variance to COVID19 disease, whereas negative weights confer protective action through variance. Variation in glycan-associated genes *B3GNT8* and *LGALS3BP* score 2^nd^ and 3^rd^ out of all (>18000) genes as the most protective, respectively (**highlighted red**). **B**),**C**): Distribution of rare variants in *B3GNT8* and *LGALS3BP*. **B**) Rare beneficial mutations distributed along the Gal-3-BP protein product of *LGALS3BP*, divided into the Scavenger Receptor Cysteine-Rich (SRCR)-domain (**light blue**) and the BACK domain (**light orange**). **C)** Rare beneficial mutations distribution along the βGlcNAcT8 protein product of *B3GNT8* divided into the predicted transmembrane (TM) domain (**light blue**) and glycosyltransferase catalytic (GT) domain (**light orange**) which catalyzes the transfer of polyLacNAc-initiating GlcNAc onto tetraantennary *N*- linked glycoproteins (see also **D**). The different colours of the mutation bands (top to bottom) refer to the severity grading of the COVID19-postive patients who carried that specific mutation (**red**: Hospitalized intubated; **orange**: Hospitalized CPAP/BiPAP; **pink**: Hospitalized Oxygen Support; **light blue**: Hospitalized w/o Oxygen Support; **blue**: Not hospitalized a/paucisymptomatic). **D**): A proposed coherent model consistent with observation of implicated *B3GNT8* and *LGALS3BP* genes and the identification of sialoside as ligands for spike by uSTA and cryo-EM. Strikingly, although independently identified, *B3GNT8* and *LGALS3BP* produce gene products βGlcNAcT8 and Gal-3-BP, respectively, that manipulate and/or engage with processes associated with a common polyLacNAc-extended chain motif found on tetraantennary *N*-linked glycoproteins. A model emerges in which any associated loss-of-function from variance leads either to loss of polyLacNAc-extended chain (due to loss of initiation by βGlcNAcT8) or enhanced sequestration of by Gal-3 polyLacNAc-extended chain (which is antagonized by Gal-3-BP). Both, would potentially lead to reduced access of virus spike to uSTA-identified motifs.

*LGALS3BP* encodes for a secreted protein Galectin-3-binding-protein (Gal-3-BP, also known as Mac-2-BP) that is a partner and blocker of a specific member (Gal-3) of the galectin class of carbohydrate-binding proteins.(*46*) Galectins are soluble, typically secreted and implicated in a wide-range of cellular functions.(*47*) Notably Gal-3 binds the so-called poly-*N*-acetyl- lactosamine [polyLacNAc or (Gal-GlcNAc)_n_] chain-extension variants found in tetraantennary *N*- linked glycoproteins (**Figure 7D**), including those displaying sialyl-Gal-GlcNAc sialoside motifs.(*48, 49*) Variants in *LGALS3BP* were present in in 9/114 (∼8%) of a/pauci-symptomatic subjects or mildly-affected patients compared to 8/419 (< 2%) of the remaining 419 patients who required more intensive care: oxygen support, CPAP/BiPAP or intubation; none of the 69 most seriously affected patients (intubated) carried variants of *LGALS3BP* (**Figure 7B**). Intact *LGALS3BP*-gene- product Gal-3-BP therefore appears correlated with more severe COVID19 outcome. The other implicated gene, *B3GNT8* encodes for a protein glycosyltransferase β-1,3-*N*-acetyl- glucosaminyltransferase-8 (GlcNAcT8 or β3GnT8) that is responsible for the creation of the anchor-point of poly-*N*-acetyl-lactosamine (polyLacNAc) in such tetraantennary *N*-linked glycoproteins (**Figure 7D**).(*50*) Again, rare variants in *B3GNT8* were present in in 11/114 (∼10%) of a/pauci-symptomatic subjects or mildly-affected patients compared to 10/419 (∼2%) of the remaining 419 patients who required more intensive care (**Figure 7C**).

## Discussion

The recognition of sugars on cell-surface glycans lies at the core of human biology. The traditional, surface-display biophysical methods for analyzing a protein’s specificity for a particular glycan are well established (e.g. ‘glycochip’/glycan- array binding analyses). There are sometimes format dependencies that can give apparently contradictory results, as seen for the spike protein from SARS-CoV-2. For small molecules binding to large proteins, STD NMR is an attractive orthogonal measurement and it is central to so-called fragment based drug discovery (FBDD) (*51*). However, in glycosylated proteins (i.e. most derived from mammalian expression), such as spike, the power of NMR analysis of sugar-binding is very limited. This is because the resonances from the highly mobile, protein-linked glycans are visible (in contrast to the protein) and overlap with the resonances from the glycan ligand. In addition, textbooks for FBDD set out particular experimental constraints including the binding kinetics. We have shown that a rigorous evaluation of the physics underpinning magnetization transfer leads to a reformulation of the approach and when coupled with the application of a modern Bayesian approach to signal analysis produces a new NMR tool that we have named uSTA. uSTA has neither the limitations nor constraints of conventional STD.

As we show with well-behaved model systems, uSTA now provides a comprehensive treatment of saturation transfer that is a ready method for not only identifying protein ligands but also their quantitative binding parameters. Its workflow does not require the use of (e.g. relaxation) filters and is automated by freely-distributed software (see **Data Availability**). Using the influenza virus hemagglutinin HA protein, uSTA was able to unambiguously identify sialoside binding (even when rendered cryptic) demonstrating its ability to handle complex sugars such as sialosides and a large protein.

The contradictory results for sugar binding to the spike of SARS-CoV-2 are hindering the field. If the protein does bind sialic acid, it is an important finding that opens up investigations as to the potential role of such recognition. Equally, if the protein does not bind sialic acid, these questions do not arise and other questions are pursued. Experimentally, there are few, if any, good orthogonal approaches to the surface display methods. Having demonstrated the power of uSTA, we applied it to the problem of spike protein sugar recognition. uSTA clearly shows that spike of B- origin-lineage of SARS-CoV-2 binds sialic acid and this binding is more potent when sialic acid terminates galactosyl oligosaccharides. uSTA reveals that the NHAc-5 *N*-acetyl group at the sugar’s tip is buried, a mode of binding we refer to as ‘end on’. Using uSTA-derived constraints we modelled binding of the sugar – this showed that the NTD of spike was the most likely site. The NTD in the related MERS virus also binds sialic acid. Our modelling was partly misled by the lower resolution of initial structures of spike; in these the NTD is highly disordered. uSTA also identified a second, differing mode of binding in SARS-CoV-2-spike only by hybrid, aromatic sugars (e.g. **5**, **6**, **9**) driven by aromatic engagement (**Figure 3**).

Encouraged by the predictions of uSTA we determined the cryo-EM structure of the lipid stabilised spike protein soaked with a heavy-atom containing sialotrisaccharide **9**. The resulting very well-ordered high resolution structure shows sialic acid bound in a pocket that specifically engages the glycerol and NHAc-5 *N*-acetyl groups of the sugar, fully validating the uSTA analysis. The pocket is located at the very edge of the NTD and is formed by loops from the central β-sheet. These structural data rationalise the observed (via uSTA) structure-activity relationships for sialic acid derivatives used as probes. The loops in the beta-strand rich NTD of SARS-CoV-2-spike, which can now be seen in a well-ordered 3D structure, are structurally distinct from MERS. As a result, although both proteins bind sialic acid they bind at adjacent but different locations. We examined the very high quality cryo-EM map very carefully and cannot find any other density consistent with sialic acid. With essentially the complete structure of the protein, we repeated our uSTA-guided docking and found no other site. We cannot exclude the presence of another sialic acid binding site, but there is no experimental support for it. The map does disclose the aromatic binding site also predicted by uSTA; this site is formed by Y660 and Q675, the latter changes its conformation to accommodate the aromatic (**Supplementary Figure S18**). We speculate that the high quality of the map may be related in part to the rigidification of the structure by this aromatic binding. The physiological relevance of a rigidifying aromatic-binding pocket is currently unclear.

Importantly, the spike sialoside-binding site that we have identified now not only provides a first functional role for NTD, it is also coincident to mutational, deletion hotspots in alpha (H69, V70, Y145) and beta variants-of-concern (beta-strand LLAL^241-244^) (**Figure 5C**). Changes remove either key interacting residues [alpha variant: 69,70,145] or perturb structurally significant residues [beta variant: beta strand] that form the pocket. The ‘end-on’ binding we observed is quite different to the ‘side-on’ binding observed for heparin (*16*). Heparins are often bound in a non-sequence-specific, charge-mediated manner, consistent with such a ‘side-on’ mode. The location of three binding sites in the trimer, essentially at the extreme edges of the spike (**Figure 6C**), imposes significant geometric constraints for avidity enhancement through multivalency (*52, 53*).

Intriguingly we find a clear link between our data and genetic analyses in patients that have been correlated with severity of disease. This suggests not only that cell-surface glycans and the modulation of binding to spike-to-sialosides may have played a role in infection and disease progression but also now identifies two glycan-associated genes. Genetic variations of *LGALS3BP* (which produces galectin-blocker Gal-3-BP) and *B3GNT8* (which produces glycosyltransferaseβ3GnT8) that reduce corresponding gene products (Gal-3-BP and β3GnT8, respectively) prove beneficial. Strikingly, despite their independent identification here, both Gal-3-BP and β3GnT8 interact around a common glycan motif: the polyLacNAc chain-extension variants found in tetraantennary *N*-linked glycoproteins (**Figure 7D**). The simplest explanation is therefore that modulation of such glycoproteins has played a role in infection and disease progression.

Consistent with the sialoside ligands we identify here, these glycoproteins contain Sia-Gal-GlcNAc motifs within *N*-linked-polyLacNAc-chains. Notably, these motifs have recently been identified in the deeper human lung.(*54*)

These data also lead us to suggest that B-origin-lineage SARS-CoV-2 virus may have exploited glycan-mediated attachment to host cells (**Figure 7D**) using *N*-linked-polyLacNAc-chains as a foothold. Reduction of Gal-3-BP function would allow its target, the lectin Gal-3, to bind more effectively to *N*-linked-polyLacNAc-chains, thereby competing with SARS-CoV-2 virus. Similarly, loss of β3GnT8 function would ablate the production of foothold *N*-linked-polyLacNAc-chains directly denying the virus a foothold. We cannot exclude other possible mechanisms including, for example, the role of *N*-linked-polyLacNAc-chains in T-cell regulation.(*55*) Notably, this analysis of the influence of genetic variation upon susceptibility to virus was confined to ‘first wave’ patients infected with B-origin-lineage SARS-CoV-2. Our discovery here also that in B-lineage virus such binding to certain sialosides is ablated further highlights the dynamic role that sugar binding may play in virus evolution and may be linked, as was previously suggested for H5N1 influenza A virus, to changing sugar preference during zoonotic transitions (*1*). The specific binding of *N*-Ac-sialosides, found in humans, that we observed might have been a contributing factor in driving zoonosis. Moreover, our combined data and models may also support decades-old hypotheses(*19*) proposing the benefit of cryptic sugar-binding by pathogens that may be ‘switched on and off’ to drive fitness in a different manner (e.g. in virulence or zoonosis) as needed.

## Author Contributions

BG, CJB, TDWC and AJB ran NMR experiments. Under the supervision of AJB, CJB developed the underlying deconvolution algorithm, UnidecNMR. AJB developed the modified Bloch- McConnell *K*_D_ analysis method. CJB performed the processing, uSTA applications and surrounding analysis. CJB and AK assigned key uSTA spectra. PH, ALB purified spike protein and prepared samples for NMR analysis; PH and ALB conducted thermal denaturation assays. AK synthesized 9-BPC-Neu5Ac, expressed and purified NmCSS and Pd2,6ST, assigned proton peaks in ^1^H NMR spectra of sialosides, designed TOCSY NMR experiments and performed the SPR experiments. AMG, AK and XX synthesized **5**, **6** and **9**. AMG generated reagents for SPR studies. AK, AMG and AJB analysed SPR data. GP generated **7** and **8** and assigned their ^1^H NMR spectra. JH cloned spike-BAP; PW and MD expressed spike protein. AL aided design of protein-ligand NMR experiments, prepared samples and contributed to data analysis. AR is coordinating the GEN-COVID Consortium and supervised the genotype-phenotype correlation of glycosylation-associated genes. AG performed WES experiments. EB performed WES alignment and joint call which was supervised by SF. NP performed logistic regression analysis which was supervised by MG. MB and FF collected and supervised the collection of clinical data. CF and SD perform specific analysis on *B3GNT8* and *LGALS3BP* genes. LPD and QJS performed HA expression and assisted with experimental design. AMJJB and PIK performed the structural modelling of the SARS-CoV-2-spike•sugar interactions. AJB and AMJJB scored these against the NMR data to obtain structural models for the complex. VCA, CJB, AJB and BGD produced figures. JHN, YY and JL performed the EM experiments and analysis. IB, CS and KG produced the protein used in EM. BGD, AJB, TDWC and JHN supervised and designed experiments, analyzed data and wrote the manuscript. All authors read and commented on the manuscript.

## Acknowledgments

Upgrades of the 600 MHz and 950 MHz spectrometers were funded by the Wellcome Trust (Grant Ref: 095872/Z/10/Z) and the Engineering and Physical Sciences Research Council (Grant Ref: EP/R029849/1), respectively, and by the University of Oxford Institutional Strategic Support Fund, the John Fell Fund, and the Edward Penley Abraham Cephalosporin Fund. We thank the Wellcome Trust for support (20289/Z/16/Z and 100209/Z/12/Z). IB is an Investigator of the Wellcome Trust (106115/Z/14/Z). EM time was funded by a philanthropic gift to support COVID-19 research at the University of Oxford. The Rosalind Franklin Institute is supported by the EPSRC.

AMJJB and PIK acknowledge financial support from the European Union Horizon 2020 projects BioExcel (823830) and EOSC-Hub (777536) and the IMI-CARE project (101005077) We would like to thank Professor Xi Chen at the University of California at Davis for providing the plasmid encoding Pd2,6ST; Professor John Mascola at the NIH for the plasmids encoding influenza A virus (IAV) NC99 (H1N1) haemagglutinin (HA) variants; Mikhail Kutuzov and Omer Dushek for useful discussions and assistance with SPR experiments; Dr Andrew Quigley and Nadisha Gamage (Diamond Light Source) for help with thermal stability assays; Professor Christina Redfield for assistance with high field Bruker instrumentation; Dr Lindsay Baker for insights into the structural models.

## Data Availability Statement

Raw spectral data for Figures 1, 2, 3, and 4, the data used to plot Figures 4A, B, C and relevant uSTA code have been deposited on the University of Oxford ORAdata server at https://ora.ox.ac.uk/objects/uuid:121699de-68d5-4b6f-b77e-cee85895da6c All associated spectral data is also shown in **Supplementary Table S7**. HADDOCK datasets and results for this work can be found at https://github.com/haddocking/SARS-COV2-NTD-SIA-modelling and https://doi.org/10.5281/zenodo.4271288. All associated clinical data have been supplied in **Supplementary Tables S1-S6**, conducted under trial (NCT04549831, www.clinicaltrial.org). The uSTA software is free for academic use.

## Methods

### Protein expression and purification: SARS-CoV-2 spike

The gene encoding amino acids 1-1208 of the SARS-CoV-2 spike glycoprotein ectodomain [with mutations of RRAR > GSAS at residues 682-685 (the furin cleavage site) and KV > PP at residues 986-987, as well as inclusion of a T4 fibritin trimerisation domain], using the construct previously described as template,(*56*) was cloned into the pOPINTTGneo-BAP vector using the forward primer (5’-GTCCAAGTTTATACTGAATTCCTCAAGCAGGCCACCATGTTCGTGTTCCTGGTGCTG -3’) and the reverse primer (5’- GTCATTCAGCAAGCTTAAAAAGGTAGAAAGTAATAC -3’), resulting in an aviTag plus 6His in the 3’ terminus of the construct.

Expi293 cells (Thermofisher Scientific) were used to express the Spike-Bap protein. The cells were cultured in expi293 expression media (Thermofisher Scientific) and were transfected using PEI MAX 40kDa (Polyscience) if cells were >95% viable and had reached a density of between 1.5 - 2 x 10^6^ cells per ml. Following transfection, cells were cultured at 37 °C and 5% CO_2_ at 120rpm for 17h. Enhancers (6mM Valproic Acid, 6.5mM Sodium Propionate, 50mM Glucose - Sigma) were then added and protein was expressed at 30 °C for 5 days before purification.

The media in which the protein spike-bap was secreted was supplemented with 1x PBS buffer at pH 7.4 (1:1 v/v) and 5mM NiSO_4_. The pH was adjusted with NaOH to pH 7.4 and filtered using a 0.8um filter. The mixture was stirred at 150 rpm for 2 hours at room temperature. The Spike protein was purified on an Akta Express system (GE Healthcare) using a 5mL His trap FF GE Healthcare column in PBS, 20mM imidazole pH 7.4 and eluted in PBS, 300mM imidazole pH 7.4. The protein was then injected on a High load superdex 200 16/600 gel filtration column (GE Healthcare) in deuterated PBS buffer pH 7.4. The eluted protein was concentrated using an Amicon Ultra-4 100kDa concentrator at 2000 RPM, 16°C, pre-washed multiple times with deuterated PBS) to a concentration of roughly 1 mg/mL.

### Protein expression and purification: Influenza HA

Freestyle 293-F cells were cultured in Freestyle expression media (Life Technologies) (37 °C, 8% CO_2_, 115 rpm orbital shaking). Cells were transfected at a density of 1x10^9^ cells/L, with pre-incubated expression vector (300 µg/L) and polyethyleneimine (PEI) MAX (Polysciences) (900 µg/L). Expression vectors encoded terminally His-tagged wild-type influenza A virus (IAV) NC99 (H1N1) hemagglutinin (HA) or a ΔRBS mutant previously described (*57*). After 5 days, supernatant was harvested, and protein was purified via immobilised metal chromatography.

### Errors

The errors in the transfer efficiencies were estimated using a bootstrapping procedure. Specifically sample STD spectra were assembled through taking random combinations with replacement of mixing times, and the analysis to obtain the transfer efficiency was performed on each. This process was repeated 100 times to enable evaluation of the mean and standard deviation transfer efficiency for each residue. Mean values correspond well with the value from the original analysis, and so we take the standard deviation as our estimate in uncertainty.

### Reagent sources

6’-Sialyllactose sodium salt and 3’-Sialyllactose sodium salt were purchased from Carbosynth and used directly: 6’-Sialyllactose sodium salt- CAS-157574-76-0, 35890-39-2, 3’-Sialyllactose sodium salt - CAS-128596-80-5,35890-38-1. Bovine serum Albumin (BSA) and L-tryptophan were purchased from Sigma Aldrich. Heparin sodium salt, from porcine intestinal mucosa, IU≥100/mg was purchased from Alfa Aesar. All other chemicals were purchased from commercial suppliers (Alfa Aesar, Acros, Sigma Aldrich, Merck, Carbosynth, Fisher, Fluorochem, VWR) and used as supplied, unless otherwise stated.

### Synthesis of 9-BPC-6’-Sialyllactose (6)

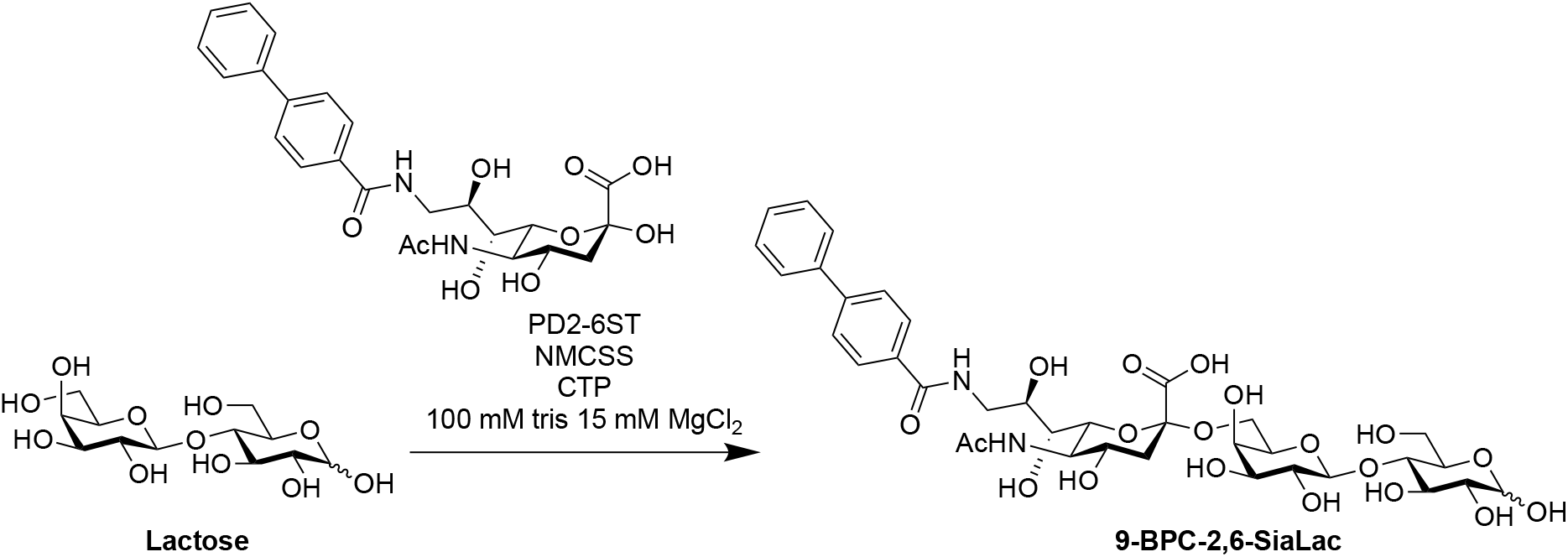

Lactose (30 mg, 0.087 mmol, 1 eq), BPC-Neu5Ac (48 mg, 0.0974 mmol, 1.12 eq, prepared using a procedure adapted from ref. (*58*)) and CTP disodium salt (92 mg, 1.74 mmol, 2 eq) were dissolved in buffer 100 mM Tris, 15 mM MgCl_2_ pH 8.5, 8.39 mL- to give achieve a final concentration of 10 mM with respect to the lactose acceptor. Pd-26ST enzyme (261 mL, 1.2 wt%, 1.38 mg/mL) and NmCSS enzyme (107.4 mL, 1 wt %, 2.73 mg/mL) were added to the reaction and the mixture shaken at 37 °C. After 22 h the reaction mixture was concentrated *in vacuo* and then purified via column chromatography (EtOAc:IPA:H_2_O 3:2:1à2.5:2:1 and then EtOAc:IPA:H_2_O: 4:2:1).

Fractions containing the desired compound were concentrated, redissolved in water and lyophilised to yield the product as a white powder (17.7 mg, 0.0218 mmol, 25%) – Note: the reaction conversion was very high, however the yield is reflective of difficult isolation of the product. **LRMS:** m/z (ES-) 811 [M-H]^-^; **HRMS**: m/z (ESI-): calc. for C_36_H_47_O_19_N_2_ [M-H]^-^ 811.2779, found 811.2773; **^1^H NMR** (400 MHz, D_2_O) δ 7.90 (2H, d, *J* = 8.5 Hz, ArH), 7.82 (2H, d, *J* = 8.3 Hz, ArH), 7.80 – 7.71 (2H, m), 7.61 – 7.53 (2H, m), 7.53 – 7.45 (1H, m), 5.16 (0.4H, d, *J* = 3.8 Hz, H-1a (alpha)), 4.63 (1H, d, *J* = 8.0 Hz, H-1a (beta)), 4.41 (1H, d, *J* = 7.8 Hz, H-1b), 4.09 (1H, ddd, *J* = 8.8, 7.2, 3.2 Hz, H-8c), 4.04 – 3.48 (m, 16H), 3.35 – 3.26 (1H, m, H-2a (beta)), 2.73 (1H, dd, J = 12.4, 4.7 Hz, H-3c_eq_), 2.02 (3H, s, NCOCH_3_), 1.77 (1H, t, *J* = 12.2 Hz, H-3c_ax_).

### Synthesis of Heparin Tetrasaacharides (7, 8)

Heparin sodium salt (UFH, 100mg) was dissolved in heparinise (Hep-1) preparations in 100mM Tris Acetate, 5Mm Ca(OAc)_2_, pH 6.8 to achieve a final concentration of 30 mg/mL UFH and 1.5mg/mL Hep-1-SUMO. The reaction was incubated in a water bath at 28°C and terminated after 36hrs by heating at 95 °C for 5 min. Upon centrifugation (2000g, 2min), the supernatant was filtered using a 0.2μm filter (Regenerated Cellulose, Sartorius) and lyophilised to yield a mixture of heparin oligosaccharides as a foam. These were then fractionated based on their degree of polymerisation on a Superdex Peptide 10/300 column (GE-Healthcare), following a procedure adapted from ref. (*59, 60*). Briefly, the column was equilibrated before each run with 2CV of 0.5M (NH_4_)_2_CO_3_ and the elution was monitored at 232nm. Sample application (100uL injection per run, ∼10mg in sugar content) was followed by an isocratic elution with 0.5M (NH_4_)_2_CO_3_ at a flow rate of 0.6mL/min. The fraction collector was set to collect 1mL fractions until well-resolved peaks were detected; at that point, fractionation was manually controlled and tetrasaccharide-containing fractions from sequential chromatographic runs were pooled together and subjected to serial lyophilisation rounds. Approximately 15mg of heparin tetrasaccharides were recovered upon removal of the volatile salt; these were further analysed with strong anion exchange HPLC on a SAX Propac PA1 column (9x250mm, Thermo Scientific) using H_2_O, pH 3.5 as solvent A and 2M NaCl (HPLC grade) pH 3.5 as solvent B. The column was equilibrated with solvent A for 30min prior to sample application (500uL injection per run, 10mg/mL) and the target oligosaccharides were eluted following a linear gradient from 30% to 70% solvent B over 180min, at a flow rate of 1mL/min. Fractions corresponding to the same retention times were pooled, neutralized with saturated NaHCO_3_ (HPLC grade) and lyophilized. Desalting was performed on an AKTA Purifier system (GE healthcare) by connecting three 5mL HiTrap Desalting columns (GE Healthcare) in a row. Elution with water was performed at a flow rate of 6mL/min and monitored at 232nm; UV absorbing fractions were pooled and lyophilized to yield the pure target tetrasaccharides.

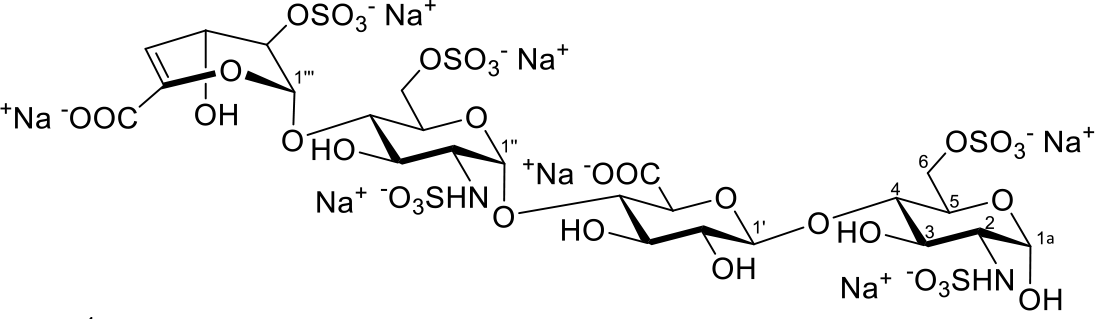

Compound **7**. White solid. ^1^H NMR (700 MHz, D_2_O) δ 5.91 (dd, *J_4’’’,3’’’_* 4.7 Hz, *J_4’’’,2’’’_* 1.4 Hz, 1H, H- 4’’’), 5.50 (d, *J_1’’,2’’_* 3.8 Hz, 1H, H-1’’), 5.43 (d, *J_1’’’,2’’’_* 2.6 Hz, 1H, H-1’’’), 5.39 (d, *J_1,2_* 3.5 Hz, 1H, H-1), 4.55 (m, 1H, H-2’’’), 4.52 (d, *J_1’,2’_* 8.0 Hz, 1H, H-1’), 4.29 – 4.26 (m, 3H, H-6a’’, 6a, 6b), 4.25 (dd, *J_3’’’,4’’’_* 4.8 Hz, *J_3’’’,2’’’_* 2.1 Hz, 1H, H-3’’’), 4.13 (dd, *J_6b’’, 6a’’_* 11.3 Hz, *J_6b’’,5’’_* 2.0 Hz, 1H, H-6b’’), 4.08 (ddd, *J_5,4_* 9.7 Hz, *J_5,6a_* 4.2 Hz, *J_5,6b_* 2.7 Hz, 1H, H-5), 3.91 (ddd, *J_5’’,4’’_* 10.1 Hz, *J_5’’,6a’’_* 4.2 Hz, *J_5’’,6b’’_* 2.2 Hz, 1H, H-5’’), 3.79 – 3.70 (m, 4H, H-3’, H-4’, H-5’, H-4’’), 3.68 – 3.59 (m, 2H, H-4, H-3), 3.56 (dd, *J_3’’,2’’_* 10.6 Hz, *J_3’’,4’’_* 8.8 Hz, 1H, H-3’’), 3.31 (dd, *J_2’,3’_* 9.5 Hz, *J_2’,1’_* 7.9 Hz, 1H, H-2’), 3.23 (dd, *J_2’’,3’’_* 10.6 Hz, *J_2’’,1’’_* 3.8 Hz, 1H, H-2’’), 3.20 (dd, *J_2α,3α_* 10.2 Hz, *J_2α,1α_* 3.6 Hz, 1H, H-2), 2.97 (dd, *J_2β,3β_* 10.0 Hz, *J_2β,1β_* 8.4 Hz, H-2β). ^13^C NMR (176 MHz, D_2_O) δ 169.3 (C-6’’’ - COOH), 160.2 (C-6’ – COOH), 144.8 (C-5’’’), 106.0 (C-4’’’), 102.1 (C-1’), 97.7 (C-1’’), 97.1 (C-1’’’), 91.0 (C-1), 78.8 (C-4), 78.0 (C-4’’), 77.5 (C-4’), 76.4, 75.9 (C-3’,5’), 74.5 (C-2’’’), 72.9 (C-2’), 69.53 (C-3’’), 69.3 (C-3), 68.7 (C-5’’), 68.1 (C-5), 66.7 (C-6), 66.1 (C-6’’), 62.8 (C-3’’’), 57.6 (C-2), 57.5 (C-2’’). Data consistent with previous reports^4^. LRMS m/z (ESI-): Found 535.5 [M-2H]^2-^; HRMS: m/z (ESI-) calc. for C_24_H_36_N_2_O_35_S_5_^2-^ [M-2H]^2-^ 535.9857, found 535.9858.

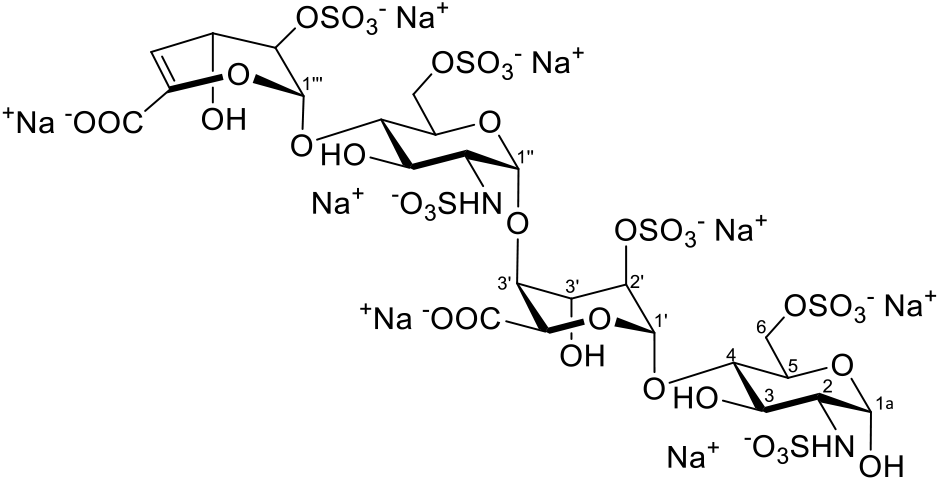

Compound **8**. White solid. ^1^H NMR (700 MHz, D_2_O) δ 5.91 (dd, *J_4’’’,3’’’_* 4.8 Ηz, *J_4’’’,2’’’_* 1.4 Hz, 1H, Η- 4’’’), 5.43 (d, *J_1’’’,2’’’_* 1.9 Hz, 1H, Η-1’’’), 5.38 (dd, *J* 3.6, 1.7 Hz, 2H, H-1, H-1’’), 5.13 (d, *J_1’,2’_* 3.4 Hz, 1H, H-1’), 4.67 (d, *J_5’,4’_* 3.0 Hz, 1H, H-5’), 4.62 (d, *J_1β,2β_* 8.3 Hz, 0.1H, H-1β), 4.55 (m, 1H, Η-2’’’), 4.30 (dd, *J_6a,6b_* 11.4 Hz, *J_6a,5_* 4.9 Hz, 1H, H-6a), 4.27 (dd, *J_6b’’,6a’’_* 10.7 Hz, *J_6b’’,_*_5’’_ 1.6 Hz, 1H, H-6b’’), 4.25 – 4.21 (m, 3H, H-6b, H-2’, H-3’’’), 4.18 (dd, *J_6a’’,6b’’_* 11.3 Hz, *J_6a’’,5’’_* 2.0 Hz, 1H, H-6a’’), 4.12 (dd, *J_3’,2’_* 6.7 Hz, *J_3’,4’_* 3.9 Hz, 1H, H-3’), 4.07 – 4.02 (m, 2H, H-4’, H-5), 3.97 (m, 1H, H-5’’), 3.75 (at, *J* 9.5 Hz, 1H, H-4’’), 3.67 (at, *J* 9.4 Hz, 1H, H-4), 3.62 (dd, *J_3,2_* 10.4 Hz, *J_3,4_* 8.9 Hz, 1H, H-3), 3.55 (dd, *J_3’’,2’’_* 10.6 Ηz, *J_3’’,4’’_* 8.8 Hz, 1H, H-3’’), 3.22 (dd, *J_2’’,3’’_* 10.6 Hz, *J_2’’,1’’_* 3.6 Hz, 1H, Η-2’’), 3.18 (dd, *J_2α,3_* 10.3 Hz, *J_2α,1α_* 3.6 Hz, 1H, Η-2α), 2.97 (dd, *J_2β,3_* 10.1 Ηz, *J_2β,1β_* 8.4 Hz, H-2β, 0.1H). ^13^C NMR (176 MHz, D_2_O) δ 174.4 (C-6’ – COOH), 169.2 (C-6’’’ – COOH), 144.8 (C-5’’’), 105.8 (C-4’’’), 99.4 (C-1’), 97.2 (C-1’’’), 96.5 (C1’’), 91.0 (C-1), 78.1 (C-4’’), 76.9 (C-4), 76.6 (C-2’), 76.3 (C-4’), 74.4 (C-2’’’), 69.7 (C-5’), 69.6 (C-3’), 69.5 (C-3’’), 69.4 (C-3), 68.8 (C-5’’), 68.4 (C-5), 67.0 (C-6), 66.2 (C-6’’), 62.7 (C-3’’’), 57.9 (C-2), 57.6 (C-2’’). Data consistent with previous reports^4^. LRMS m/z (ESI-): Found 575.5 [M-2H]^2-^; HRMS: m/z (ESI-) calc. for C_24_H_36_N_2_O_38_S_6_^2-^ [M-2H]^2-^ 575.9641, found 575.9651.

### Protein NMR Experiments

All NMR experiments in **NMR Methods Table 1** were conducted at 15 °C on a Bruker AVANCE NEO 600 MHz spectrometer with CPRHe-QR-1H/19F/13C/15N-5mm-Z helium-cooled cryo-probe. Samples were stored in a Bruker SampleJet sample loader while not in magnet, at 4 °C.

1D 1H NMR spectra with w5 water suppression were acquired using the Bruker pulse sequence zggpw5, using the smooth square Bruker shape SMSQ.10.100 for the pulsed field gradients. The spectrum was centered on the water peak, and the receiver gain adjusted. Typical acquisition parameters were sweep width of 9615.39 Hz, 16 scans per transient (NS), with 4 dummy scans, 32768 complex points (TD) and a recycle delay (d1) of 1 s for a total acquisition time of 54 s. Reference 1D spectra of protein only were acquired similarly with 16384 scans per transient with a total acquisition time of 12.5 hrs.

An STD experiment with excitation sculpted water suppression was developed from the Bruker pulse sequence stddiffesgp.2. The saturation was achieved using a concatenated series of 50ms Gaussian shaped pulses to achieve the desired total saturation time (d20). The shape of the pulses was specified by the Bruker shape file Gaus.1.1000, where the pulse is divided into 1000 steps and the standard deviation for the Gaussian shape is 165 steps. The field of the pulse was set to 200Hz, which was calculated internally through scaling the power of the high power 90° pulse.

The total relaxation delay was set to 5s, during which the saturation pulse was applied. The experiment was acquired in an interleaved fashion, with each individual excitation frequency being repeated 8 times (L4) until the total desired number of scans was achieved. Again, the spectrum was centred on the water peak, and the receiver gain optimized. Following recording of the FID, and prior to the recycle delay, a pair of water selective pulses are applied to destroy any unwanted magnetization. For all gradients (excitation sculpting and spoil), the duration was 3 ms using the smooth-square shape SMSQ10.100. Typical acquisition parameters were sweep width of 9615.39 Hz with typically 128 scans per transient (NS=16 * L4=8), 32768 complex points in the direct dimension and 16 dummy scans were executed prior to data acquisition.

In a typical experiment, two excitation frequencies were required, one exciting protein, and one exciting far from the protein (+20,000Hz, +33ppm from the carrier). A range of mixing times were acquired to allow us to carefully quantify the buildup curve to obtain K_D_ values. A typical set of values used was 0.1s, 0.3s, 0.5s, 0.7s, 0.9s, 1.1s, 1.3s, 1.5s, 1.7s, 1.9s, 2.0s, 2.5s, 3.0s, 3.5s, 4.0s and 5.0s.

Off and on-resonance spectra were acquired for 16 saturation times, giving a total acquisition time of 8.7 hrs

The experiment was acquired as a pseudo 3D experiment, with each spectrum being acquired at a chosen set of excitation frequencies and mixing times. Relaxation delays were set to 10 s for BSA + tryptophan STDs, and were 5 s otherwise.

### NMR Methods Table 1

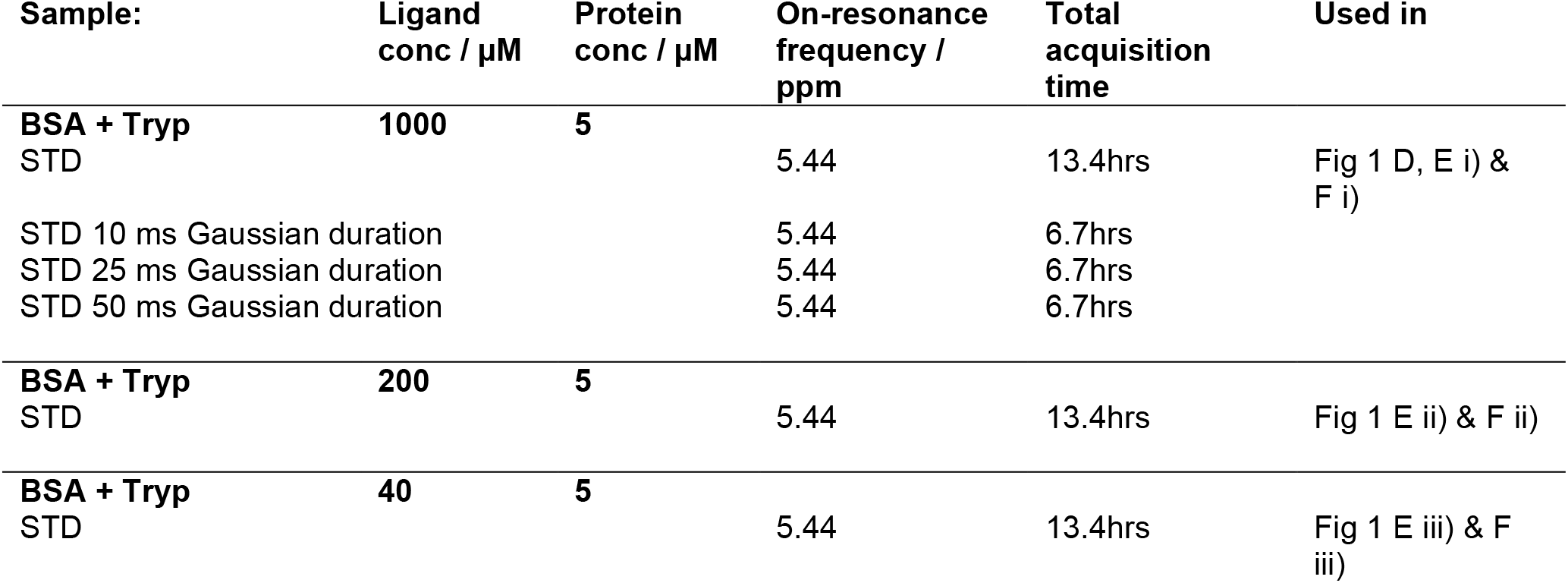
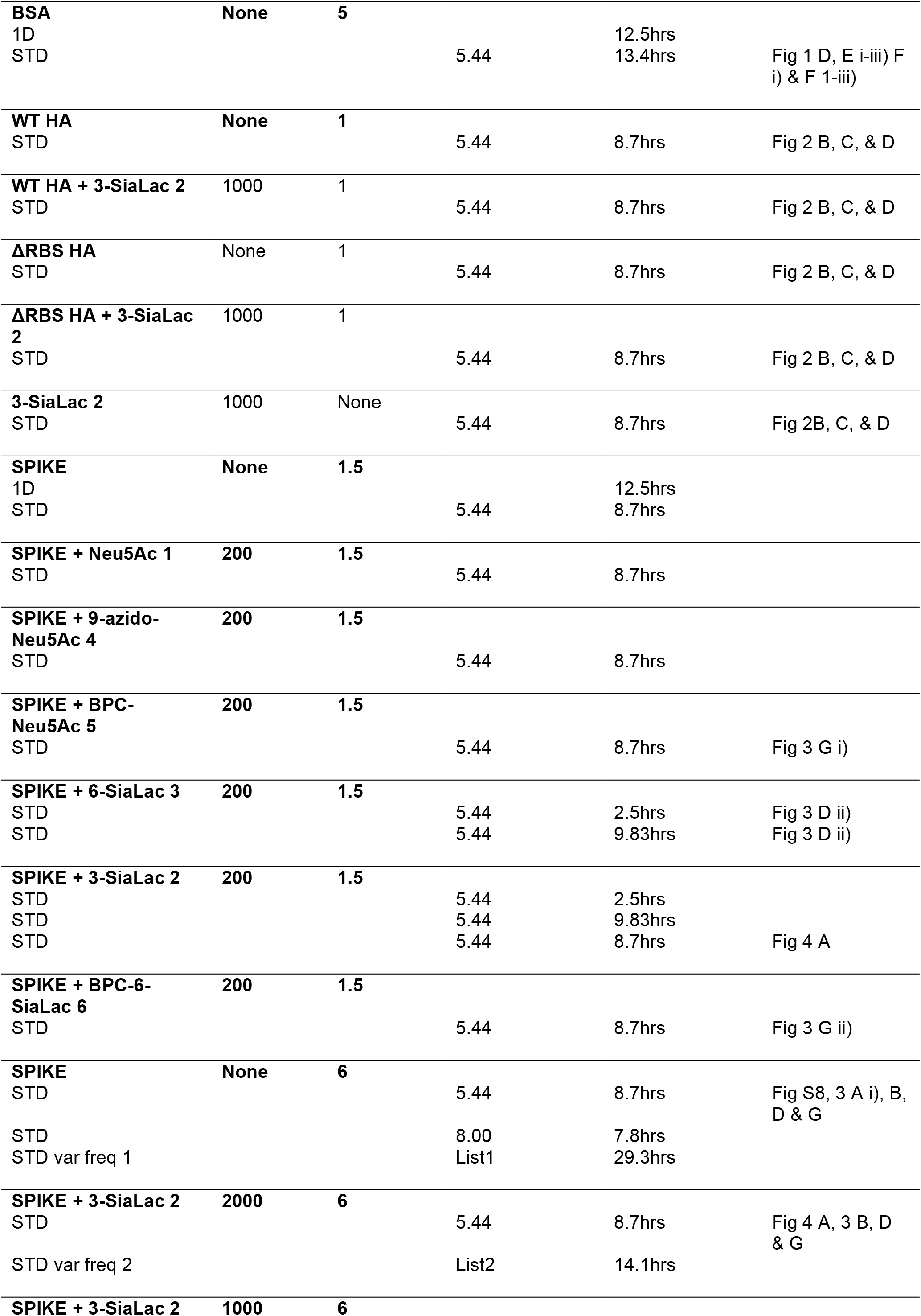
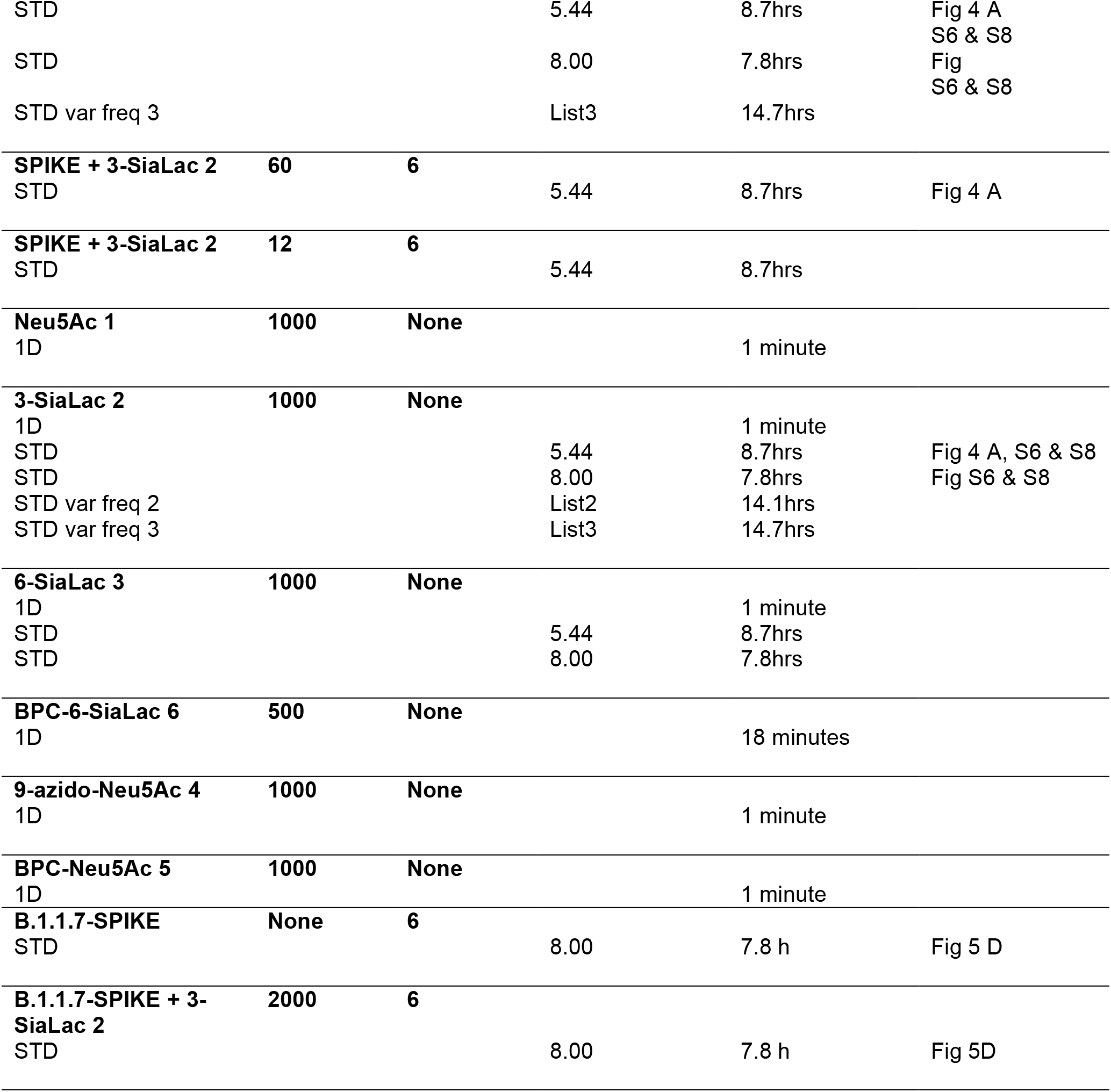

For STD 10-50 ms Gaussian experiments, the saturation times used were every other time from the default STD: 0.1 s, 0.5 s, 0.9 s, 1.3 s, 1.7 s, 2 s, 3 s, 4 s.

List1: For STD var freq 1, the on-resonance frequencies in Hz relative to an offset of 2820.61 Hz are: 337.89, 422.36, 524.93, 736.11, 914.10, 1276.12, 1336.46, 1380.21, 1458.64, 1556.69, 1693.96, 2494.93, 2597.50, 2790.58, 2930.86, 3362.27, 3663.95, 3986.75, 4099.88, 4326.15, 4484.53, 4703.25, 4896.33, 5824.01, 6006.53 and 6208.65. The saturation times used were 2 s, 3 s, 4 s and 5 s.

List2: For STD var freq 2, the on-resonance frequencies in Hz relative to an offset of 2820.61 Hz are: -2399.99, -1979.99, -1530.00, -1050.01, -330.021, 338.096, 1679.95, 1829.94, 1979.94, 2129.94, 2279.94 and 2579.93. The saturation times used were 0.1 s, 0.5 s, 2 s and 5 s.

List3: For STD var freq 3, the on-resonance frequencies in Hz relative to an offset of 2820.61 Hz are: -2579.98, -2459.99, -2339.99, -2039.99, -1488.00, -1120.03, -345.02, 311.97, 1079.96, 1379.95, 1679.95, 1979.94, 2279.94 and 2579.93. The saturation times used were 0.1 s, 0.3 s, 0.5 s and 0.9 s.

List4: For STD var freq 4, the on-resonance frequencies in Hz relative to an offset of 2820.61 Hz are: -2461.55, -1973.77, -1518.69, -1270.72, -693.69, -274.80, 280.08, 808.02, 1047.05, 2055.57, 2630.61 and 2979.65. The saturation times used were 0.1 s, 0.5 s, 0.9 s, 2 s, 3 s and 4 s.

Spectra were also acquired on a 600 MHz spectrometer with Bruker Avance III HD console and 5mm TCI CryoProbe, running TopSpin 3.2.6, recorded in **NMR Methods Table 2**, and a 950 MHz spectrometer with Bruker Avance III HD console and 5mm TCI CryoProbe, running TopSpin 3.6.1, recorded in **NMR Methods Table 3**. The 950 used a SampleJet sample changer. Samples were stored at 15 °C. The parameters used for the STD experiments were the same as above, with the following varying by instrument:

On the 600, typical acquisition parameters were sweep width of 9615.39 Hz with typically 128 scans per transient (NS=16 * L4=8), 32768 complex points in the direct dimension and 2 dummy scans, executed prior to data acquisition.

On the 950, typical acquisition parameters were sweep width of 15243.90 Hz with typically 128 scans per transient (NS=16 * L4=8), 32768 complex points in the direct dimension and 2 dummy scans, executed prior to data acquisition.

### NMR Methods Table 2

**NMR Methods Table 2:**
| Sample: | Ligand conc. / $\mu\text{M}$ | Protein conc. / $\mu\text{M}$ | On-resonance frequency / ppm | Total acquisition time | Used in |
| --- | --- | --- | --- | --- | --- |
| **Trp**<br>STD var freq 4 | **1000** | **None** | List 4 | 19.2hrs | Fig S7 |
| **BSA**<br>STD var freq 4 | **None** | **5** | List 4 | 19.2hrs | Fig S7 |
| **BSA**<br>STD | **None** | **10** | 5.44 | 7.9hrs |  |
| **BSA + Trp**<br>STD var freq 4 | **1000** | **5** | List 4 | 19.2hrs | Fig S7 |
| **BSA + Trp**<br>STD | **25** | **10** | 5.44 | 11.9hrs |  |
| **SPIKE + Neu5Ac 1**<br>STD | **2000** | **3** | 8.00 | 7.9hrs |  |
| **SPIKE**<br>STD | **None** | **3** | -1.00 | 7.9hrs | Fig 5 A & B |
| **Heparin B**<br>STD | **500** | **None** | -1.00 | 7.9hrs | Fig 5 A |
| **Heparin C**<br>STD | **500** | **None** | -1.00 | 7.9hrs | Fig 5 B |
| **SPIKE + Heparin B**<br>STD | **500** | **3** | -1.00 | 7.9hrs | Fig 5 A |
| **SPIKE + Heparin C** | **500** | **3** |  |  |  |
| STD |  |  | -1.00 | 7.9hrs | Fig 5 B |
| **SPIKE B.1.351** | **None** | **6** |  |  |  |
| STD |  |  | -1.00 | 7.9hrs | Fig 5 D |
| STD |  |  | 5.44 | 7.9hrs |  |
| **3-SiaLac 2** | **1000** | **None** |  |  |  |
| STD |  |  | -1.00 | 7.9hrs | Fig 5 D |
| **SPIKE B.1.351 + 3-SiaLac 2** | **60** | **6** |  |  |  |
| STD |  |  | -1.00 | 7.9hrs | Fig 5 D |
| **SPIKE B.1.351 + 3-SiaLac 2** | **200** | **6** |  |  |  |
| STD |  |  | -1.00 | 7.9hrs | Fig 5 D |
| **SPIKE B.1.351 + 3-SiaLac 2** | **1000** | **6** |  |  |  |
| STD |  |  | -1.00 | 7.9hrs | Fig 5 D |
| **SPIKE B.1.351 + 3-SiaLac 2** | **2000** | **6** |  |  |  |
| STD |  |  | -1.00 | 7.9hrs | Fig 5 D |

### NMR Methods Table 2.1

**NMR Methods Table 2.1:**
| Sample: | Ligand conc. / $\mu\text{M}$ | Protein conc. / $\mu\text{M}$ | Nanobody conc. / $\mu\text{M}$ | On-resonance frequency / ppm | Total acquisition time | Used in |
| --- | --- | --- | --- | --- | --- | --- |
| **SPIKE** | **None** | **6** | **None** |  |  |  |
| STD |  |  |  | 8.00 | 7.9hrs | Fig 5 E i) & F |
| **3-SiaLac 2** | **2000** | **None** | **None** |  |  |  |
| STD |  |  |  | 8.00 | 7.9hrs | Fig 5 E & F |
| **C5 Nanobody** | **None** | **None** | **6** |  |  |  |
| STD |  |  |  | 8.00 | 7.9hrs |  |
| **SPIKE + 3-SiaLac 2** | **2000** | **None** | **6** |  |  |  |
| STD |  |  |  | 8.00 | 7.9hrs | Fig 5 E i) & F |
| **SPIKE + C5 Nanobody** | **None** | **6** | **6** |  |  |  |
| STD |  |  |  | 8.00 | 7.9hrs | Fig 5 E ii) & F |
| **SPIKE + 3-SiaLac 2 + C5 Nanobody** | **2000** | **6** | **6** |  |  |  |
| STD |  |  |  | 8.00 | 7.9hrs | Fig 5 E ii) & F |
| **3-SiaLac 2 + C5 Nanobody** | **2000** | **None** | **6** |  |  |  |
| STD |  |  |  | 8.00 | 7.9hrs | Fig 5 E ii) & F |

### NMR Methods Table 3

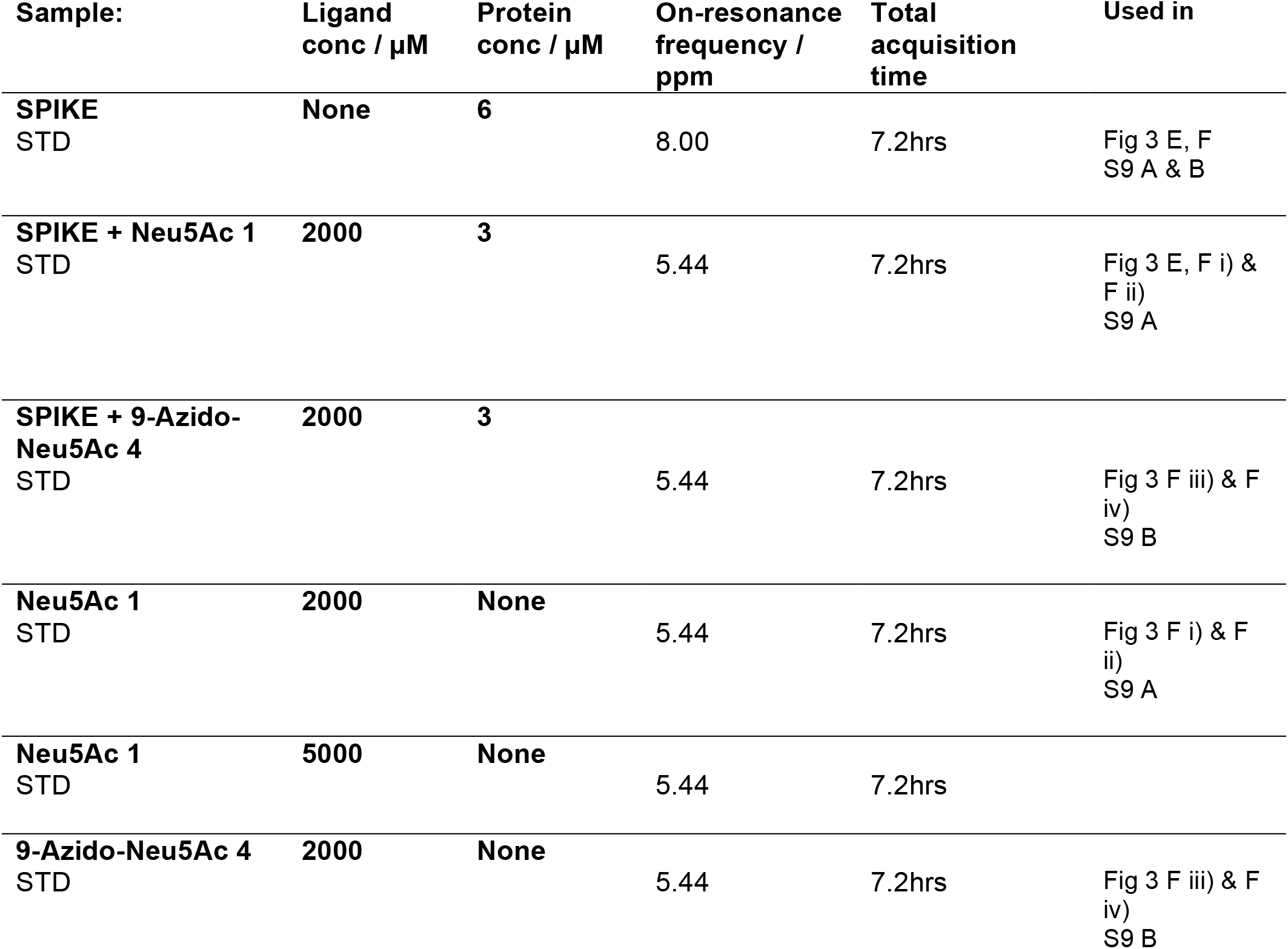

### uSTA Data Analysis

NMR spectra with a range of excitation frequencies and mixing times were acquired on ligand only, protein only, and mixed protein/ligand samples (**Supplementary Figure S6**).

To analyse an STD dataset, two projections were created by summing over all 1D spectra, and summing over all corresponding STD spectra. These two projections provide exceptionally high signal to noise, suitable for detailed analysis and reliable peak detection. The UnidecNMR algorithm was first executed on the raw spectra, to identify peak positions and intensities. Having identified possible peak positions, the algorithm then analyses the STD spectra but only allowing resonances in places already identified in the 1D spectrum. Both analyses are conducted using the protein only baselines for accurate effective subtraction of the protein baseline without the need to use relaxation filters (**Supplementary Figure S8**).

The results for the ligand only spectra were first analysed. In each case, excellent agreement with the known assignments was obtained, providing us with confidence in the algorithm. The mixed protein/ligand spectrum was then analysed, which returns very similar results to ligand only case. Contributions from the protein, although small however, are typically evident in the spectra justifying the explicit inclusion of the protein only baseline during the analysis. When analysing the mixture, we included the protein only background as a peak shape whose contribution to the spectrum can be freely adjusted. In this way, the spectra of protein/ligand mixtures could be accurately and quickly deconvolved, with the identified ligand resonances occurring in precisely the positions expected from the ligand only spectra. The results from the previous steps were then used to analyse the STD spectra. As these have much lower signal to noise, we fixed the ligand peak positions to be only those previously identified. Otherwise the protocol performed as described previously, where we used a protein only STD data to provide a baseline.

These analyses allow us to define a ‘transfer efficiency’, which is simply the ratio of the signal from a given multiplet in the STD spectrum, to the total expected in the raw 1D experiment. To obtain ‘per atom’ transfer efficiencies, signal from the various pre-assumed components on the multiplets from each resonance were first summed before calculating the ratio. In the software, this is achieved by manually annotating the initial peak list using information obtained from independent assignment experiments (see **Supplementary Figures S19,S20**).

Over the course of the project, it became clear that subtracting the transfer efficiencies obtained from a ligand only sample was an essential part of the method (**Supplementary Figure S9,S10**). Depending on the precise relationship between the chemical shift of excitation, the location of the ligand peaks, and the excitation profile of the Gaussian train, we observed small apparent STD transfer in the ligand only sample that cannot be attributed to ligand binding, arising from a small residual excitation of ligand protons, followed by internal cross relaxation. It is likely that this excitation occurs at least in part via resonances of the ligand that are exchange broadened, such as OH protons, which are not directly observed in the spectrum. When exciting far from the protein, zero ligand excitation is observed, as we would expect, but when exciting close to the methyls, or in the aromatic region, residual ligand excitation could be detected in ligand only samples (**Supplementary Figure S9,S10**). Without the ligand only correction, the uSTA surface may appear to be highly dependent on choice of excitation frequency. However, with the ligand correction, the relative uSTA profiles become invariant with excitation frequency. In general therefore, we advise acquiring these routinely, and so the uSTA analysis assumes the presence of this data (**Supplementary Figure S6,S8**). The invariance of relative transfer efficiency with excitation frequency suggest that the internal evolution of magnetization within the protein during saturation (likely on the ms timescale) is much faster than the effective cross relaxation rate between protein and ligand (occurring on the seconds timescale).

Having identified the relevant resonances of interest, and performed both a protein, and residual ligand subtraction, the spectra were then re-analysed without first summing over the different mixing times, in order to develop the quantitative atom-specific build-up curves. These were quantitatively analyzed as described below to obtain *K*_D_ and k_off_ rates. The values we obtain performing this analysis on BSA/Trp closely match those measured by ITC, and the values we measure for ligand **2** and SPIKE are in good agreement with those measured by SPR as described in the text.

The coverage of protons over the ligands studied here was variable, as for example, there are no protons on carboxyl groups. To enable a complete surface to be rendered, the transfer efficiencies for each proton were calculated as described above, and the value is then transferred to the adjacent heteroatom. For heteroatoms not connected to an observed proton, a 1/r^6^ weighted average score was calculated. This approach allows us to define a unique surface. Caution should be exercised when quantitatively interpreting such surfaces where there are no reliable measurements of the heteroatom.

In practice, raw unformatted FIDs are submitted to the uSTA pipeline, and the various steps are performed largely automatically, where a user needs to manually adjust processing settings such as phasing and choosing which regions to focus on, iteratively adjust the peak shape to get a good match between the final reconvolved spectrum and the raw data, and input manual atomic assignments for each observed multiplet. The uSTA analysis pipeline then provides a user with a report that shows the results of the various stages of analysis, and uses pymol to render the surfaces. The final transfer efficiencies delivered by the program can be combined with a folder containing a series of HADDOCK models to provide final structural models (**Figure 5**).

### Quantitative analysis via uSTA

In principle, a complete description of the saturation transfer experiment can be achieved via the Bloch-McConnell equations. If we can setup a density matrix describing all the spins in the system, their interactions and their rates of chemical change in an evolution matrix R, then we can follow the system with time according to:

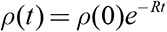

The challenge comes from the number of spins that needs to be included, and the need to accurately describe all the interactions between them, which will need to also include how these are modulated by molecular motions in order to get an accurate description of the relaxation processes. This is illustrated by the CORCEMA method,(*61*) that takes a static structure of a protein/ligand complex and estimates STD transfers. The calculations performed to arrive at cross relaxation rates assumes the complex is rigid, which is a poor approximation for a protein, and because of the large number of spins involved, the calculation is sufficiently intensive such that this calculation cannot be routinely used to fitted to experimental data.

It would be very desirable to extract quantitative structural parameters, as well as chemical properties such as interaction strengths and association/dissociation rates directly from STD data. In what follows we develop a simple quantitative model for the STD experiment to achieve this goal. We will treat the system as comprising just two spins, one to represent the ligand and one to represent the protein, and we allow the two spins to exist either in isolation, or in a bound state. We can safely neglect scalar coupling and so we only need to allow the x,y and z basis operators for each spin, together with an identity operator to ensure the system returns to thermal equilibrium at long times. As such, our evolution matrix R will be square matrices with 13 x 13 elements.

For the spin part, our model requires us to consider the chemical shift of the ligand in the free and bound states, and the chemical shifts of the protein in the free and bound states. In practice however, it is sufficient to set the free protein state on resonance with the pulse, and ensure that the free ligand chemical shift is far removed from the pulse as we would expect to have in the experiment.

The longitudinal and transverse relaxation rates are calculated for the free and bound states using a simple model assuming in each state there are two coupled spins separated by a distance R. In addition, cross relaxation between ligand and protein is allowed only when the two are bound. The relaxation rates are characterised by an effective distance, and an effective correlation time.

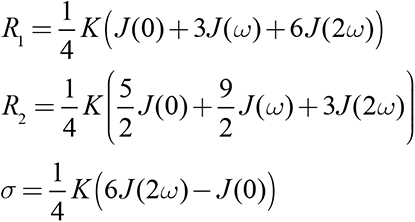

Which are each parameterized in terms of an interaction constant 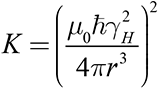 and a spectral density function 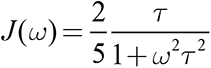. The longitudinal and transverse relaxation rates R_1_ and R_2_ describe auto-relaxation of diagonal z, and xy elements respectively. The cross-relaxation rates *σ* describes cross relaxation and couples z elements between the ligand and protein in the bound state. We ensure that the system returns to equilibrium at long times by adding elements of the form *R*_1_ *M*_0_ or *σ M*_0_ linking the identify element, and the z matrix elements. Overall, the relaxation part of the model is parametised by two correlation times, one for the ligand, and one for the protein/complex, and three distances, one for the ligand auto relaxation rates, one for protein auto relaxation rates, and one for the protein/ligand separation.

Finally, the chemical kinetics govern the rates at which the spins can interconvert. We will take a simple model where *PL* ⇄ *P* + *L* , whose dissociation constant is given by:

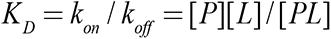

The free protein concentration can be determined from knowledge of the K_D_, and the total ligand and protein concentrations:

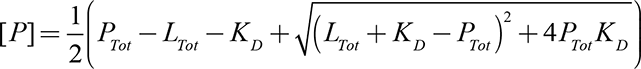

From which the bound protein concentration, and the free and bound ligand concentrations can be easily calculated.

The density matrix is initialised with the free and bound protein/ligand concentrations assigned to the relevant z operators. It was found to be important to additionally include a factor that accounts for the increased proton density within the protein. The saturation pulse is then applied either as a concatenated series of Gaussian pulses whose duration and peak power in Hz needs to be specified, exactly matching the pulse shapes and durations used in the experiment (see NMR methods above).

Build up curves and transfer efficiencies can be easily simulated using this model and compared to data, and the various parameters can be optimized to fit to the data. In total, the model is characterized by nine parameters: *K_D_*, *k_off_*, the correlation times of the ligand and the protein, the three distances described above and the proton density within the protein. There is substantial correlation between the effects of the various parameters. By obtaining data at various protein and ligand concentrations however, it is possible to break this degeneracy and obtain well described values as described in the text.

In practical terms, the initial rate of the buildup curve is predominantly affected by the cross-relaxation rate and the off rate, and the final height of the buildup curve is mostly influenced by the proton density in the protein and *K*_D_. Software to perform this analysis has been directly incorporated into the uSTA software.

### Parameters Fitted by the Model

Overall the model is parameterised by a set of values that characterise the intrinsic and cross relaxation. From tG and rIS(ligand) we estimate R_1_ and R_2_ of the ligand, from tE, rIS(protein) we obtain R_1_ and R_2_ of the protein, and from tE and rIS(complex) we calculate the cross relaxation rate. These values are combined with a factor that accounts for the larger number of spins present in the protein, ‘fac’, and the on and off rates, to complete a set of 8 parameters that specify our model. The distances should be considered ‘effective’ values that parasitize the relaxation rates though in principle it should be possible to obtain physical insights form their interpretation. The concentration independent relaxation rates can be separated from the exchange rates by comparing the curves as a ruction of ligand and protein concentration. By treating the system as comprising of two spins we are effectively assuming that the cross relaxation within the protein is very efficient. In the STD experiment, saturation pulses are applied for several second, which is sufficient for near-saturating spin diffusion within a protein.

### Thermostability assays

Thermal stability assays were performed using a NanoTemper Prometheus NT.48 (Membrane Protein Laboratory, Diamond Light Source). To 11 uL of 2 uM Spike (deuterated PBS), 2 uL of trisaccharide 2 (deuterated PBS) was titrated to give final concentrations of 0.1, 0.2, 0.4, 0.8, 1.6 and 2.0 mM. Samples were then loaded into capillaries and heated from 15 to 95 °C. Analysis was performed using PR.ThermControl v2.3.1 software.

### SPR binding measurement assays

All experiments were performed on a Biacore T200 instrument. For the immobilisation of SiaLac onto sensor chip a flow rate 10 μL/min was used in a buffer solution of HBP-EP (0.01 M HEPES pH 7.4, 0.15 M NaCl, 3 mM EDTA, 0.005% v/v Surfactant P20). A CM5 sensor chip (carboxymethylated dextran) was equilibrated with HBS-EP buffer at 20°C. The chip was activated by injecting a mixture of *N-*hydroxysuccinimide (50 mM) and EDC-HCl (200 mM) for 10 min followed by a 2 min wash step with buffer. Ethylenediamine (1 M in PBS) was then injected for 7 min followed by a 2 min wash step followed by ethanolamine-HCl (1 M, pH 8.5) for 10 min and then a further 1 min wash step. Finally, SiaLac-IME (5.6 mM in PBS) was injected over 10 min and a final 2 min wash step was performed (see **Supplementary Figure S13**)

For analysis of spike binding a flow rate 10 μL/min was used at 16 °C. Serial dilutions of spike (0.19, 0.50, 1.36, and 3.68 μM) were injected for 30 s association and 150 s dissociation starting with the lowest concentration. Buffer only runs were carried out before injection of spike and after the first two dilutions. BSA (3.03 μM in PBS) was used as a negative control and a mouse serum in a 100-fold dilution was used as a positive control.

### Analysis of SPR Data

To analyse the SPR data, we assume an equilibrium of the form *PL* ⇄ *P* + *L* characterised by a dissociation constant 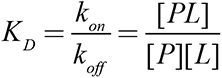. To follow the kinetics of binding and dissociation, we assume the SPR response Γ is proportional to the bound complex *Γ* = *κ*[*PL*], which leads to the following kinetic equation:

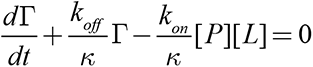

This can be solved when restrained by the total number of binding sites, *L_tot_* = [*L*]+[*PL*]. Under conditions of constant flow, we assume the free protein concentration is constant, which leads to the following:

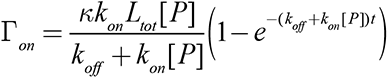

And similarly, for dissociation where we take the concentration of free protein to be zero:

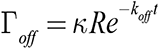

The recovery of the chip was not complete after each protein concentration and wash step, as has been observed for shear-induced lectin-ligand binding with glycans immobilised onto a chip surface.(*62*) Nevertheless, the data were well explained by a global analysis where the on and off rates were held to be identical for each replicate, but the value of k was allowed to vary slightly between runs, and an additional constant was introduced to Γ*_off_* to account for incomplete recovery of the SPR signal following standard approaches. Concentration of spike was insufficient to get the plateau region of the binding, and so the specific time values taken for the on rate affect the fitted values.

### Modelling of the N-terminal Domain of SARS-CoV-2 with Glycans

We modelled the structure of the N-terminal domain (NTD) on Protein Data Bank (PDB) entry 7c2l (*18*) since it provided significantly better coverage of the area of interest when compared to the majority of the templates available on the PDB as of July 15th 2020. The models were created with Modeller (*63*), using the ‘`automodel`’ protocol without refining the ‘`loop`’. We generated 10 models and ranked them by their DOPE score (*64*), selecting the top 5 for ensemble docking.

### Docking of 3’-sialyllactose to SARS-CoV-2 NTD

We docked 3’-sialyllactose to NTD with version 2.4 of the HADDOCK webserver. (*39, 40*) The binding site on NTD was defined by comparison with PDB entry 6q06 (*5*), a complex of MERS-CoV spike protein and 2,3-sialyl-N-acetyl-lactosamine. The binding site could not directly be mapped because of conformational differences between the NTDs of MERS-CoV and SARS-CoV- 2, but by inspection a region with similar properties (aromatics, methyl groups and positively charged residues) could be identified. We defined in HADDOCK the sialic acid as ‘active’ and residues 18, 19, 20, 21, 22, 68, 76, 77, 78, 79, 244, 254, 255, 256, 258 and 259 of NTD as ‘passive’, meaning the sialic acid needs to make contact with at least one of the NTD residues but there is no penalty if it doesn’t contact all of them, thus allowing the compound to freely explore the binding pocket. Since only one restraint was used we disabled the random removal of restraints.

Following our small molecule docking recommended settings (*65*) we skipped the ‘hot’ parts of the semi-flexible simulated annealing protocol (‘`initiosteps`’ and ‘`cool1_steps`’ set to 0) and also lowered the starting temperature of the last two substages to 500 and 300K, respectively (‘`tadinit2_t`’ and ‘`tadinit3_t`’ to 500 and 300, respectively). Clustering was performed based on ‘RMSD’ with a distance cut-off of 2Å and the scoring function was modified to:

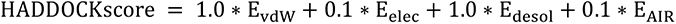

All other settings were kept to their default values. Finally, the atom-specific transfer efficiencies determined by uSTA were used to filter cluster candidates.

### Cryo-EM analysis

SARS-CoV-2 spike protein, purified as previously described(*44*), in 1.1 mg /ml was incubated with 10mM ethylbenzamide-tri-iodo Siallyllactose (please check) over night at 4 °C 3.5 µl sample was applied to glow-discharged Quantifoil gold R1.2/1.3 300-mesh grids and blotted for ∼ 3 s at 100% humidity and 6 °C before vitrification in liquid ethane using Vitrobot (FEI). Two datasets were collected on Titan Krios equipped with K2 direct electron detector at cryo-EM facility (OPIC) in the Division of Structural Biology, University of Oxford. Both datasets were collected by SerialEM, at a magnification of × 165,000 with the physical pixel size 0.82 Å/pixel. Defocus range was -0.8 µm to - 2.4 µm. Total dose for two datasets were 60 e/Å^2^ (5492 movies) and 61 e/Å^2^ (8284 movies) respectively.

Motion correction was performed by MotionCor2 (*66*) (2). The motion-corrected micrographs were imported into cryoSPARC (*67, 68*) and contrast transfer function values were estimated using Gctf (*69*) in cryoSPARC. Templates were produced by 2D classification from 5492 micrographs with particles auto-picked by Laplacian-of-Gaussian (LoG) based algorithm in RELION 3.0 (*70, 71*). Particles were picked from all micrographs using Template picker in cryoSPARC. Multiple rounds of 2D classification were carried out and the selected 2D classes (372,157 particles) were subjected to 3D classification (Heterogenous Refinement in cryoSPARC) using 6 classes. One class was predominant after 3D classification. Non-uniform Refinement (*68*) was performed for this class (312,018 particles) with C1 and C3 symmetry respectively, yielding 2.27 Å map for C3 symmetry and 2.44 Å map for C1 symmetry.

### Cryo-EM data collection, refinement and validation statistics

|  | Spike_C1<br>(EMDB-xxxx)<br>(PDB xxxx) | Spike_C3<br>(EMDB-xxxx)<br>(PDB xxxx) |
| --- | --- | --- |
| **Data collection and processing** |  |  |
| Magnification | 165,000 | 165,000 |
| Voltage (kV) | 300 | 300 |
| Electron exposure (e-/Å<sup>2</sup>) | 60; 61 | 60; 61 |
| Defocus range (μm) | -0.8 ~ -2.4 | -0.8 ~ -2.4 |
| Pixel size (Å) | 0.82 | 0.82 |
| Symmetry imposed | C1 | C3 |
| Initial particle images (no.) | 5,419,671 | 5,419,671 |
| Final particle images (no.) | 312,018 | 312,018 |
| Map resolution (Å) | 2.44 | 2.27 |
| FSC threshold | 0.143 | 0.143 |
| Map sharpening *B* factor (Å<sup>2</sup>) | 87.9 | 89.4 |
| **Refinement** |  |  |
| Initial model used (PDB code) |  | 7jji |
| Model resolution (Å) |  | 2.27 |
| FSC threshold |  | 0.143 |
| Model composition |  |  |
| Non-hydrogen atoms |  | 27168 |
| Protein residues |  | 3303 |
| Ligands |  | 1131 |
| Waters |  | 282 |
| Model data correlation |  |  |
| CC(mask) |  | 0.85 |
| CC(box) |  | 0.70 |
| CC(peaks) |  | 0.68 |
| CC(volume) |  | 0.84 |
| CC(ligand) |  | 0.60 |
| *B* factors (Å<sup>2</sup>) |  |  |
| Protein |  | 62 |
| Ligand |  | 81 |
| Water |  | 52 |
| R.m.s. deviations |  |  |
| Bond lengths (Å) |  | 0.002 |
| Bond angles (°) |  | 0.50 |
| Validation |  |  |
| MolProbity score |  | 1.64 |
| Clashscore |  | 4.9 |
| Poor rotamers (%) |  | 1.36 |
| Ramachandran plot |  |  |
| Favored (%) |  | 95.9 |
| Allowed (%) |  | 4.0 |
| Disallowed (%) |  | 0.1 |

### Genetic analysis of clinical samples

Variant calling: Reads were mapped to the hg19 reference genome by the Burrow-Wheeler aligner BWA. Variants calling was performed according to the GATK4 best practice guidelines.

Namely, duplicates were first removed by *MarkDuplicates*, and base qualities were recalibrated using *BaseRecalibration* and *ApplyBQSR*. *HaplotypeCaller* was used to calculate Genomic VCF files for each sample, which were then used for multi-sample calling by *GenomicDBImport* and *GenotypeGVCF*. In order to improve the specificity-sensitivity balance, variants quality scores were calculated by *VariantRecalibrator* and *ApplyVQSR*, and only variants with estimated truth sensitivity above 99.9% were retained. Variants were annotated by ANNOVAR.

Rare variant selection: Missense, splicing and loss of function variants with a frequency lower than 0.01 according to ExAC_NFE (Non Finnish European ExAC Database) were considered for further analyses. A score of 0 was assigned to each samples where the gene is not mutated and the score of 1 was assigned when at least one variant is present on the gene.

### Genes prioritization by Logistic regression

Discriminating genes in COVID-19 disease were interpreted in a framework of feature selection analysis using a customized feature selection approach based on the recursive feature elimination algorithm applied to the LASSO (Least Absolute Shrinkage and Selection Operator) logistic regression model. Specifically, for a set of *n* samples *x*_i_, *y*_i_ (*i* = 1, … , *n*) each of which consists of *p* input features **x*_i_,_k_* ∈ **χ*_i_ k* = 1, . . . , *p* and one output variable *y*_i_ ∈ *Y*, these features assumed the meaning of genes, whereas the samples were the patients involved in the study. The space *χ* = *χ*_1_×*χ*_2_. . .×*χ_p_* was denoted “input space”, whereas the “hypothesis space” was the space of all the possible functions *f*: *χ* → *Y* mapping the inputs to the output. Given the number of features (*p*) is substantially higher than the number of samples (*n*), LASSO regularization(*45*) has the effect of shrinking the estimated coefficients to zero, providing a feature selection method for sparse solutions within the classification tasks. Feature selection methods based on such regularization structures (Embedded methods) were most applicable to our scope because they were computationally tractable and strictly connected with the classification task of the ML algorithm.

As the baseline algorithm for the embedded method, we adopted the Logistic Regression (LR) model that is a state-of-the-art ML algorithm for binary classification tasks with probabilistic interpretation. It models the log-odds of the posterior success probability of a binary variable as the linear combination of the input:

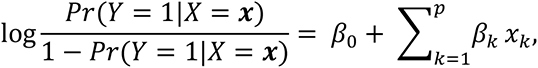

where ***x*** is the input vector, *β_i_* are the coefficients of the regression and *X* and *Y* are the random variables representing the input and the output respectively. The loss function to be minimized is given by the binary cross-entropy loss

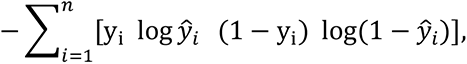

where 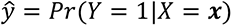 is the predicted target variable and *y* is the true label. As already introduced, in order to enforce both the sparsity and the interpretability of the results, the model is trained with the additional LASSO regularization term:

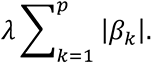

In this way, the absolute value of the surviving weights of the LR algorithm was interpreted as the feature importances of the subset of most relevant genes for the task. Since a feature ranking criterion can become sub-optimal when the subset of removed features is large (*72*), we applied Recursive Feature Elimination (RFE) methodology. For each step of the procedure, we fitted the model and removed the features with smallest ranking criteria in a recursive manner until a certain number of features was reached.

The fundamental hyper-parameter of LR is the strength of the LASSO term tuned with a grid search procedure on the accuracy of the 10-fold cross-validation. The k-fold cross-validation provided the partition of the dataset into k batches, then exploited k^-1^ batches for the training and the remaining test batch as a test, by repeating this procedure k times. In the grid search method, a cross validation procedure was carried out for each value of the regularization hyperparameter in the range [10^-4^, …, 10^6^]. Specifically, the optimal regularization parameter is chosen by selecting the most parsimonious parameter whose cross-validation average accuracy falls in the range of the best one along with its standard deviation. During the fitting procedure, the class unbalancing was tackled by penalizing the mis-classification of minority class with a multiplicative factor inversely proportional to the class frequencies. For the RFE, the number of excluded features at each steps of the algorithm as well as the final number of features was fixed at 100. All data pre- processing and the RFE procedure was coded in Python; the LR model was used, as included, in the scikit-learn module with the liblinear coordinate descent optimization algorithm.

## List of Supplementary Figures

**Supplementary Figure S 1.** Homology Analyses.

**Supplementary Figure S 2.** Schematic Representation of the Classical STD Experiment and the Deconvolution Approach.

**Supplementary Figure S 3.** A Summary of the Workflow in a Classical NMR STD Experiment and Application of uSTA to Spike•Sugar-hybrid **5** and TreR•Tre systems.

**Supplementary Figure S 4 .** Modelling of Current STD Dependencies/Methods.

**Supplementary Figure S 5.** The uSTA Concept

**Supplementary Figure S 6 .** Workflow of uSTA.

**Supplementary Figure S 7 .** Preparation and Purification of SARS-CoV-2 Spike Protein- BAP.

**Supplementary Figure S 8 .** Advantages of the uSTA Pipeline, Emphasising Specific Advantages Over A More Conventional Analysis

**Supplementary Figure S 9 .** The Importance and Reliability of Ligand Subtraction when Calculating uSTA Surfaces [BSA•Trp Examples].

**Supplementary Figure S 10 .** The Importance and Reliability of Ligand Subtraction when Calculating uSTA Surfaces [SARS-CoV-2 Spike examples].

**Supplementary Figure S 11 .** uSTA Observes Stereochemical Discrimination in Binding even within Dominated Sugar Ligand Equilibria.

**Supplementary Figure S 12**. Synthetic Routes for Hybrid Sugars **5** and **6**.

**Supplementary Figure S 13 .** Preparation of SPR Chip and SPR Analysis.

**Supplementary Figure S 14 .** Combination of uSTA with HADDOCK Allowed Ranking of Docked Model Ensembles.

**Supplementary Figure S 15 .** uSTA Analysis of Spike from Variants of Concern.

**Supplementary Figure S 16 .** Effects Upon Sialoside Binding of the RBD-Blocking Neutralizing C5 Antibody

**Supplementary Figure S 17 .** Cryo-EM Coulombic Maps.

**Supplementary Figure S 18 .** Cryo-EM Confirms Additional Binding Mode for Aromatics in a Distinct Region of Spike.

**Supplementary Figure S 19 .** LASSO Regularization Profile of Clinical Data.

**Supplementary Figure S 20 .** Thermal Denaturation Analysis.

**Supplementary Figure S 21 .** Manual Assignment of Alpha Anomer of Neu5Ac.

**Supplementary Figure S 22 .** Manual Assignment of Alpha Anomer of 9-azido-Neu5Ac

## List of Supplementary Tables

**Supplementary Table S1**. COVID-19 Cohort

**Supplementary Table S2**. *B3GNT8* and *LGALS3BP* genetic variants

**Supplementary Table S3**. *B3GNT8* chi-square five categories

**Supplementary Table S4**. *B3GNT8* chi-square 2x2

**Supplementary Table S5**. *LGALS3BP* chi-square five categories

**Supplementary Table S6**. *LGALS3BP* chi-square 2x2

**Supplementary Table S7**. Raw data and UniDecNMR fits used to calculate the uSTA surfaces shown in the manuscript.

