## Supplementary material for "Cryptic pathogen-sugar interactions revealed by universal saturation transfer analysis": Supplmentary Notes, Figures and Tables

### SUPPLEMENTARY INFORMATION

#### Supplementary Notes

**Supplementary Note 1. Discussion of the Classical STD Experiment.**

Specifically, in a dynamic binding equilibrium, binding-rebinding cycles repeat during the saturation pulse. Although 'saturation' is not itself a measurable phenomenon, after a pre-set duration the remaining signal on the ligand can be recorded and the difference in signal for each resonance (sometimes called the 'saturation transfer difference') from a reference spectrum used to indicate ligand-protein contact. This magnetization transfer has often been colloquially described in terms of modes of 'saturation transfer' and, indeed, sometimes given an interpreted converse direction of 'saturation transfer' from protein-to-ligand[1] although, in practice, signal on the ligand is in fact reduced by this process.

**Supplementary Note 2. Discussion of the Utility of uSTA.**

'Universal' saturation transfer analysis (uSTA) only requires a series of specific simplified magnetization transfer spectra to be acquired on a protein-only sample, and then with a range of ligand concentrations (**Supplementary Figures S5, S6, S8**). The signal intensity from individual resonances in these spectra are obtained automatically. These are then converted into on and off rates (and $K_D$s) via complete theoretical treatment. These can also provide per resonance transfer efficiencies that, when used as constraints to high-level (e.g. HADDOCK) computational modelling environments provide exact structural models. In this way, uSTA analysis provides an automated pipeline from raw NMR free induction decay (FID) signals all the way to protein•ligand structures in a freely available form for the non-expert.

**Supplementary Note 3. Analysis and Circumvention of the Current Limits of Classical Methods of 'STD'.**

There are significant challenges that prevent classical 'STD' experiments being reliably applied to mammalian-derived proteins to quantitatively survey ligand binding (**Supplementary Figures S6,S8**). Our theoretical analyses (**Supplementary Figure S4**) suggested that many common assumptions or limits that are thought to govern the applicability of magnetization transfer might in fact be circumvented and we set out to devise a complete treatment that might accomplish this (**Supplementary Figures S5,S6,S8**). The result is a series of modifications to classical methods that combine to provide a more general and now quantitative, automated means of analyzing ligand-binding: uSTA (**Supplementary Figure S1**).

First, and perhaps most notably, appreciable transfer of signal can in fact occur for a wider range of timescales and strength of protein•ligand interaction than previously recognised (**Supplementary Figures S2,S4**). The sensitivity of such experiments is strictly independent of chemical exchange, in that the chemical shift difference between the free and bound ligand conformation is not in itself relevant to the mechanism of 'saturation'/magnetization transfer.

Instead, such modes are governed by the number of ligands that come into contact with protein and the cross relaxation between the two – this does not restrict the saturation transfer to the 'fast' exchange regime, as is commonly argued.[1] It also means that use of equations based on fast exchange that are typically used to analyze STD data cannot provide a general method. As a result this can lead to very larger errors in extracted $K_D$s, depending on the specific association and dissociation rates (**Supplementary Figures S6,8**).

Second, in seeking to precisely quantitate magnetization transfer, our analysis also reveals that it is vital to systematically vary both the protein and the ligand concentration in order to robustly separate exchange parameters from concentration-independent relaxation processes (**Supplementary Figure S4**).[1,2] In this way, forward and backward rates (and $K_D$) can be consistently obtained via experiments at multiple concentrations coupled with global, complete analyses of cross-relaxation.

Third, as in the case of heavily glycosylated human-derived proteins covered by highly mobile sugars,[3] the problems may be greatly exacerbated. The conformational flexibility of these groups is such that their $R_2$ relaxation rates become extremely short[3-5] leading to greatly confounded difference spectra. In conventional STD experiments, using for example the 'group epitope' method,[6] relaxation filters are added at the end of an experiment where protein signal is hoped to evenly decay away, ideally leaving only signal from ligands. However, in a protein that contains both mobile modifications as well as mobile disordered regions, such as SARS-CoV-2-spike, this approach is no longer viable since protein signal remains, even after aggressive use of such relaxation filters (**Supplementary Figures S5,S6,S8**). Moreover, relaxation filters inevitably result in reduced overall sensitivity and effects of intra-ligand cross-relaxation via the nuclear Overhauser effect (NOE) and ROE inhibit the elucidation of atom-specific data. In principle, however, relaxation filters would not be necessary at all with reliable extraction of ligand-only signal. We considered that baseline subtraction of residual signal transfer using reference to samples containing only protein could prove possible if methods for precise resonance identification could be developed.

Finally, NMR spectra from biological ligands such as sugars can be extremely complex, containing a large number of multiplets across diverse chemical shifts, each of which can be highly overlapped. Typically only the small number of resonances that are not overlapped are selected for detailed analysis in classical experiments, and so important information characterizing the ligand present in the spectrum cannot be easily extracted. In order to accurately quantify such a spectrum for use in saturation transfer methods, it is necessary to accurately determine the degree of magnetization (and its change) for each and all observed proton resonances. This too would be addressed by precise resonance identification (**Supplementary Figures S5,S6,S8**).

**Supplementary Note 4. Detailed Analysis of the Design of uSTA.**

STD analysis is a well-used method for studying protein/ligand interactions. It is widely thought to be a ‘fast exchange’ method, suitable only for weakly binding ligands with fast on/off rates. Quantitative analysis methods for STD data have been proposed based on the assumption that exchange is fast, though the agreement between measured $K_D$s from this method of analysis and alternative biophysical methods such as ITC and SPR can diverge by orders of magnitude.

We performed a theoretical analysis (**Supplementary Figure S4**) that suggests that many common assumptions or limits that are thought to govern the applicability of magnetization transfer might in fact be circumvented (see **Supplementary Note 3** for more details) and we set out to devise a more general experiment including pulse sequences as well as a data analysis pipeline that might accomplish this (**Supplementary Figures S5,S6,S8**). The result is 5 modifications that combine to provide a more general and now quantitative, semi-automated means of analyzing ligand-binding: uSTA (**Supplementary Figure S1**, **S5** for abstract, **S8** for improvements). Overall, the uSTA package provides a mechanism for semi-automatically analyzing protein/ligand interactions using NMR that is more general, sensitive and accurate than previously available methods.

First, in order to completely and quantitatively determine kinetic factors associated with binding ($k_{on}$, $k_{off}$, $K_D$) via uSTA we considered full aspects of the appropriate spin physics (**Figure 1** and **Supplementary Figure S2**). In brief, initially, the protein and ligand resonances are initially at equilibrium (**Figure 1A, grey**). Magnetization transfer experiments[7] move spin systems inside the protein that are within the bandwidth of the excitation ‘pulse ’out of equilibrium (**Figure 1A** and **Supplementary Figure S2**). These resonances then cross-relax with adjacent spins in the protein (spin-diffusion), which results in wide-ranging sets of spin systems within the protein that are effectively out of equilibrium. During this period, previously free ligand binds the protein and the ligand-protein cross relax via the nuclear Overhauser effect (magnetization is passed from ligand that is initially at equilibrium to out- of-equilibrium spin systems in the protein).

Modelling the degree to which signal has passed between the ligand and protein is complex; transfer depends on ligand binding kinetics $k_{on}$ and $k_{off}$ (and hence $K_D$) and the intrinsic cross-relaxation rates, which depend on the tumbling time of the complex (see **Figure 1A** and **Supplementary Figure S4** and **Methods** for mathematical analysis). A model for this signal transfer suitable for fitting to data has therefore not, to our knowledge, been addressed to date. However, these factors would, in principle, be straightforward to accommodate in theoretical analyses of the transfer, using comprehensive numerical approaches. In this approach, all relevant interactions could be fully and hence quantitatively described by modified Bloch-McConnell equations. Indeed, Bloch-McConnell approaches prove successful in other protein NMR methods involving dynamic processes such as CEST[8,9] or DEST.[10] Although it is more parametrically complex to analyze such magnetization transfer data using this formalism, all that would be required, in principle, is calculation[11] of the ratio of signal loss before and after the saturation pulse for each and every identified resonance, as well as the total concentration of protein and ligand (**Supplementary Figure S5, S6, S8**).

In this way, the theoretical description of the experiment would allow calculation of the expected transfer of signal with varying $k_{ex}$ and $K_D$ values (see **Methods**), where concentration dependent factors of biological significance, $k_{on}$, $K_D$, can be rigorously numerically disentangled from concentration independent factors such as relaxation rates. We demonstrated the effectiveness of the analysis by demonstrating that the BSA•Trp (**Figure 1**), and the SPIKE•sialoside $K_D$s (**Figure 3**) derived from NMR, and from ITC/SPR respectively are essentially identical. The uSTA analysis in addition to a $K_D$ provides 'true' in solution $k_{on}$, $k_{off}$ as well as an atomic resolution description of the ligand binding that allows us to determine binding poses of the ligands, and identify precisely which anomeric form of the sialosides (alpha or beta configurations of Sia) are binding with the protein.

This approach has similarity in principle to the CORCEMA method [12], where a protein structure is taken and all inter-proton distances are calculated to estimate how relaxation will evolve over the protein and between ligand and protein. As the matrices obtained for this calculation are extremely large, this is not an effective method for data analysis, as the calculation extremely expensive if used for optimization, and the rates obtained for cross relaxation in this way are known to be insufficiently accurate to describe the evolution of magnetization throughout a dynamic protein. For example, NOE intensities are not considered reliable indicators of distance in protein structure calculations for precisely the same reason. In our approach, we treat the system as both protein and ligand existing in 2 states, free and bound, which means we have effectively 4 spin states to consider, which can be represented as a 13 x 13 evolution matrix in the Bloch-McConnell equation. Here, the various relaxation rates are parameterized by relevant tumbling rates and geometric factors. This might appear to be a poor model for such a complicated system as a protein. Our confidence in the method comes primarily from our observation that the $K_D$s we obtain through fitting are consistent with those obtained using orthogonal methods. We rationalize its success from the fact that we are not seeking to get accurate descriptions of the intrinsic relaxation processes within the protein that are inherently complicated – we are instead trying to numerically separate the concentration dependent factors that tell us about binding interactions ($k_{ex}$, $K_D$) from the concentration-independent factors that will govern the relaxation in the system which we merely need to approximately parameterize.  Like when using 2-state models in analyzing protein dynamics via CEST/DEST/CPMG NMR experiments, our 2-state model here appears to strike an excellent balance between simplicity, and capturing the essential physics of the system in a manner that allows us to obtain useful biological data ($k_{on}$, $k_{off}$, $K_D$).

Second, central to this process is precise resonance identification and accurate extraction of signal intensity from all resonances. This was made possible by the design of a Bayesian computational method[13] to detect ligand and protein resonances in the 'raw' (unperturbed/reference) spectral data, even when complex and/or obscured by competing signals (see **Figure 1B** and **Supplementary Figures S5, S6, S8**). By this method, spectra could be automatically reduced to a series of constituent peaks and intensities (**Figure 1B,D**).[14,15] Resonances in the NMR spectrum can be described as a convolution between peak shape and a set of weighted delta functions (a 'delta matrix', see **Figure 1B** and **Methods**), and an algorithm developed previously for analyzing mass spectrometry data, Unidec [13] was used to iteratively determine the location of the 'true 'peaks.

The uSTA algorithm allows automatic detection of resonances returning a set of unique delta functions that describe peak positions and intensities, and a 'simulated' spectrum that allows a user to immediately visually verify the success of the calculation through comparison to the original 'raw' data. This process was executed simultaneously for two acquired spectra (excitation pulse on and 'off '(at -38ppm ppm, termed '1D' in figures)) in order to exactly and precisely determine not only peak positions, but also the change in intensity due to magnetization transfer (**Figure 1C**).

The effectiveness of the uSTA algorithm was measured via overlapped 1D spectra (see **Figures 1B, 1D, 3D, 4A** for examples – see also overlaps in all subsequent uSTA analyses and **Supplementary Table S7**) – the success of uSTA was immediately evident from comparison of raw data and algorithmically-derived spectra that were essentially identical in all cases. The peak positions identified by the algorithm mapped well to the locations established using conventional, manual NMR assignments (see **Methods**). Importantly, this enabled signal intensity determination in the background of other confounding resonances (**Supplementary Figure S5, S6, S8**). Moreover, those background resonances themselves could also be similarly analyzed. In this way, the two contributing components to signal intensity found in all magnetization transfer experiments (i.e. protein (P) and ligand (L)) were therefore determined and precisely dissected. Such precise determination of all contributions and consequent intensity changes upon magnetization transfer enabled a detailed quantitative analysis via uSTA by revealing signals that would be otherwise 'hidden '(**Figure 1**). The intensity from each multiplet was summed via scalar coupling, and the uSTA pipeline calculates transfer efficiencies on a per resonance, not a per multiplet basis.

Thirdly, we make adjustments to the experimental protocol that differ from the standard STD and 'epitope mapping' methodology [6]. In the case of mammalian proteins covered in highly flexible glycans, it is not possible to simply remove signals from protein using a 'relaxation filter'. A period in the pulse sequence where magnetization is held and resonances with a large $R_2$ from the protein decay to zero, aiming to leave only ligand resonances in the final spectrum. As these flexible residues with low $R_2$s are directly attached to the protein, they have a very strong STD response on their own. Moreover, during the 'relaxation filter', magnetization on the ligand is free to cross-relax internally thus making it more challenging to separate protein/ligand binding effects of biological interest from relaxation processes. In uSTA this is addressed simply by removing the relaxation filter, and recording a 'protein only' set of STD spectra that can be used by our software to baseline subtract. This builds on Saturation Transfer Double Difference (STDD) NMR [16], with uSTA enabling automatic deconvolution of the protein contribution (**Supplementary Figure S8**). The result is a simple addition to the method that generalizes its applicability and also increases its sensitivity by removing a segment of the pulse sequence where ligand signal unnecessarily decays.

Fourthly, we introduce a 'ligand subtraction'. When applying the excitation pulse during the saturation transfer process, the objective is to excite protein, but entirely 'miss' ligand. The bandwidth of the pulse ultimately sets the spectral region whereupon excitation is achieved, and so this ultimately sets the spectral 'distance', in ppms, that should separate ligand signals from the centre of the pulse. The action of a signal perfectly rectangular pulse element with well defined $B_1$ inhomogeneity has an excitation profile that resembles a sinc function, and so no matter where one excites, there is always a risk of small but transient excitation of regions of the ligand, that during a long excitation time can become significant, despite the fact that a simulation of the excitation profile of a 90˚ or 180˚ pulse might give the impression that there has been zero excitation outside the bandwidth of the pulse. We can effectively deal with this small residual excitation using a ligand subtraction (**Supplementary Figures 9 and 10**). We demonstrate that providing one excites in a position that is 2x the expected bandwidth of the pulse, and small residual effects of ligand excitation are subtracted, the final 'transfer efficiency' is invariant to precisely where the excitation is performed. This is physically sensible, and amounts to arguing that internal cross relaxation within the protein is much faster than protein/ligand cross relaxation. Without the ligand subtraction, the small effects of transient ligand excitation can compete with very small saturation transfer efficiencies in remote binding positions in the ligand, resulting in interaction/binding heat maps that are visually no longer identical. Thus, no matter which ppm is excited by an experimentalist, provided the ligand is avoided, the degree of cross relaxation from the ligand to the protein and hence the STD response is invariant to excitation frequency. When simulating the effects of the saturation transfer using the Bloch-McConnell equations, the STD response is highly sensitive to the details of the excitation pulse, and so it is vital to simulate exactly the pulse that was executed by the spectrometer (shape, phase, $B_1$ field amplitudes and duration).

Finally, we sum signal intensity from the different multiplets coming from a resonance to get true 'resonance' signal intensities, from which transfer efficiencies are calculated. These are represented as true atom specific measurements, and as a $<1/r^6>$ interpolated binding map 'that allows rapid and easy visual inspection of the ligand, that allows comparison of binding poses between sequences of ligands as we show in the text.

All this analysis is handled by our software, that is free to use for academic users. Bruker FIDs for the 'pulse on/pulse off' STD experiment are read-in, automatically processed to get spectra and analyzed. A peak list is provided, for which a user has to manually input an assignment (peak X is associated with resonance Y etc). This information is used to automatically generate the binding maps. If data is provided as a function of concentration, varying both protein and ligand concentrations, the software allows data fitting using the Bloch-McConnell formalism described to provide $k_{on}$, $k_{off}$, $K_D$ values

**Supplementary Note 5. Precision of uSTA in Allowing Dissection of Two Binding Modes in Unnatural Hybrid Sugars.**

In classical STD, the analysis of hybrid sugars would have been dominated simply by the most potent hydrophobic interaction (with ligand sugar protons lost within glycoprotein sugar responses). In this way, we could demonstrate that such modified ligands bind through *all* portions of their surfaces but a distinct difference of interaction is observed in hybrid ligands: a greater and distinct contact was seen with the hydrophobic moiety than with the carbohydrate moiety. Such is the precision of uSTA, graded binding even within these two portions of the ligand was also determined (see atom-specific scoring). For example in **5**, greatest binding for the hydrophobic moiety was seen at the tip with lowering, graduated binding towards the amide junction at C9. In the sugar portion, despite a quite different contact type, individual gradation was also observed: the C7-C9 sidechain bound most strongly but a clear significant contribution from the NHAc-5 group can also be discerned. Trisaccharide-BPC hybrid sugar **6** (with both hydrophobic BPC-moiety and sugar) was also synthetically generated (**Supplementary Figure S12**) to test the relative dominance of the two most potent moieties identified by uSTA. Interestingly, significantly extended binding interfaces were seen in both moieties, consistent with two modes of binding. In this way, uSTA rapidly allowed the mapping and iterative design of natural and unnatural sugar ligands for SARS-CoV-2-spike and the identification of multiple potential binding modes present in these unnatural variants.

**Supplementary Figures**

.

**Supplementary Figure S 1.** Homology Analyses.
**Supplementary Figure S 2.** Schematic Representation of the Classical STD Experiment and the Deconvolution Approach.
**Supplementary Figure S 3.** A Summary of the Workflow in a Classical NMR STD Experiment and Application of uSTA to Spike•Sugar-hybrid **5** and TreR•Tre systems.
**Supplementary Figure S 4 .** Modelling of Current STD Dependencies/Methods.
**Supplementary Figure S 5.** The uSTA Concept
**Supplementary Figure S 6 .** Workflow of uSTA.
**Supplementary Figure S 7 .** Preparation and Purification of SARS-CoV-2 Spike Protein-BAP.
**Supplementary Figure S 8 .** Advantages of the uSTA Pipeline, Emphasising Specific Advantages Over A More Conventional Analysis
**Supplementary Figure S 9 .** The Importance and Reliability of Ligand Subtraction when Calculating uSTA Surfaces [BSA•Trp Examples].
**Supplementary Figure S 10 .** The Importance and Reliability of Ligand Subtraction when Calculating uSTA Surfaces [SARS-CoV-2 Spike examples].
**Supplementary Figure S 11 .** uSTA Observes Stereochemical Discrimination in Binding even within Dominated Sugar Ligand Equilibria.
**Supplementary Figure S 12**. Synthetic Routes for Hybrid Sugars **5** and **6**.
**Supplementary Figure S 13 .** Preparation of SPR Chip and SPR Analysis.
**Supplementary Figure S 14 .** Combination of uSTA with HADDOCK Allowed Ranking of Docked Model Ensembles.
**Supplementary Figure S 15 .** uSTA Analysis of Spike from Variants of Concern.
**Supplementary Figure S 16 .** Effects Upon Sialoside Binding of the RBD-Blocking Neutralizing C5 Antibody
**Supplementary Figure S 17 .** Cryo-EM Coulombic Maps.
**Supplementary Figure S 18 .** Cryo-EM Confirms Additional Binding Mode for Aromatics in a Distinct Region of Spike.
**Supplementary Figure S 19 .** LASSO Regularization Profile of Clinical Data.
**Supplementary Figure S 20 .** Thermal Denaturation Analysis.
**Supplementary Figure S 21 .** Manual Assignment of Alpha Anomer of Neu5Ac.
**Supplementary Figure S 22 .** Manual Assignment of Alpha Anomer of 9-azido-Neu5Ac

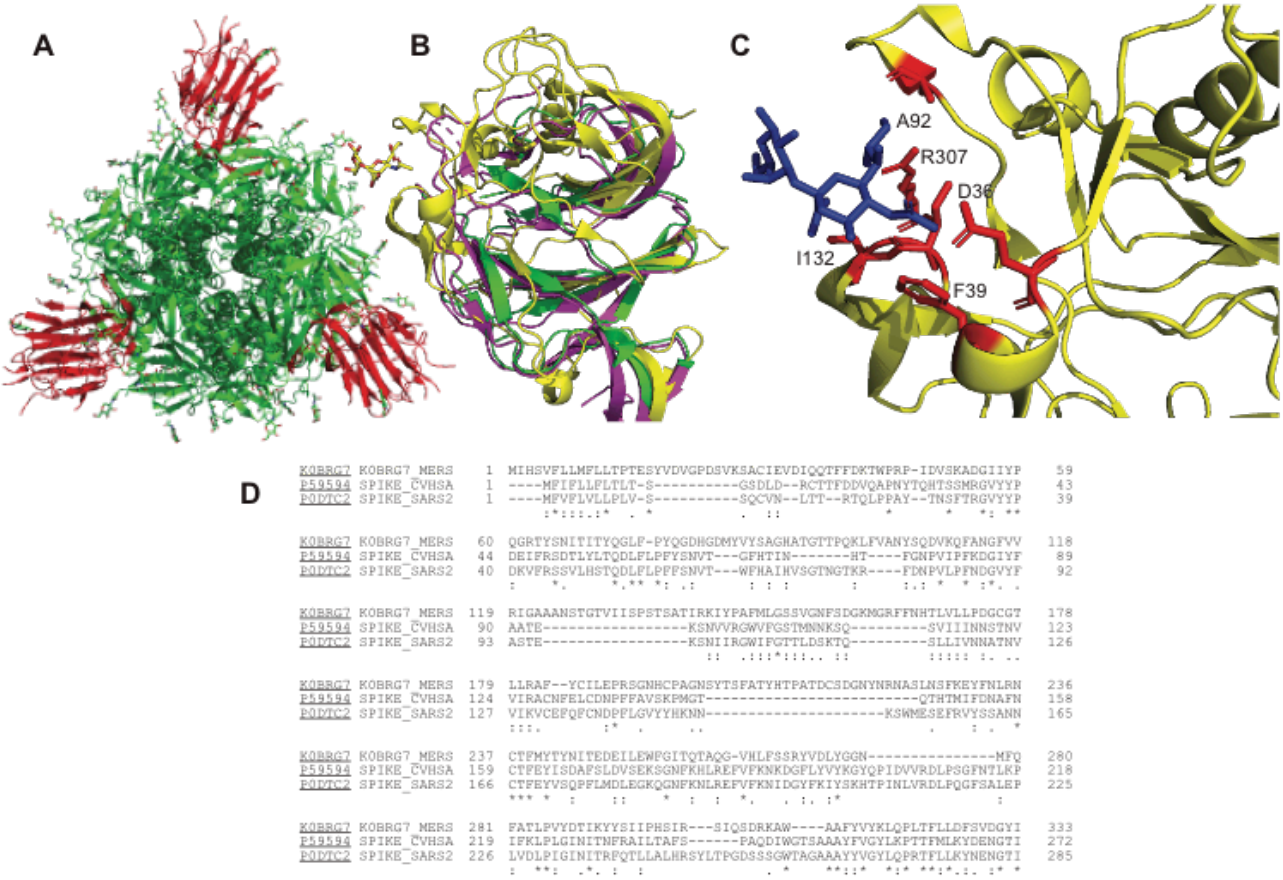

**Supplementary Figure S1. Homology Analyses. A**) The N-terminal domain of the SARS-CoV-2 spike protein (green, 6vxx). The N-terminal domain comprises the first 280 residues (red). **B**) The structural homology of MERS (yellow, 6q06), sars-cov1 (magenta, 5x58) and sars-cov2 (green, 6vxx) spike proteins in this region is partial. The C-terminal domains show substantial similarities, but the N-terminus show substantial differences. **C**) The interaction between Gal-Sia and MERS NTD. The interaction can be largely attributed to 5 amino acid residues with the NAc binding F39, I132 and D36, and the glycerol arm binding to A92 and R307. D) Sequence homology of the N-terminal domains of MERS, SARS-CoV-1 and SARS-CoV-2 using BLASTP. Substantial differences between the three proteins are observed. The five key residues for sialic acid binding in MERS are indicated. These all fall within regions in SARS-CoV-1 and SARS-CoV-2 that differ substantially from MERS, thus rendering the likelihood of generating a reliable homology model exceedingly unlikely.

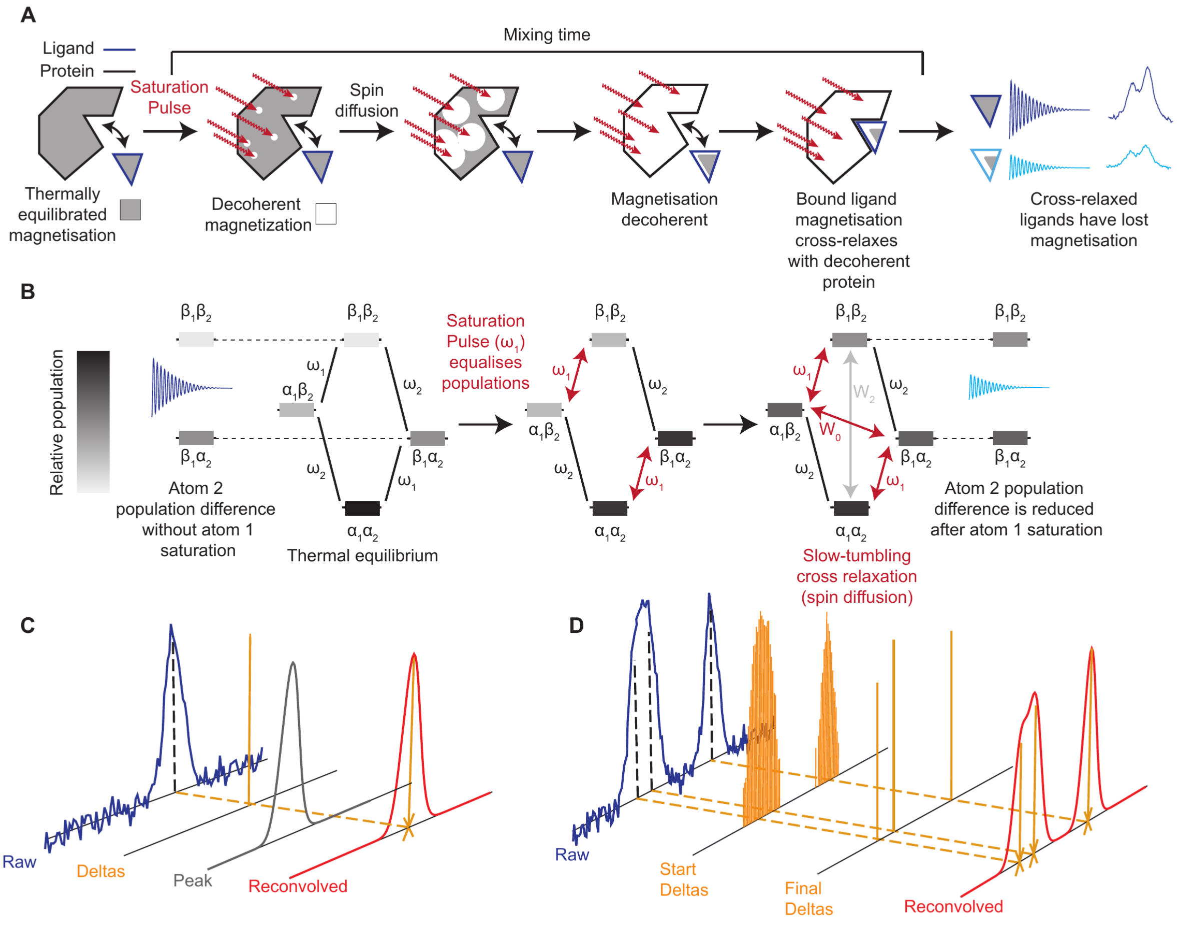

**Supplementary Figure S2: Schematic Representation of the Classical STD Experiment and the Deconvolution Approach**.

**A**): A 'saturation pulse' is applied to a protein and ligand mixture that covers a specific but narrow range of ppm frequencies selected to avoid excitation of the ligand. The nuclei excited in the protein will have specific spatial locations within the protein, for example methyl groups or amides. Spins in the protein that have been excited will cross-relax with their neighbours, aiming to restoring the initially excited spins to equilibrium. This process continues during the period in which the excitation pulse is applied and the net magnetisation on each spin within the protein is gradually reduced to zero, a process termed ‘spin diffusion’. The rate of the many interior cross relaxations increases as tumbling time of the protein decreases. When a ligand binds rigidly to the protein, intermolecular cross-relaxation can occur if adjacent spins in the protein have been knocked out of equilibrium by either the initial excitation, or spin diffusion. The action will be to transfer unperturbed longitudinal magnetisation from the ligand to partially restore the now unequilibrated longitudinal magnetization on the various spins of the protein. After the mixing time, the quantity of magnetization on the ligand is detected, following a 90˚ pulse. The action of the saturation pulse therefore is to reduce the magnetisation on the ligand in a manner that depends on the on and off rates, the various tumbling times of ligand and protein that govern intrinsic ($R_1/R_2$) and cross (intra/inter) relaxation and the details of the saturation pulse. A 'build-up curve' follows the amount of lost signal from the ligand versus the duration of the mixing time. These processes can be effectively modelled using modified Bloch-McConnell equations and by recording 'build-up curves' at a range of varying ligand and protein concentrations, and fitting them to an appropriate model based on the modified Bloch-McConnell equations (see methods) concentration independent

effects such as relaxation rates, and concentration dependent effects, such as binding, can be separated, as is demonstrated in this article.
**B**): The details of the experiment can be understood with a series of single spin energy level illustration. Initially, protein and ligand spins are thermally equilibrated. The application of a saturation pulse at ω1 (always chosen to be a protein spin) induces coherence between the a and b spin states, and with sustained application will eventually equalises populations in the two α1 to β1 transitions (the transition becomes 'saturated', net magnetisation for the spin in the ensemble has gone to zero). In our slowly-tumbling protein, the W0 cross-relaxation pathway is most efficient, providing a path to move magnetization from the state closes to equilibrium (ligand) to the state furthest from equilibrium (protein), here pushing population from the β1α2 state to the α1β2. The combination of continuous application of saturation pulse and cross-relaxation (both intra to ligand and inter within the protein) lowers the population differences between the interacting spin states. This is the nuclear Overhauser effect (NOE). The rate for cross relaxation, both inter and intra depends on the tumbling time of the complex, and the distance between the spins. If atom 2 is a nearby protein spin, we are interrogating part of intra-protein cross relaxation or spin diffusion, if atom 2 is a bound ligand spin, we are interrogating intermolecular cross-relaxation. These complex processes can be well accounted for in a simple two spin model as described in the methods. Our analysis is validated in this work by demonstrating that fitted $K_D$s using our picture returns values that are equal to those obtained by independent means, together with the on and off rates.
**C),D):** Schematic of process for automatically quantifying the signal intensities in NMR spectra using deconvolution. The analysis determined the number of peaks that can give rise to the signal and return a simulated spectrum by convolving these with a peak shape function. Precise peak positions and their intensities are returned, which can be attributed to single protons observed in a spectrum.

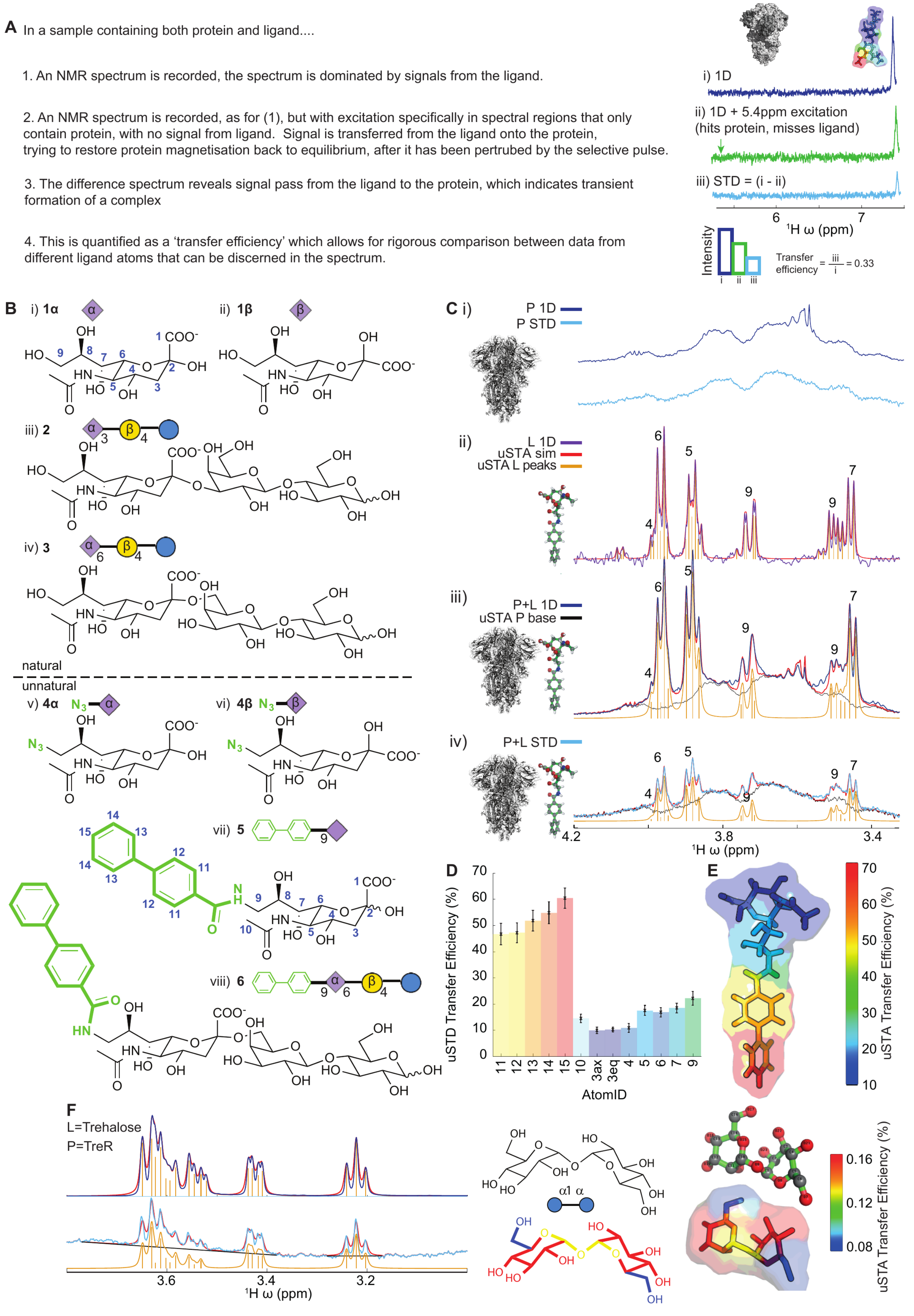
A In a sample containing both protein and ligand....
1. An NMR spectrum is recorded, the spectrum is dominated by signals from the ligand.
2. An NMR spectrum is recorded, as for (1), but with excitation specifically in spectral regions that only contain protein, with no signal from ligand. Signal is transferred from the ligand onto the protein, trying to restore protein magnetisation back to equilibrium, after it has been pertrubed by the selective pulse.
3. The difference spectrum reveals signal pass from the ligand to the protein, which indicates transient formation of a complex
4. This is quantified as a 'transfer efficiency' which allows for rigorous comparison between data from different ligand atoms that can be discerned in the spectrum.
i) 1D
ii) 1D + 5.4ppm excitation (hits protein, misses ligand)
iii) STD = (i - ii)
1H ω (ppm)
Intensity
Transfer efficiency = iii / i = 0.33
B i) 1α
ii) 1β
iii) 2
iv) 3
natural
unnatural
v) 4α
vi) 4β
vii) 5
viii) 6
C i)
P 1D
P STD
ii)
L 1D
uSTA sim
uSTA L peaks
iii)
P+L 1D
uSTA P base
iv)
P+L STD
1H ω (ppm)
D
uSTD Transfer Efficiency (%)
AtomID
E
uSTA Transfer Efficiency (%)
F
L=Trehalose
P=TreR
1H ω (ppm)
α1 α
uSTA Transfer Efficiency (%)

**Supplementary Figure S3. A Summary of the Workflow in a Classical NMR STD Experiment and Application of uSTA to Spike•Sugar-hybrid 5 and TreR•Tre systems.**

**A):** When the protein is selectively excited, its magnetisation is pushed far from its equilibrium value. When a ligand binds the protein when in this state, cross relaxation occurs where signal from the ligand moves onto the protein, effectively trying to restore the protein to equilibrium. By comparing the intensity of ligand resonances with, and without the selective excitation, the quantity of magnetisation that has been transferred can be determined, which indicates binding. In order to compare accurately between different atoms in the same spectrum, the degree of transferred magnetisation needs to be expressed relative to the initial magnetisation. This ratio is referred to in this work as the 'transfer efficiency', which can be physically interpreted as a fraction of magnetisation from an atom on the ligand that has moved onto the protein. Increasing the power and the duration of the selective excitation will increase this signal. The specific transfer efficiency will depend on the proximity of the ligand atom and the protein when the complex forms, the overall correlation time of the complex, and the chemical kinetics of the complex formation, the on and the off rate. With the uSTA method, these factors are all treated rigorously using the Bloch-McConnell equations in order to ascertain a reliable $K_D$.

**B):** A panel of natural, unnatural and hybrid variant sialoside sugars **1-6** was used to probe interaction between sialic acid moieties and spike. Unnatural variations (**4-6**, green) allowed mapping of C7-C9 side-chain interactions in the sialoside whilst use of extended sugars probed differing cell-surface glycan structures

**C):** Application of the uSTA workflow (**Supplementary Figure S6**) to SARS-CoV-2-spike protein (shown in detail for **5**, see also **Figures 1,2**) i) The 1D $^1$H-NMR of SARS-CoV-2-spike protein shows considerable signal in the glycan-associated region despite protein size, indicative of mobile internal glycans in spike protein. This effectively masks traditional analyses, as without careful subtraction of the protein's contributions to the spectrum (**Supplementary Figure S8**), the ligand cannot be effectively studied. The uSTA process of: ii) ligand peak assignment and deconvolution → iii) p + L peak assignment and deconvolution → iv) application to p + L STD yields precise atom-specific transfer efficiencies (**Supplementary Figure S6**). Note how in ii) individual multiplet components, have been assigned (yellow); the back-calculated deconvolved spectrum (red) is an extremely close match for the raw data (purple). In iii) the spectrum is a complex superposition of the ligand spectrum (and protein only yet uSTA again accurately deconvolves the spectrum revealing the contribution of protein only (black) and the ligand peaks (yellow). Using these data, uSTA analysis of the STD spectrum in iv) pinpoints ligand peaks and signal intensities.

**D),E):** Using these intensities, atom-specific transfer efficiencies can be determined with high precision and reveal in hybrid **5** the details of both the unnatural BPC moiety and the natural sialicacid moiety. Although the aromatic BPC dominates the interaction for the unnatural ligand **5**, the subtleties of the associated sugar contribution in this ligand can nonetheless be determined (**Supplementary Figure S5,S6,S8**).

**F):** Application of the same uSTA workflow allows precise determination of even weakly binding sugar ligand trehalose (Glc-α1,1α-Glc) to E. coli trehalose repressor TreR. Again, the uSTA allows determination of transfer efficiencies with atom-specific precision.

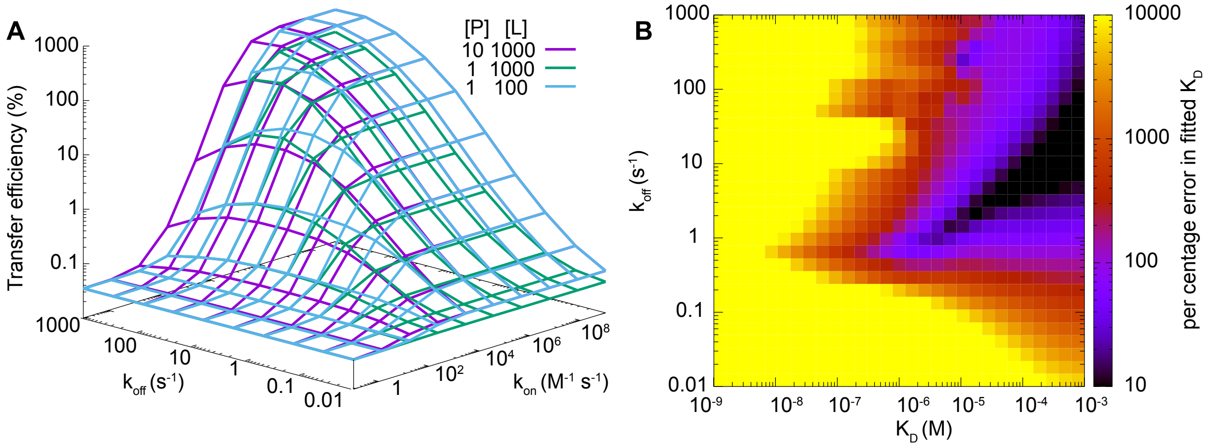

**Supplementary Figure S4. Modelling of Current STD Dependencies/Methods. A)** The region of validity of the saturation transfer experiment is wider than has been realised, and that by taking both variable protein and ligand concentrations, the two rates, $k_{on}$ and $k_{off}$, can be numerically separated from the concentration independent factors that describe the relaxation processes. Transfer efficiency from the saturation transfer experiment was simulated using the Bloch-McConnell equations (see **Methods**) using parameters optimised for the alpha3/spike interaction that describe the concentration independent relaxation effects ($t_G$ 2.5 ns, $t_E$ 85 ns, $r_{IS}P$ 1.6 A, $r_{IS}l$=3.1 A, $r_{IS}$(mix) 2.4 A, *fac* 18) simulated using a Gaussian pulse train of peak $B_1$ field of 200 Hz with each element running for 50 ms for a total duration of 5 s, as utilised in the experiments, as a function of $k_{on}$ and $k_{off}$ for various protein and ligand concentrations (specified on figure, units of μM). The specific value of the parameters that affect relaxation alter the expected transfer efficiency, but fitting data with varying protein and ligand concentration is able to uniquely determine on and off rates. Experimentally, the ability to measure a transfer efficiency depends on the specific signal to noise, which can always be increased with more scans. In this work, we reliably determine transfer efficiencies on the order of 0.5%. At this sensitivity level, $k_{on}$ needs to exceed 100 $M^{-1}$ $s^{-1}$, and $k_{off}$ needs to exceed 0.1 $s^{-1}$. Above these limits, the transfer efficiency is increases with increasing on and off rates. Above these limits, suitable combinations of ligand and protein concentration can give sufficiently different values of transfer efficiency at different mixing times to enable the concentration independent intrinsic relaxation rates to be reliably separated from the concentration dependent parameters that describe the protein/ligand interaction, $k_{on}$ and $k_{off}$ during numerical fitting. Notably, we obtain two physically sensible limits from these simulations. With high $k_{off}$, the transfer efficiency becomes independent of ligand concentration (green and blue converge), whereas at high $k_{on}$, the transfer efficiency becomes independent of the overall protein to ligand ratio, L/P (purple and blue converge). In a general case, determining errors in the fitting parameters is sufficient to tell a user whether or not they are well defined. If insufficient ligand/protein concentration dependence has been sampled to accurately determine $k_{on}$ and $k_{off}$, the numerical uncertainties in these parameters obtained from a bootstrapping procedure will be large.

**B)** It has been previously proposed that an approximate treatment can be applied to reliably obtain $K_D$ values.[1,17] Applications of this method have resulted in $K_D$ values being significantly underestimated, by orders of magnitude or more, suggesting limitations in this approach.[18] By using this approach to fit simulated data, we establish that this method has a very narrow region where it can be considered effective. As has been argued previously,[17] a Bloch-McConnell treatment that covers the chemical kinetics and the relaxation processes, exactly as we perform in this work, is the method by which saturation transfer data should be analysed, and with appropriate choices of protein and ligand concentrations, is expected to yield accurate exchange parameters for the range of on and off rates described in (A). In brief, the method described by Angelo et al. determines the initial gradient of the build-up curve. These values are then followed versus ligand concentration to create an isotherm, which is fitted to $A[L]/(K_D+[L])$ to obtain $K_D$ values. Here, we perform this analysis by simulating isotherms with 10 ligand concentrations spaced between 10 μM and 1 mM for a wide range of pairs of $K_D$ and $k_{off}$. The simulated data is fitted according to the protocol described above, and the per centage error in the fitted $K_D$ versus the simulated $K_D$ is obtained. This fitting method is shown to be effective only in a narrow window of parameter space (black, errors below 10%). These points are roughly described by a locus centred on 50 μM $K_D$ and an off rate of 10 $s^{-1}$. In the cases encountered in this work, the off rate is determined to be in the vicinity of 0.1 $s^{-1}$, a region of parameter space where the expectation for reliable parameters using this method is expected to give, at a minimum errors of 1,000 %. Fundamentally, the assumptions that underpin this method requires that the saturation transfer is proportional to the percentage of bound complex, and the proportionality constant is itself independent of protein/ligand combination. This is reliable only for a narrow region of parameter space. The differences shown here, between 'actual' and 'fitted' $K_D$ demonstrate that 'non-specific binding' is not required to explain the discrepancy.[18] The fitting methods described in this work using the Bloch-McConnell equations do not need to make assumptions about the binding regime and provide a reliable and rigorous method to obtain parameters such as $k_{on}$, $k_{off}$ and $K_D$.

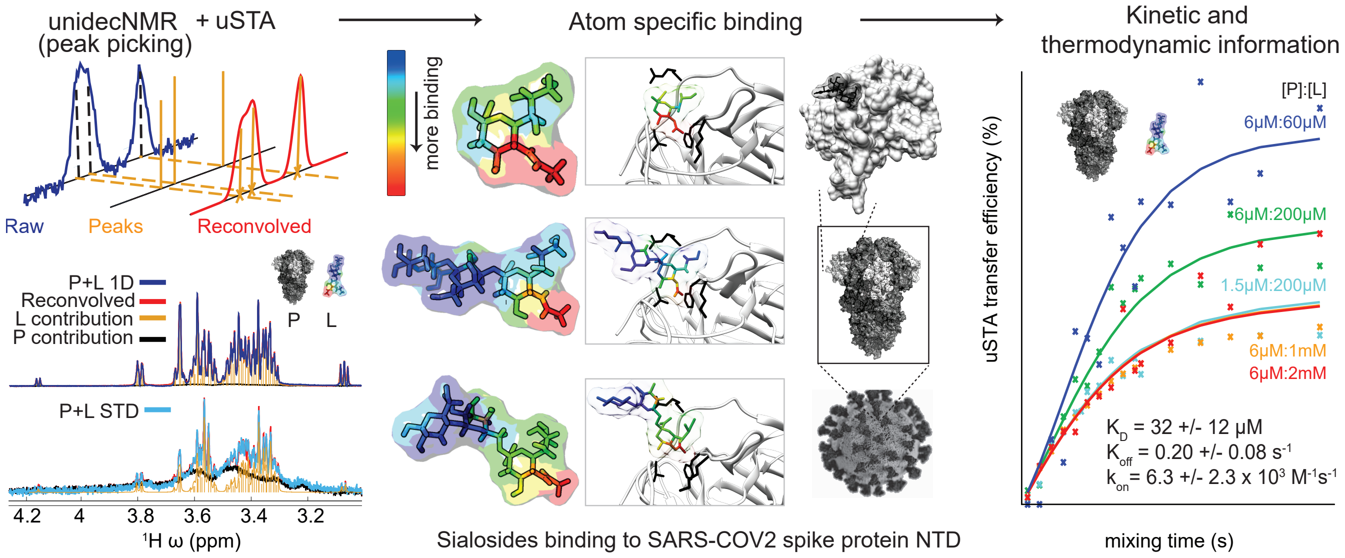

**Supplementary Figure S5. The uSTA Concept.**

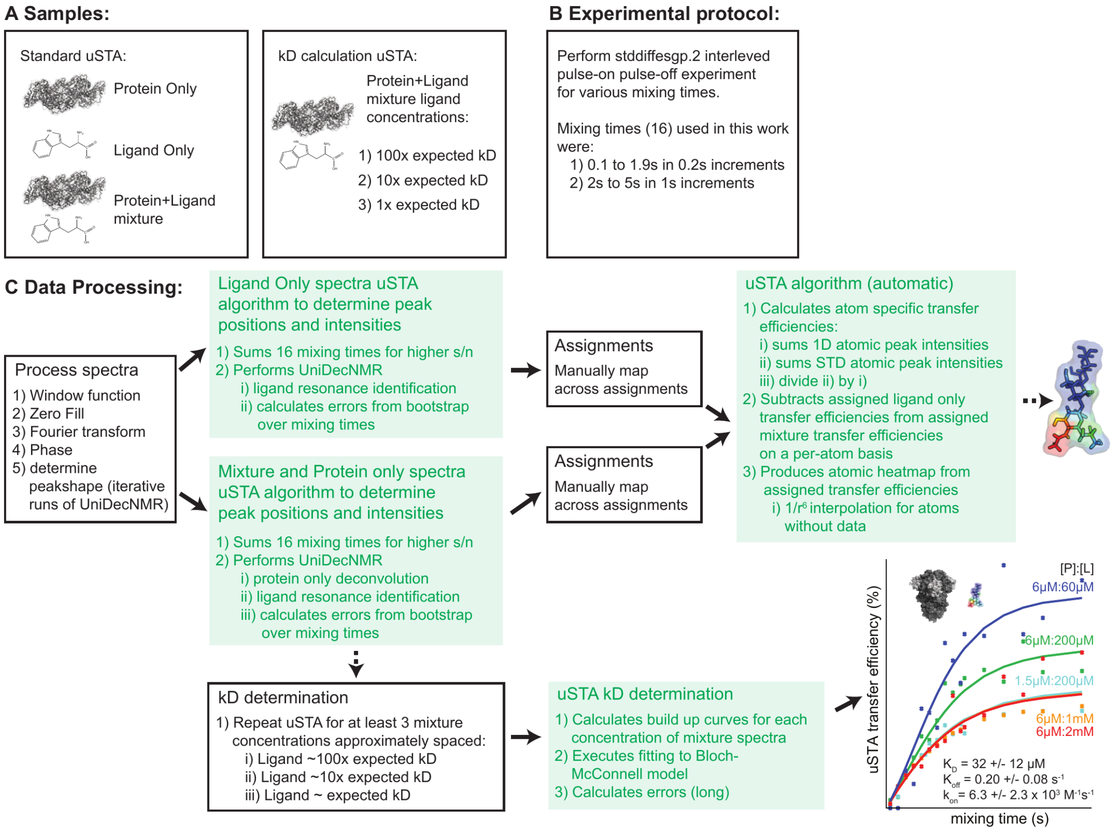

**Supplementary Figure S6. Workflow of uSTA.** Flowchart describing the specific samples required for a uSTA analysis, and the data processing steps involved. The boxes with a white inlay are the parts that require manual intervention, and the boxes in green show the automated analysis by the uSTA pipeline including figure generation.

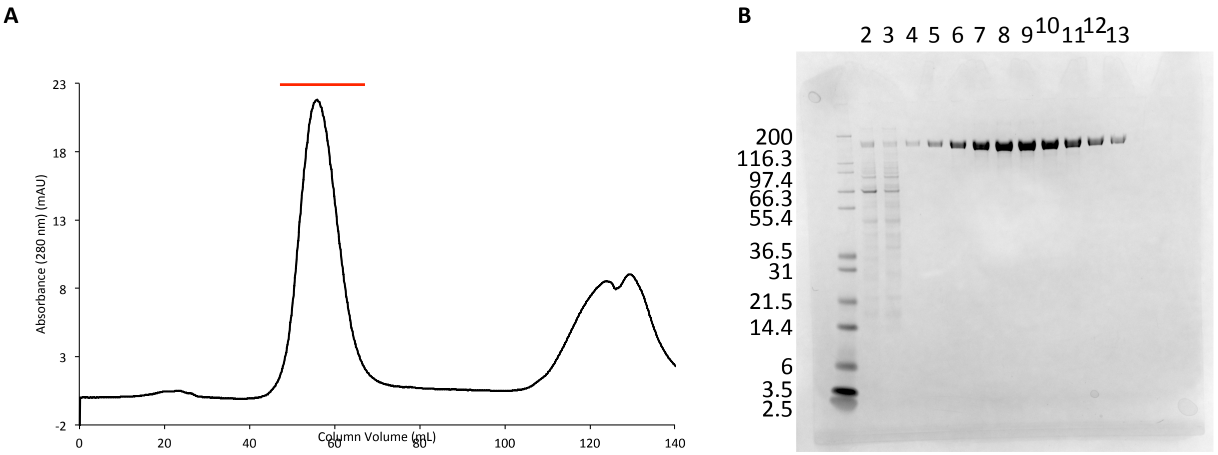

**Supplementary Figure S7.** Preparation and Purification of SARS-CoV-2 Spike Protein-BAP. **A**: Gel filtration (Superdex S200, GE Life Sciences) profile of Spike-BAP following elution from Ni resin. Elution volume 56.7 mL. Red line denoted fractions analyzed on SDS-PAGE. **B**: SDS-PAGE analysis of Spike-BAP purification. 1 - Mark 12 ladder (labels in kDa); 2 - Load onto Ni column; 3 - Flow through from Ni; 4 - Gel filtration fraction A3; 5 - Gel filtration fraction A4; 6 - Gel filtration fraction A5; 7 - Gel filtration fraction A6; 8 - Gel filtration fraction A7; 9 - Gel filtration fraction A8; 10 - Gel filtration fraction A9; 11 - Gel filtration fraction A10; 12 - Gel filtration fraction A11; 13 - Gel filtration fraction A12. Fractions A3 to A12 were pooled and concentrated to a final volume of 1.6 mL at 7.91 μM.

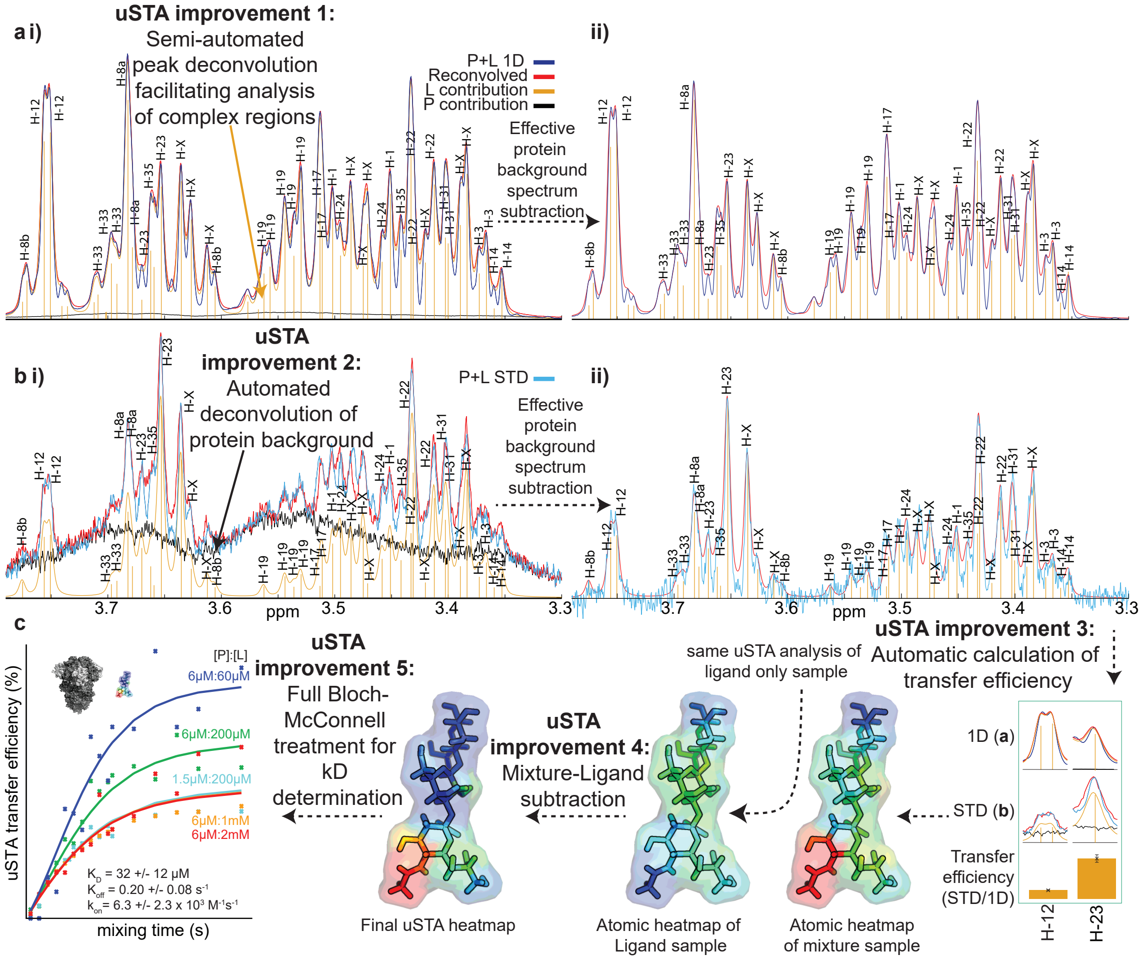

**Supplementary Figure S8. Advantages of the uSTA Pipeline, Emphasising Specific Advantages Over A More Conventional Analysis.** The UnidecNMR algorithm allows for highly efficient and automated peak detection, including the ability to subtract the protein background. The analysis when combined with a manual assignment of the individual resonances directly computes the transfer efficiencies which when combined with ligand-only data, provide a heat map of the molecule of interest. Finally, where data are available with variable protein/ligand concentrations, the method computes a $K_D$ via a model using the Bloch-McConnell equations.

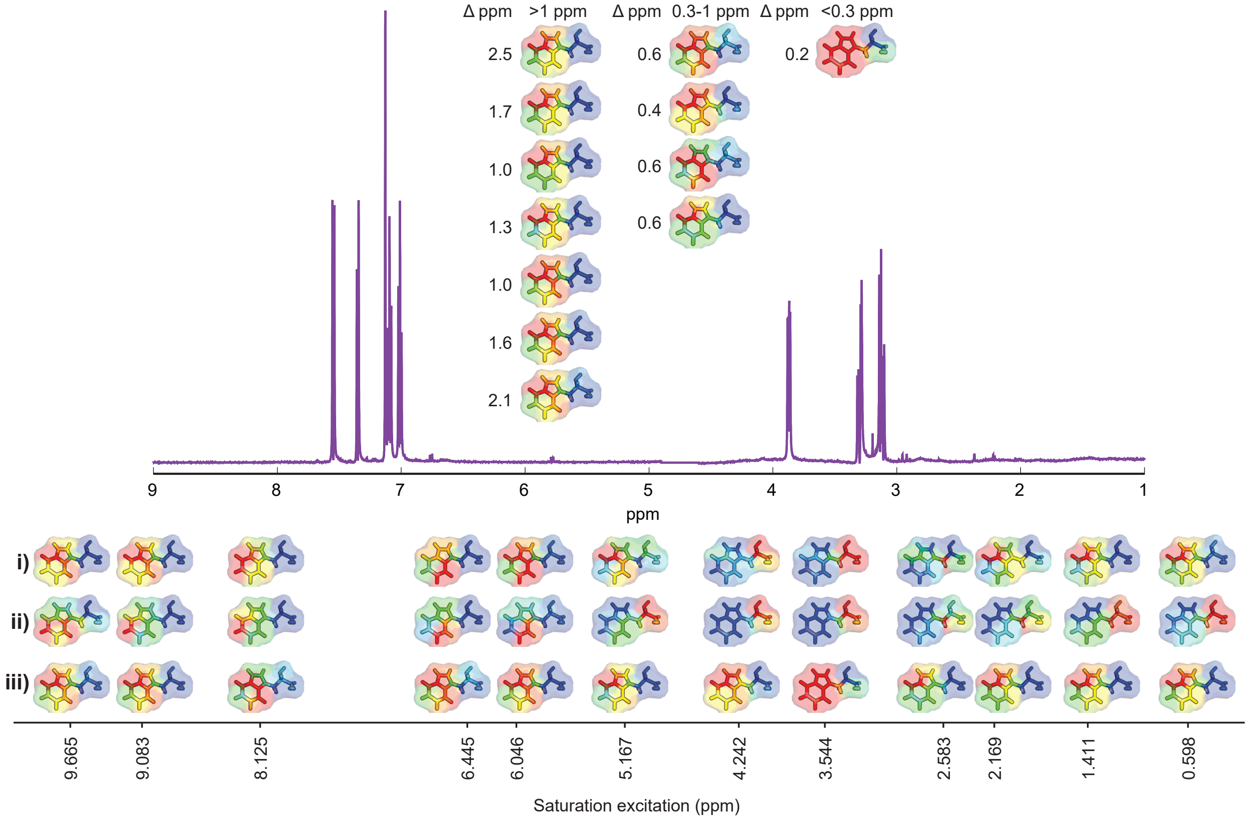

**Supplementary Figure S9. The Importance and Reliability of Ligand Subtraction when Calculating uSTA Surfaces [BSA•Trp Examples].** The interaction between BSA and tryptophan was examined while varying the excitation frequency of the saturation pulse. The 'apparent' uSTA surface was found to vary slightly depending on precisely where the excitation pulse was applied **(i)**. However, an analogous result was observed for a sample containing purely ligand **(ii)**. Subtracting the ligand-only contributions produced highly conistent heatmaps **(iii)** when the excitation was further than 1 ppm from a ligand resonance. The excitation bandwidth of the Gaussian pulse cascade with a $B_1$ field of 200 Hz should be ca. 0.3ppm at 600MHz. When the excitation pulse comes within 0.6 ppm of a ligand resonance however, excitation effects can be clearly observed in a pure ligand sample. In the high concentration limit, the transfer efficiency tends to a constant, and so to a reasonable first approximation we can correct the measurement by subtracting the ligand value. Thus when we excite > 1ppm from the ligand, although in some cases there is variation in the 'apparent' uSTA surface deriving from the P+L sample, the uSTA surfaces we obtain following ligand subtraction are nevertheless highly similar. In cases where the excitation is within 0.3 ppm of a peak centre, the ligand subtractions are large, though the final surface still captures the major features. It is desirable to pulse > 1ppm from a ligand peak. We might expect the uSTA surface to vary with excitation frequency, as different residues in the protein will be excited. These results suggest that the cross relaxation within the protein is highly efficient, and providing the ligand only results are subtracted from the mixture, the final surface is largely invariant of where we excite.

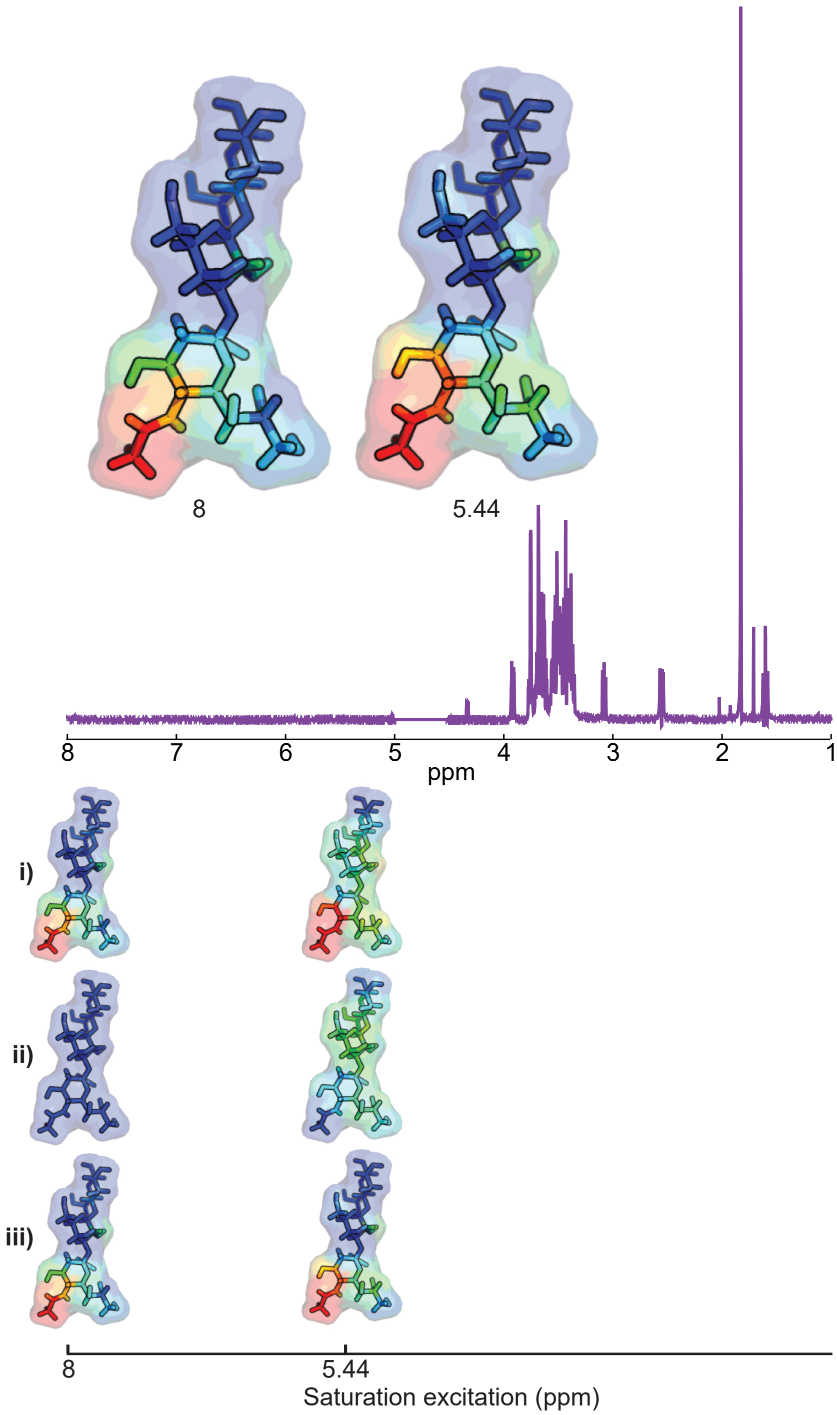

**Supplementary Figure S10. The Importance and Reliability of Ligand Subtraction when Calculating uSTA Surfaces [SARS-CoV-2 Spike examples].** The variation of the uSTA surface depending on excitation frequency. uSTA surfaces for alpha3 were calculated before **(i)** and after **(iii)** subtracting the ligand-only contribution **(ii)**, exciting at 8 ppm (2.7 ppm from a ligand resonance) and at 5.4 ppm (0.3 ppm form a ligand resonance). After subtracting the ligand contributions, the final surfaces are highly similar, a result mirroring our findings for BSA and tryptophan (**Supplementary Figure S9**).

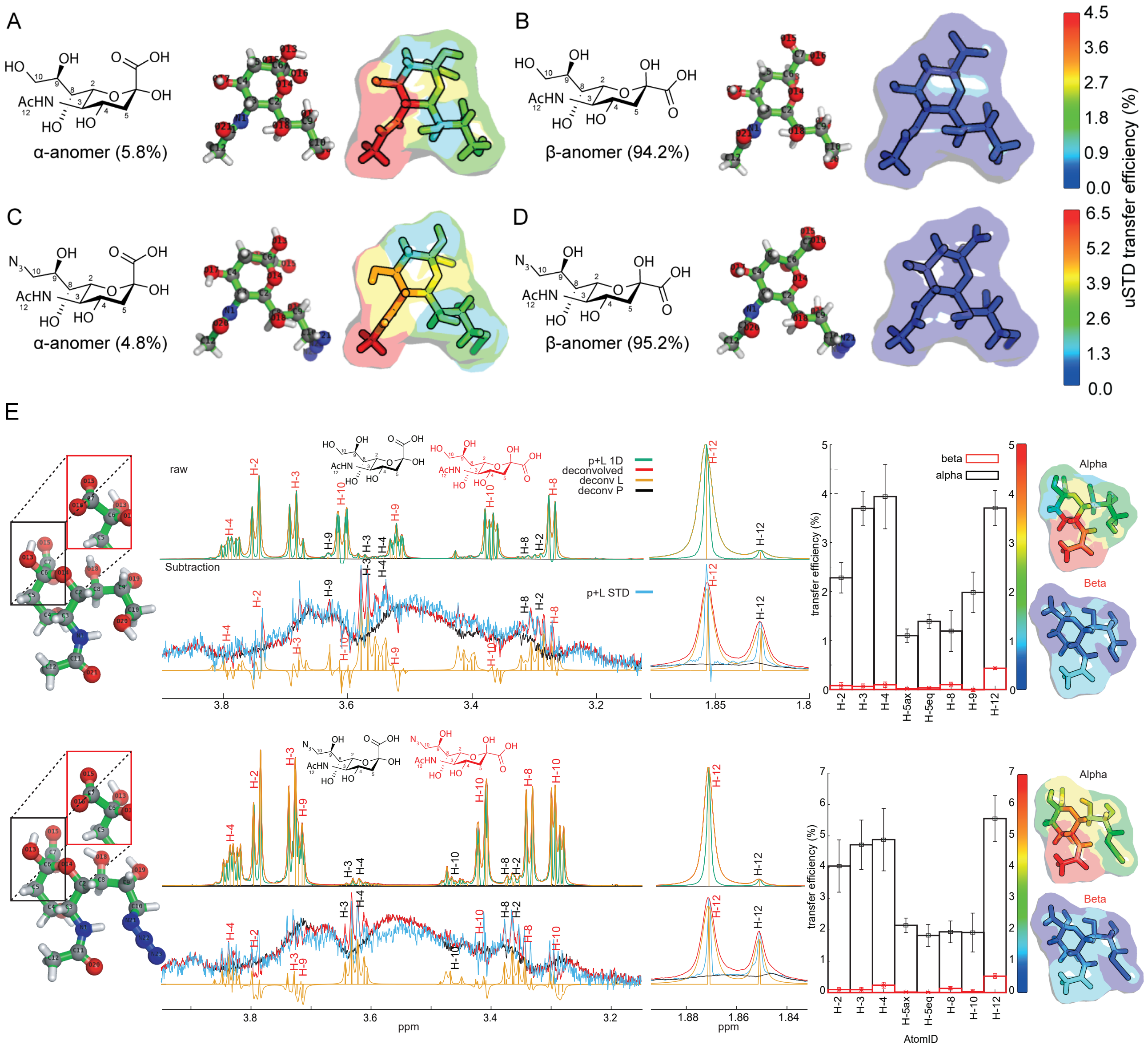

**Supplementary Figure S11. uSTA Observes Stereochemical Discrimination in Binding even within Dominated Sugar Ligand Equilibria.**
In the spectra of sialic acid (**1**) and azido-sialic acid (**4**), both α and β anomeric forms could be readily identified, with the overall population being dominated by the β form (94 and 95%, respectively). Even despite this strong population difference, application of the uSTA method following assignment of resonances from the two forms allowed determination of binding surfaces simultaneously. Spike shows strong binding preference for the α anomers, as revealed by surfaces (**A**, **C**) even though its population is minor overall; both surfaces were highly similar for these two simpler monosaccharides but closely resemble those of extended trisaccharides **2** and **3** (Figure **3**). While the β form is dominant in terms of population and overall contribution to the 1D NMR spectra, its ability to bind spike (**B**, **D**), and hence its proportional contribution to the difference spectrum, was found to be significantly lower than the α form.
(**E**) For α and β anomers of **1** and **4**. While the 1D spectrum is dominated by the β form of both ligands, the STD spectrum is dominated by the α form. The uSTA surface shows a similar pattern for these, and ligands **2** and **3**, where the NAc from the methyl group and adjacent protons interact most strongly with the protein. Given the low signal intensities for the resonances from the β form, we cannot conclusive demonstrate any binding of the β form from these data.

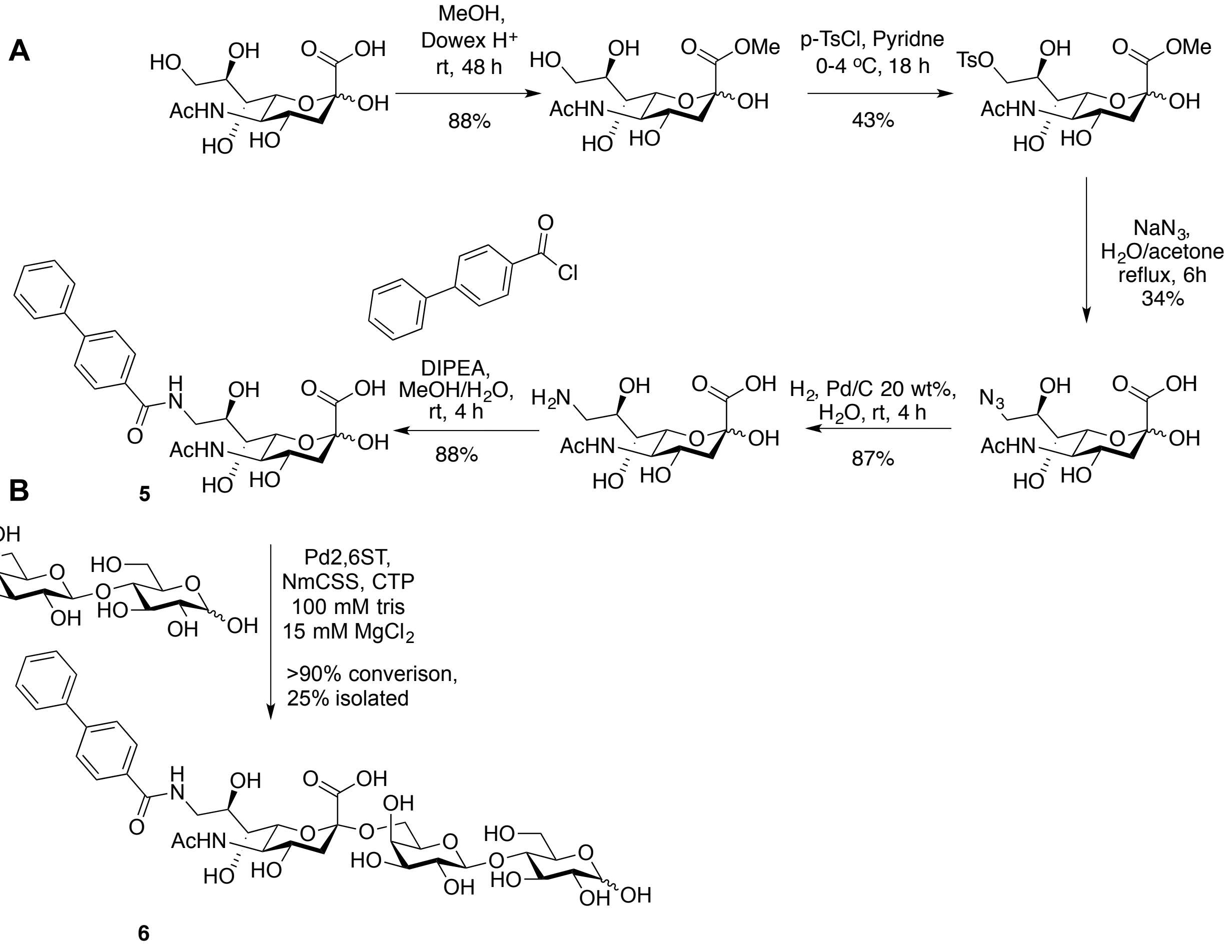

**Supplementary Figure S12. Synthetic Routes for Hybrid Sugars 5 and 6.** (**A**) Synthesis of BPC-Neu5Ac **5** over 5 steps from Neu5Ac. (**B**) Synthesis of **6** from **5**. See **Supplementary Methods** for full details.

**A**

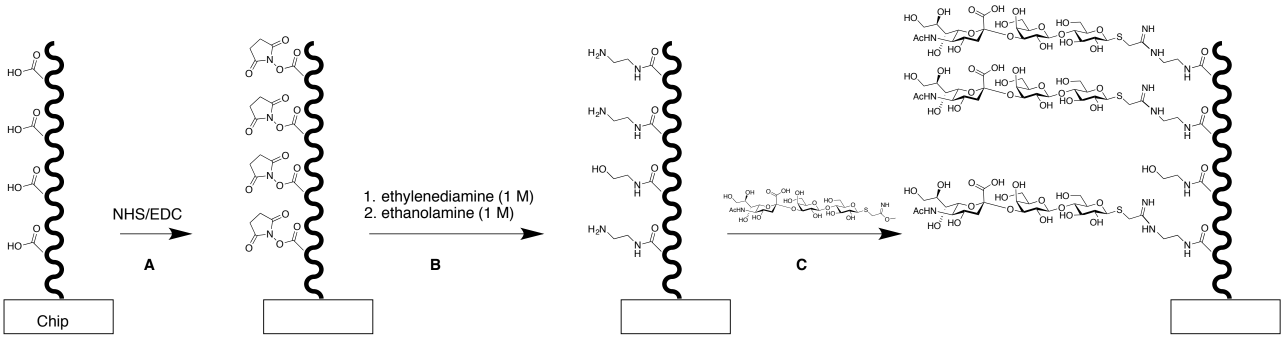

A- CM5 chip activated with NHS (50 mM) and EDC (200 mM) 10 min at 10 uL/min .
B- ethylenediamine (1 M) injected to functionalise chip with free amine groups for 7 min at 10 uL/min.
Then ethanolamine (1 M) was injected over 10 min to block any unreacted NHS activated esters at 10 uL/min.
C- SiaLac-IME (5.6 mM) injected over 10 min at 10 uL/mL followed by washing with HBP-EP buffer (wash steps with buffer between each step and during sensor equilibration).

**B**

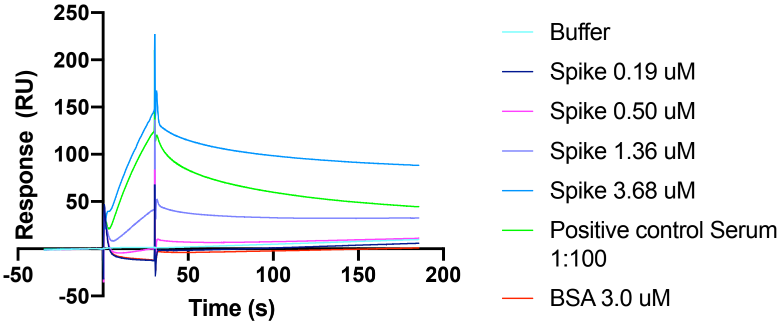

**C**

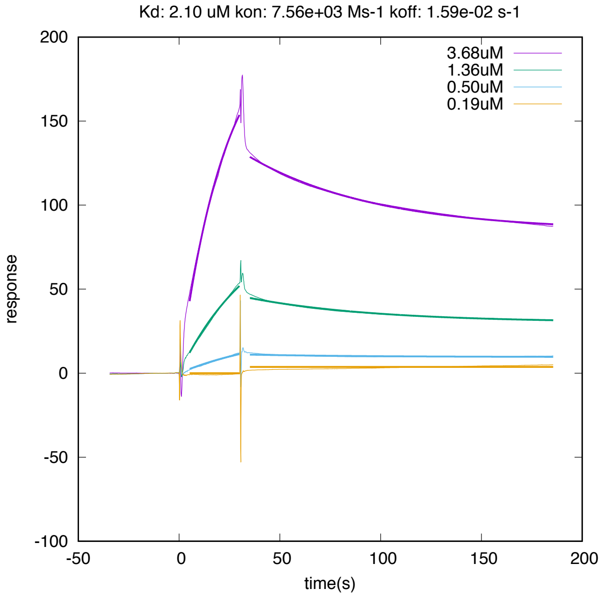

**Supplementary Figure S13. Preparation of SPR Chip and SPR Analysis. A**) Immobilisation of **2** onto carboxymethylated SPR sensor chip. A carboxymethylated CM5 sensor chip was activated with *N*-hydroxysuuccinimide (NHS) and *N*-Ethyl-*N*′-(3-dimethylaminopropyl)carbodiimide hydrochloride (EDC-HCl) and subsequently coupled to ethylenediamine followed by blocking of unreacted esters with ethanolamine. In order to introduce the sugar, a solution of SiaLac-IME was then injected into the flow cell, where it reacts with the surface bound free amines generating an amidine linkage to the glycan (SiaLac-Chip). Specifically, in step A, a CM5 chip activated with NHS (50 mM) and EDC (200 mM) 10 min at 10 μL/min. In step B, ethylenediamine (1 M) was injected to functionalise chip with free amine groups for 7 min at 10 μL/min. Then ethanolamine (1 M) was injected over 10 min to block any unreacted NHS activated esters at 10 μL/min. In step C, SiaLac-

IME (5.6 mM) was injected over 10 min at 10 μL/mL followed by washing with HBP-EP buffer (wash steps with buffer between each step and during sensor equilibration).
**B**) Binding curves of spike at (3.68, 1.36, 0.50 and 0.19 μM) with 30 s association and 150 s dissociation with negative BSA control and positive serum control at 16 °C, corrected by subtracting a buffer only run in a control flow cell.
**C**) Fitted binding curves to determine $K_{on}$ and $K_{off}$ at each spike concentration to determine $K_D$ , corrected by subtracting BSA control. Spike was then flowed over the chip at various concentrations along with a mouse serum as positive control and BSA as a negative control. When accounting for background, the binding curves could be fitted to calculate apparent $k_{on}$ and $k_{off}$ rates giving $K_D$ = 23 μM; $k_{on}$ = 1030 $M^{-1}s^{-1}$; $k_{off}$ = 0.024 $s^{-1}$ from a global analysis of all 4 binding curves.

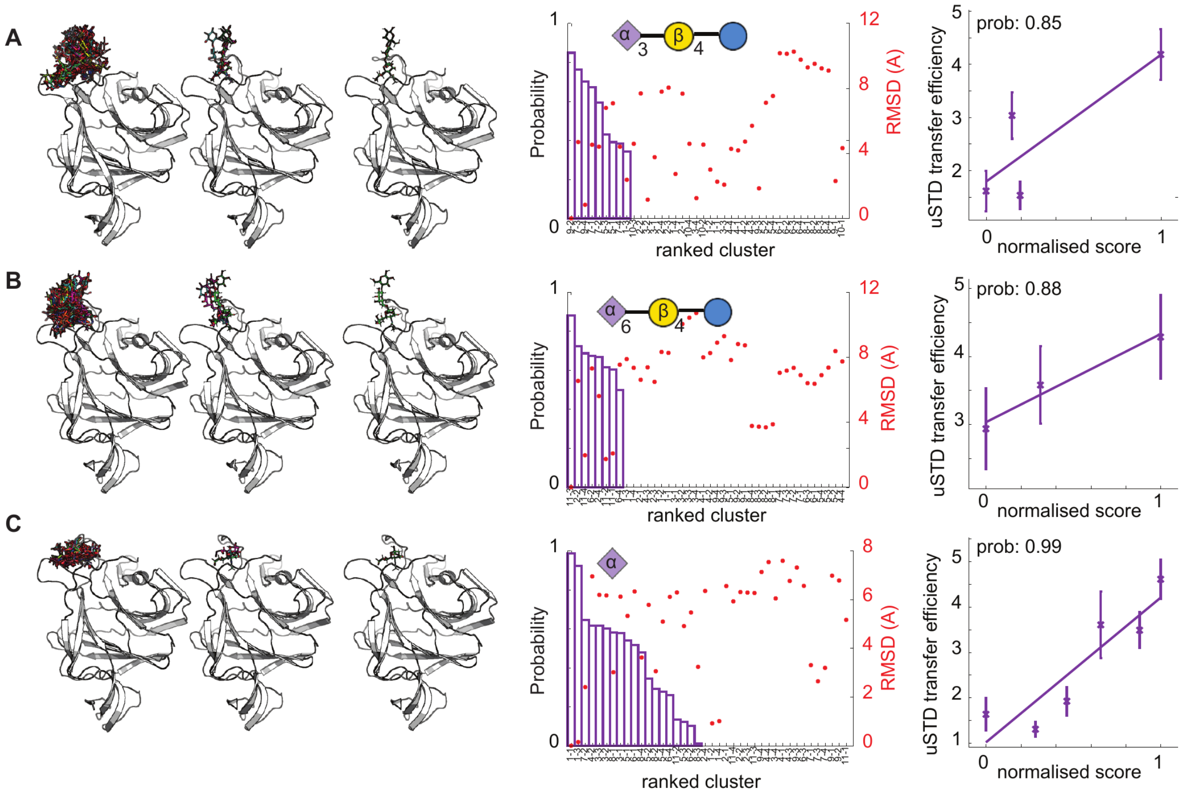

**Supplementary Figure S14. Combination of uSTA with HADDOCK Allowed Ranking of Docked Model Ensembles. A)** A series of models describing the interaction between relevant sugars and spike were calculated using HADDOCK2.4 following standard methods (see **Supplementary Methods**). The docking resulted in 13 clusters ranked by the score of their top four models (i). After comparison to the NMR data, the top 4 members of all clusters were scored based on uSTA data, which allowed selection of 3 very similar models showing excellent agreement with the data (ii) of which the top scorer is shown (iii, main text **Figure 5**). The interactions between Sia2,3Lac and the protein are highly similar for the sialic acid moiety for the top scoring models, with variation occurring predominantly in the flexible Glc residue that has relatively little interaction with the protein. **B)** For each proton in the ligand, the following score was calculated through summing all adjacent protons from the protein: $(<1/r^6>)^{\wedge(1/6)}$. Thus any proton in close proximity to a large number of protons from the protein will receive a high score. This score is expected to correlate with the cross-relaxation rate between the protein and the ligand,[19] and so a high score should correlate with a high transfer efficiency, as measured using STD NMR spectroscopy. The Pearson 'R' correlation coefficient was calculated for each model for the correlation between the score for each atom in the ligand and converted to a probability that the correlation is not generated through random noise,[20] and the transfer efficiency, for the atoms with the highest transfer efficiencies (mapped to 5 heteroatoms, atoms 22,23,24 (Sia) 14 (Gal) and 42 (NAc)). The top-scoring model was an outlier, with a probability level of 98% making it statistically highly significant. The top three models form a cluster of models with a probability >90%, with all falling within 2Å all atom RMSD relative to the model with the highest R value (calculated with the proteins aligned). **C)** The correlation plots between a normalised score and the % transfer efficiency from the uSTA data for the best model (green, i, P(r)=98%), and two excluded models with P(r) values of 75% (blue, ii) and 39% (orange, iii) respectively. Qualitatively, the correlation line passes through the error bars of the selected models (P(r)>90%) and does not for the models that were excluded. Overall, all the top scoring models indicate highly similar interactions between the sialic acid and the protein, and they all predict that the NAc methyl group should be located in the vicinity of the highest proton density from the protein, in agreement with the measurements.

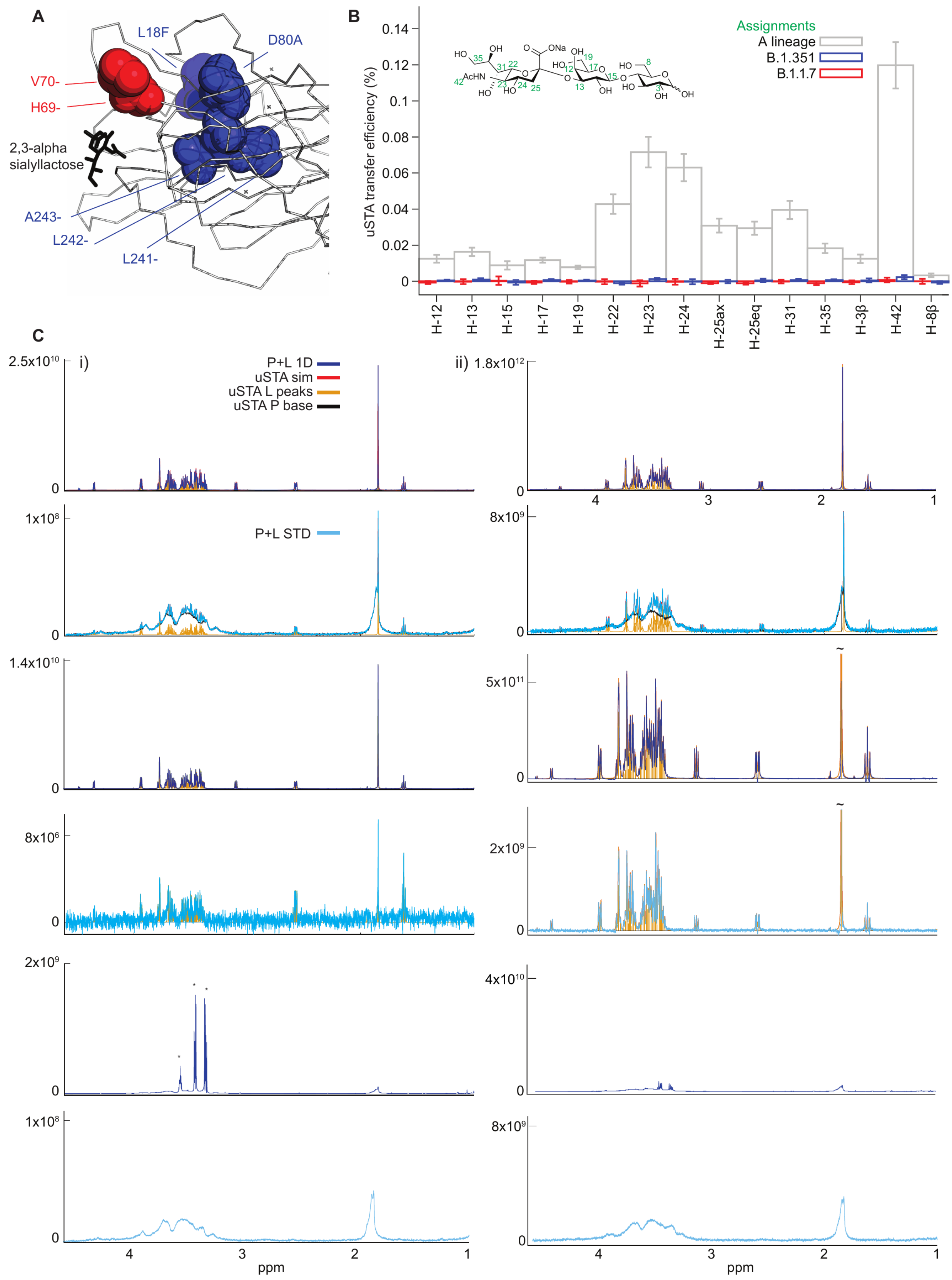

**Supplementary Figure S15: uSTA Analysis of Spike from Variants of Concern. A**) Sialic acid binding pocket indicating site of B.1.1.7 and B.1.351 mutations. **B**) atom specific binding comparison of the three variants showing greatly diminished binding in the B-lineage variants. **C**) raw data for B.1.351 (i) and B.1.1.7 (ii), showing mixture 1D, STD, ligand 1D, STD and protein 1D and STD.

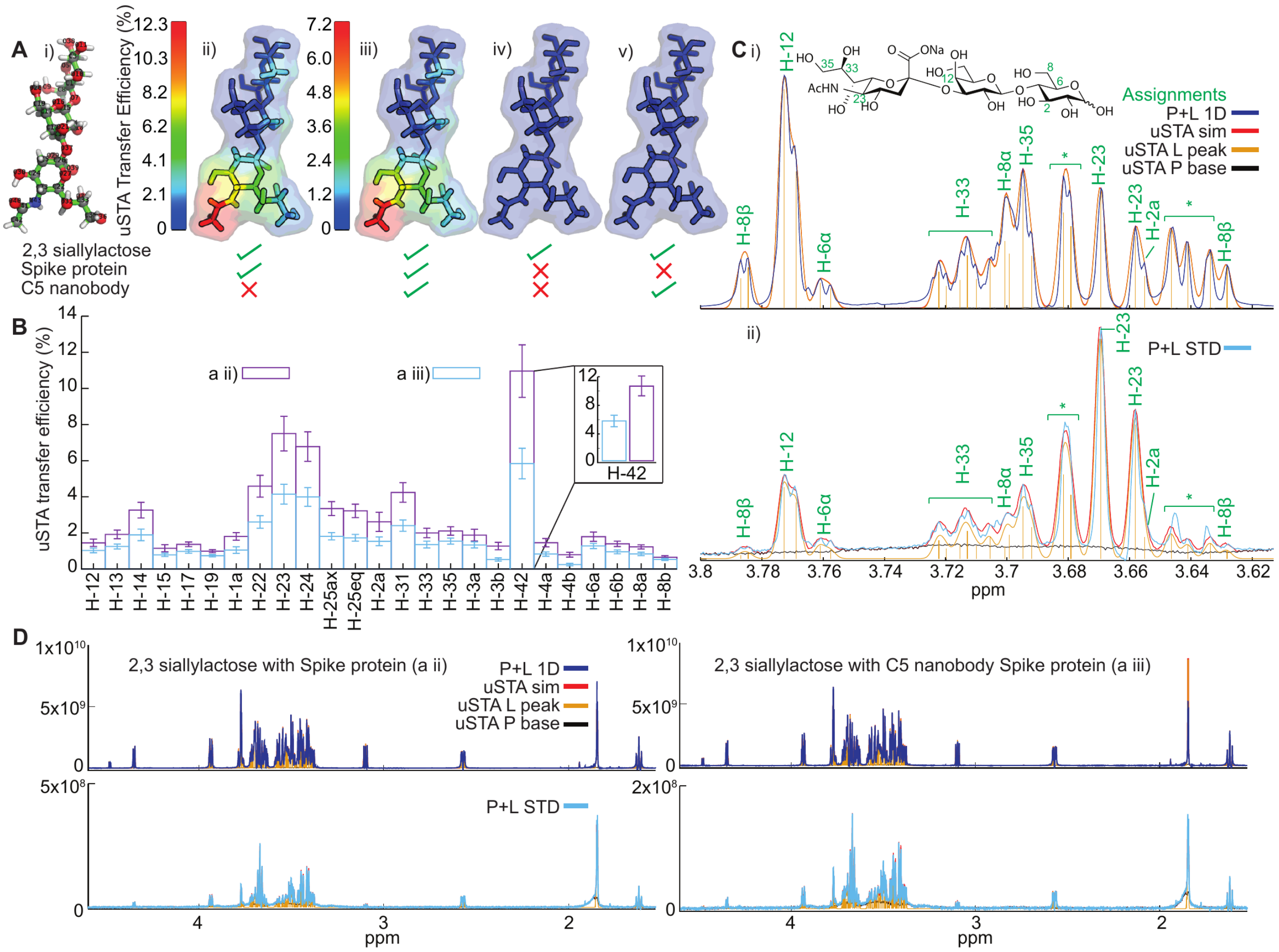

**Supplementary Figure S16: Effects Upon Sialoside Binding of the RBD-Blocking Neutralizing C5 Antibody A i)** a wireframe with atom specific numbering **ii)** heatmap showing the uSTA binding of 2,3 sialyllactose to the SARS-CoV-2 Spike protein **iii)** heatmap showing an almost identical binding pose, but mediated transfer efficiency of the C5 nanobody-bound spike protein **iv)** heatmap showing essentially no uSTA response from 2,3 sialylactose without spike or C5 nanobody present, **v)** heatmap showing essentially no uSTA response from 2,3 sialyllactose with C5 nanobody present but no spike protein. **B**) the uSTA transfer efficiencies of 2,3-sialyllactose in spike and spike and nanobody systems. **C i)** a section of a 1-H spectrum of 2,3-sialyllactose with SARS-CoV-2 Spike protein. **C ii)** difference spectrum of 2,3-sialyllactose with SARS-CoV-2 Spike protein. The assignments are marked in green. * indicates a region with unassignable overlap. **D**) the reference spectra and difference spectra for mixtures without and with C5 nanobody

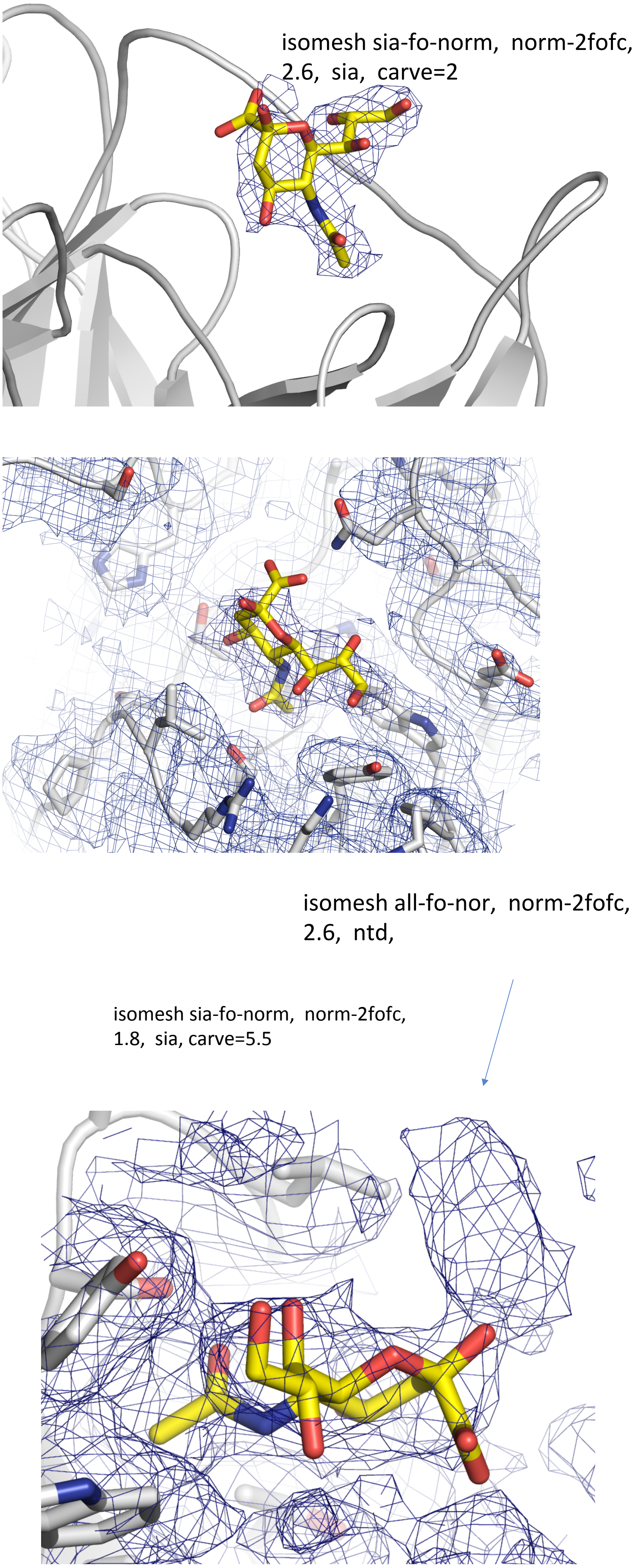

**Supplementary Figure S17. Cryo-EM Coulombic Maps.**

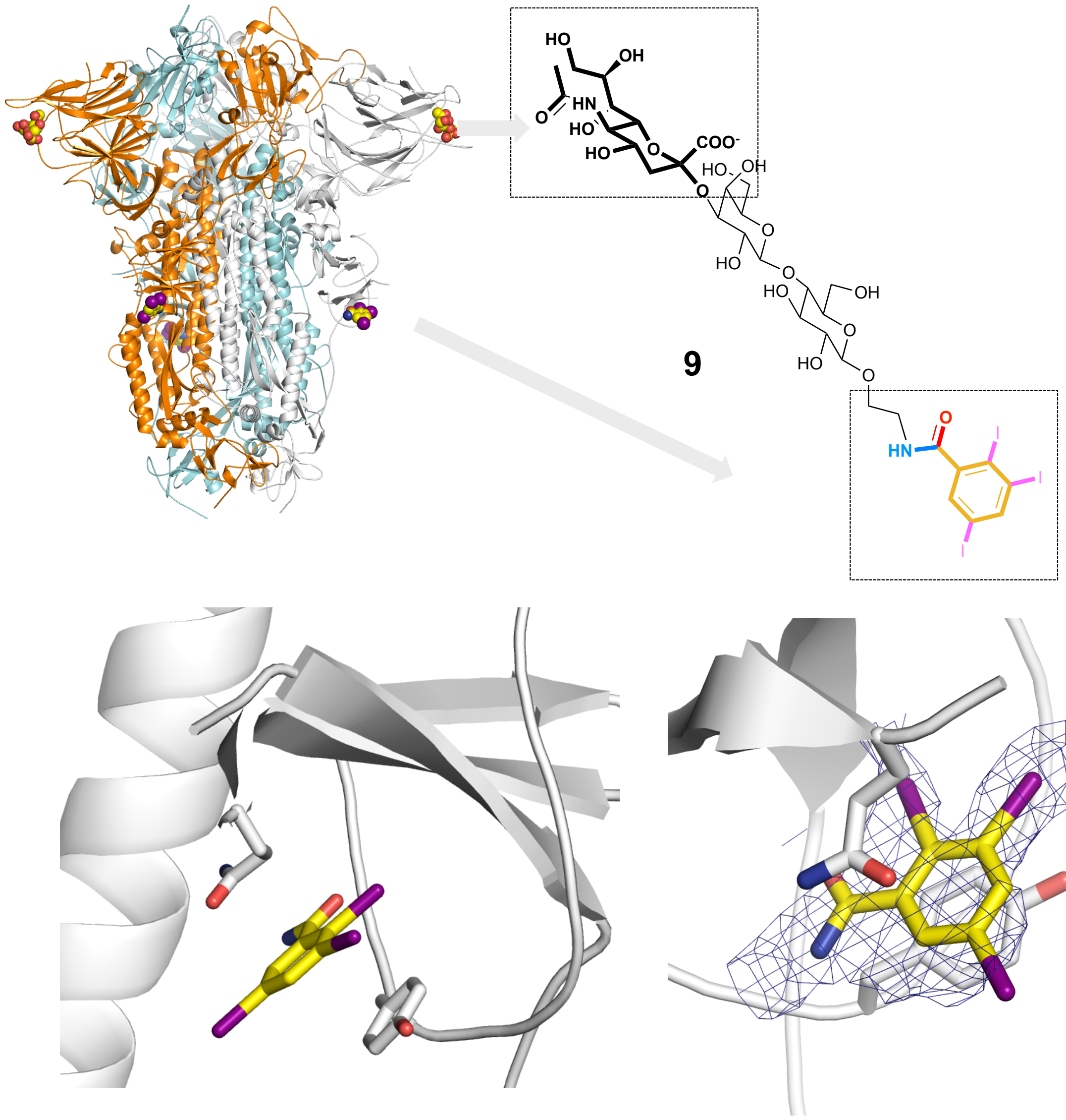

isomesh iod-fo-norm,  norm-2fofc,
4, iodo-mola,  carve=2

**Supplementary Figure S18. Cryo-EM Confirms Additional Binding Mode for Aromatics in a Distinct Region of Spike.**

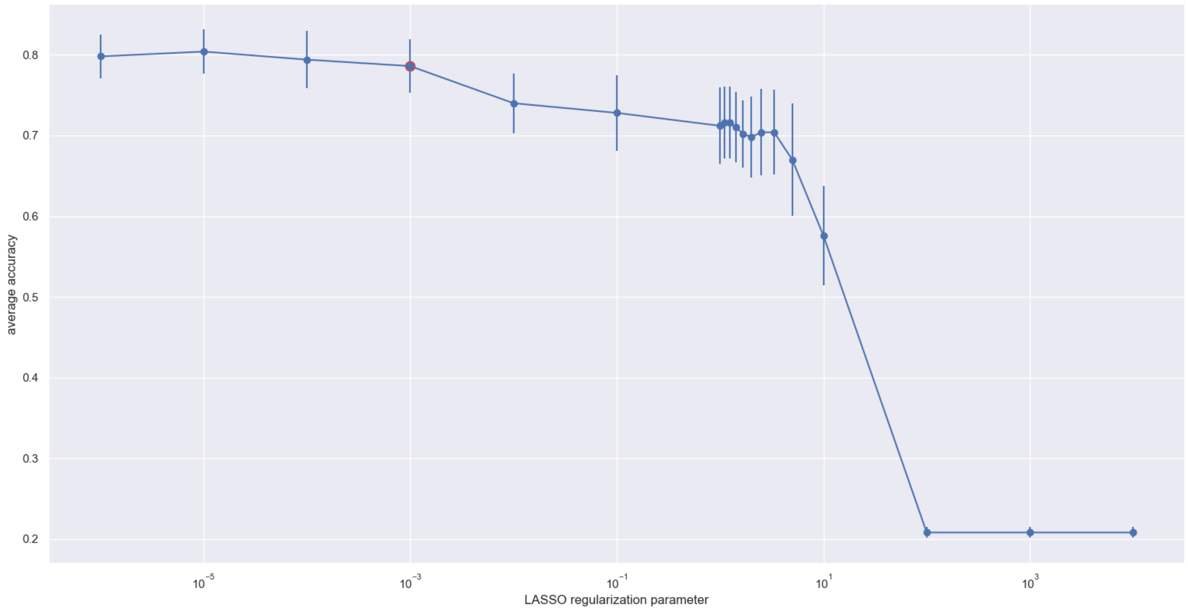

**Supplementary Fig. S19. LASSO Regularization Profile of Clinical Data.**

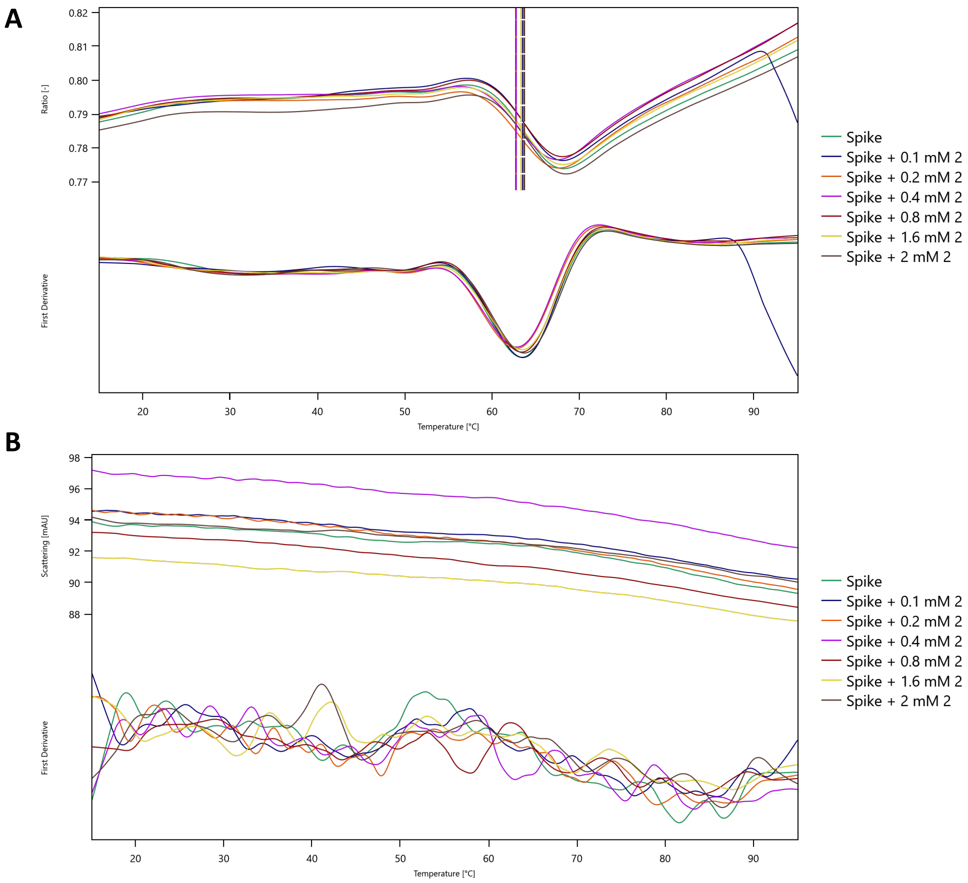

**C**

| Sample | $T_m$ (°C) |
|---|---|
| Spike | 63.4 |
| Spike + 0.1 mM 2 | 63.5 |
| Spike + 0.2 mM 2 | 62.8 |
| Spike + 0.4 mM 2 | 62.8 |
| Spike + 0.8 mM 2 | 63.3 |
| Spike + 1.6 mM 2 | 63.3 |
| Spike + 2.0 mM 2 | 63.6 |

**Supplementary Figure S20**. **Thermal Denaturation Analysis.** Of SARS-CoV-2-spike in the presence of different concentrations of trisaccharide **2**. **A**: Fluorescence ratio at 330/350 nm, indicating the melting temperature of the protein. No significant change in melting temperature is seen in the presence of increasing concentration of trisaccharide **2**. **B**: Light scattering of Spike-BAP observed at increasing concentrations of trisaccharide **2**, used to give an indication of protein aggregation. No aggregation of Spike-BAP is seen at increasing concentrations of different concentrations of trisaccharide **2**. **C**: Melting temperatures of Spike-BAP in the presence and absence of different concentrations of trisaccharide **2**.

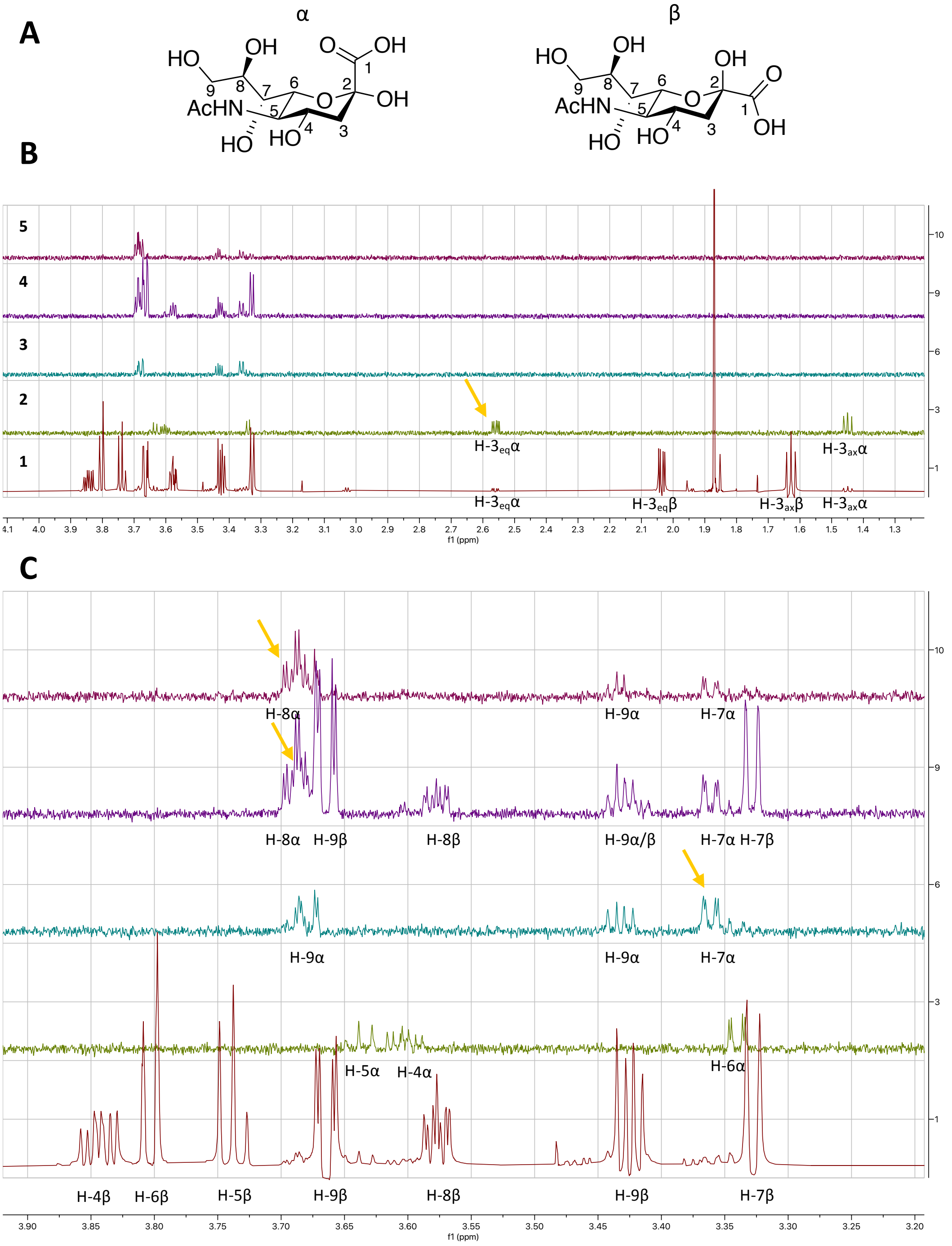

**Supplementary Figure S21. Manual Assignment of Alpha Anomer of Neu5Ac. A)** Alpha and beta anomers of Neu5Ac with atom numbers labelled. **B)** Overlays of proton and 1D TOCSY spectra indicating H-3 beta positions which can be observed easily in the $^{1}$H proton (**1**) and H-3 alpha positions which can be seen in the 1D TOCSY (**2**). Irradiated frequencies- **2**= 2.550 ppm, **3**= 3.361 ppm, **4**= 3.680 ppm, **5**= 3.695 ppm (indicated as yellow arrows on spectra). **C)** Assigned peaks corresponding to alpha and beta anomers. Peaks were assigned by following coupling constants for each proton around the sugar ring.

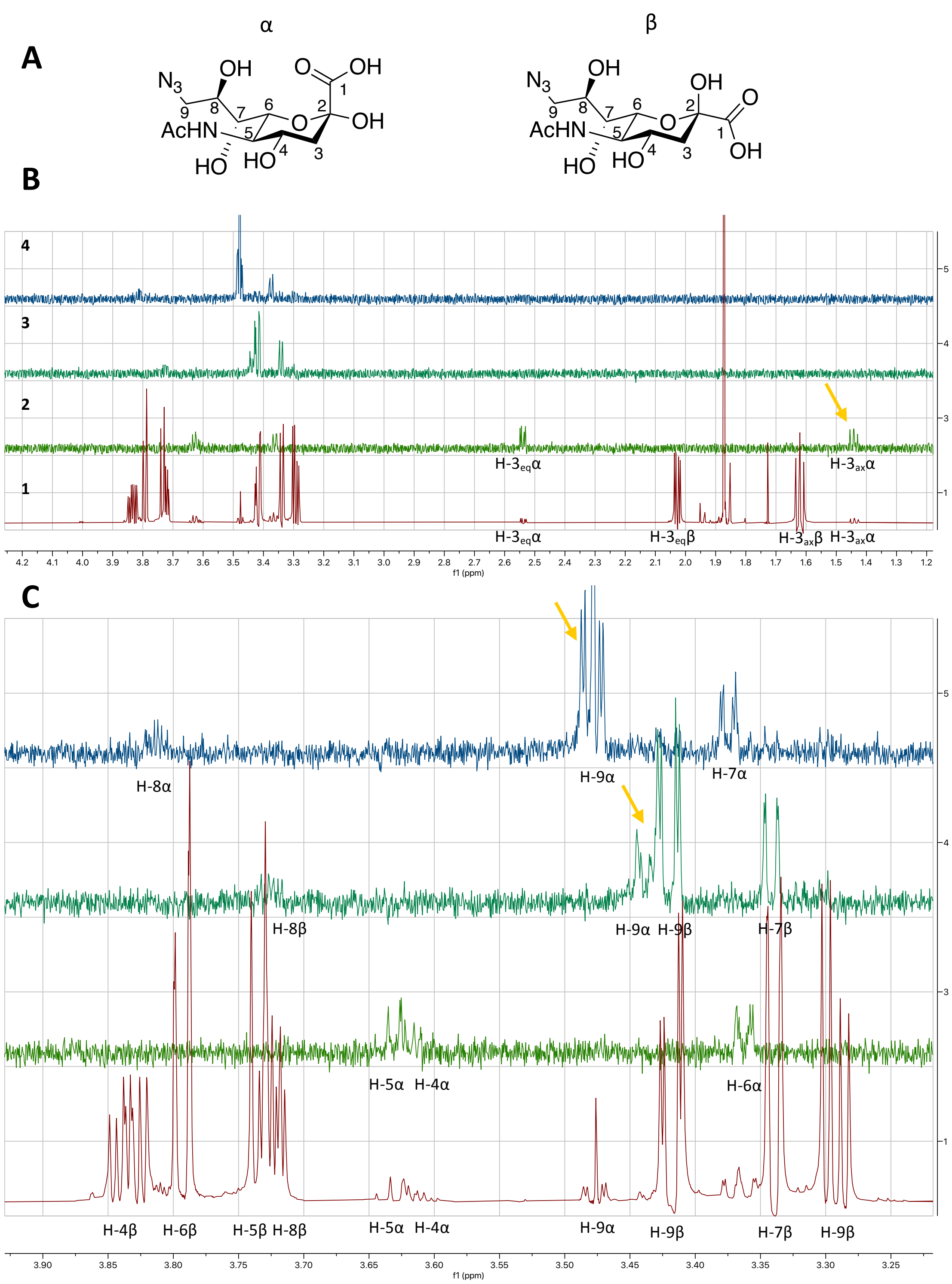

**Supplementary Figure S22. Manual Assignment of Alpha Anomer of 9-azido-Neu5Ac. A**) Alpha and beta anomers of 9-azido-Neu5Ac with atom numbers labeled. **B**) Overlays of proton and 1D TOCSY spectra indicating H-3 beta positions which can be observed easily in the $^{1}$H proton (**1**) and H-3 alpha positions which can be seen in the 1D TOCSY (**2**). Irradiated frequencies- **2**= 1.438 ppm, **3**= 3.442 ppm, **4**= 3.477 ppm (indicated as yellow arrows on spectra). **C**) Assigned peaks corresponding to alpha and beta anomers. Peaks were assigned by following coupling constants for each proton around the sugar ring.

**Supplementary Tables**

**Supplementary Table S1**. COVID-19 Cohort
**Supplementary Table S2**. *B3GNT8* and *LGALS3BP* genetic variants
**Supplementary Table S3**. *B3GNT8* chi-square five categories
**Supplementary Table S4**. *B3GNT8* chi-square 2x2
**Supplementary Table S5**. *LGALS3BP* chi-square five categories
**Supplementary Table S6**. *LGALS3BP* chi-square 2x2
**Supplementary Table S7**. Raw data and UniDecNMR fits used to calculate the uSTA surfaces shown in the manuscript.

As separate xlsx:

**Supplementary Table S1.** COVID-19 Cohort
**Supplementary Table S2.** *B3GNT8* and *LGALS3BP* genetic variants
S**upplementary Table S3.** *B3GNT8* chi-square five categories
**Supplementary Table S4.** *B3GNT8* chi-square 2x2
**Supplementary Table S5.** *LGALS3BP* chi-square five categories
**Supplementary Table S6.** *LGALS3BP* chi-square 2x2

**Supplementary Table S7: a-i). Raw data and UniDecNMR fits used to calculate the uSTA surfaces shown in the manuscript.** For each protein/ligand combination, the uSTA transfer efficiency surfaces were calculated from the protein only, ligand only and mixture STD datasets. A variety of impurities (*) were observed in the various spectra and were accounted for during analysis as described below. Small resonances coming from impurities are commonly detected in 1D $^{1}$H NMR spectra. For the most part, these were of low intensity and did not overlap with the ligand resonances. In two cases, impurities were detected perhaps from cryoprotectants used during the protein preparation, and their concentration was sufficiently high that the underlying ligand resonances could not be resolved. In each spectrum, the symbol ~ indicates where intensity has been cut-off, in order to scale the spectrum so that resonances of low dynamic range can be clearly discerned.
The specific impurities encountered across the various spectra are tabulated below, together with a summary of their individual effects on the spectra.

| | $^{1}$H chemical shift | Features | Present in |
|---|---|---|---|
| **I1** | 1.71 ppm (s) | Negative STD when excited at 5.4 ppm or 8 ppm | a) mixture, protein<br>b) mixture, protein only<br>c) ligand only, mixture<br>d) ligand only, mixture<br>e) all<br>f) all<br>g) ligand only |
| **I2** | 7.93 ppm (d) | Positive STD ligand only at 8 ppm, no STD when excited at 5.4 ppm | a) ligand only, mixture<br>b) ligand only |
| **I3** | Glycerol intense peaks: 3.46 ppm (dd),<br>3.37 ppm (dd),<br>3.57-3.62 ppm (m) | Negligible STD when excited at 5.4 ppm | b) mixture |
| **I4** | 2.02 ppm (s) | Negligible STD when excited at 5.4 ppm | a) protein only<br>c) protein only<br>d) protein only |
| **I5** | 3.16 ppm (s) | Negative STD when excited at 5.4 ppm | c) mixture, ligand only |
| **I6** | 3.536 ppm (s),<br>3.45-3.497 ppm (q) | Positive STD at 5.4 ppm | e) mixture |
| **I7** | 3.42 ppm (dd),<br>3.26 ppm (dd) | No STD | g/h/i) protein only, mixture |
| **I8** | 3.8 ppm (s) | No STD | g/h/i) ligand only |
| **I9** | 0.98 ppm (d) | No STD | g/h/i) protein only, mixture |

**a) 2,3-sialyllactose and SARS-CoV-2 Spike Protein**
Mixture 1 & 2, Ligand 2, Protein 1 & 4: no overlap with ligand, zero impact on analysis.
**b) 2,6-sialyllactose and SARS-CoV-2 Spike Protein**
Mixture 1, Ligand 2, Protein 1: no overlap with ligand, zero impact on analysis.
Mixture 3: Glycerol coming from protein batch used. Due to its high intensity (the intensity is 10x higher than the resonances from the ligand), the region was excluded.
Excluded atoms: 24, 31, 13, 14, 2β, 1β, 6β, 3α, 1α
We would expect an appreciable STD response for 24 and 31, and close to zero for the others. We can see evidence for this in the spectrum, but we can cannot accurately quantify due to the large glycerol resonance that covers this region. While we have no measurement of 24 and 31, we have

measurements of the surrounding protons which we can use to reasonably interpolate these values (see Methods).

**c) Sialic acid and SARS-CoV-2 Spike Protein**

Mixture 1 & 5, Ligand 1 & 5, Protein 4: no overlap with ligand, zero impact on analysis.

**d) 9-azido-sialic acid and SARS-CoV-2 Spike Protein**

Mixture 1, Ligand 1, Protein 4: no overlap with ligand, zero impact on analysis.

**e) 9-BPC-2,6-sialyllactose and SARS-CoV-2 Spike Protein**

Mixture 1, Ligand 1, Protein 1: no overlap with ligand, zero impact on analysis.

Mixture 6: coming from the ligand batch used. Due to its high intensity (the intensity is 10x higher than the resonances from the ligand), the region was excluded.

Excluded atoms: 24, 13

In the STD spectra, a modest STD response can be discerned for 24 and 13 whose magnitude is comparable to adjacent atoms. Although we cannot accurately measure the transfer efficiency, the interpolation will be accurate.

**f) 9-BPC-sialic acid and SARS-CoV-2 Spike Protein**

Mixture 1, Ligand 1, Protein 1: no overlap with ligand, zero impact on analysis.

**g), h) and i) various concentrations of tryptophan and BSA**

Mixture 7 & 9, Ligand 8, Protein 7 & 9: no overlap with ligand, zero impact on analysis.

a) 2,3-sialyllactose and SARS-CoV-2 Spike protein

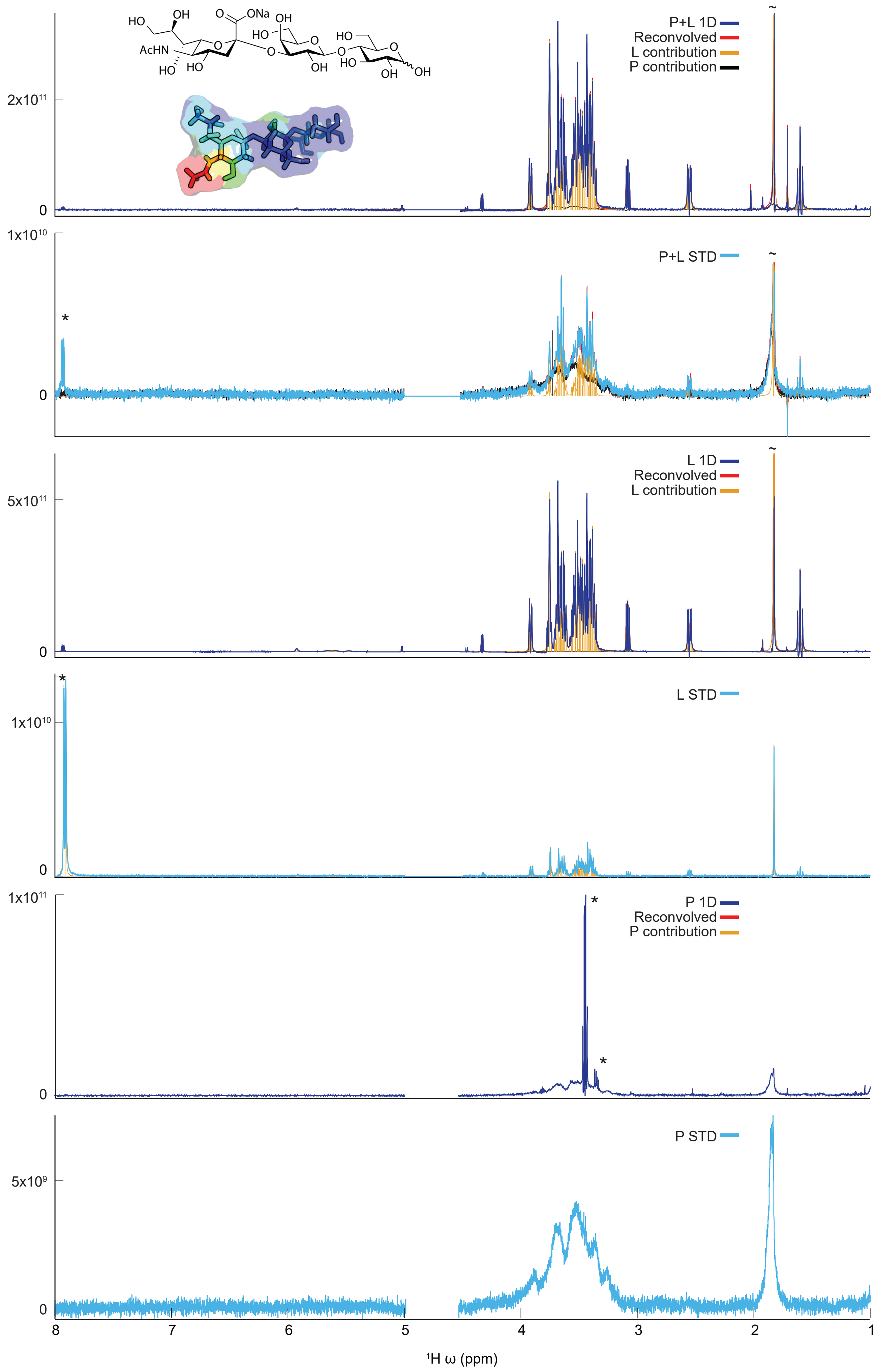

b) 2,6-sialyllactose and SARS-CoV-2 Spike protein

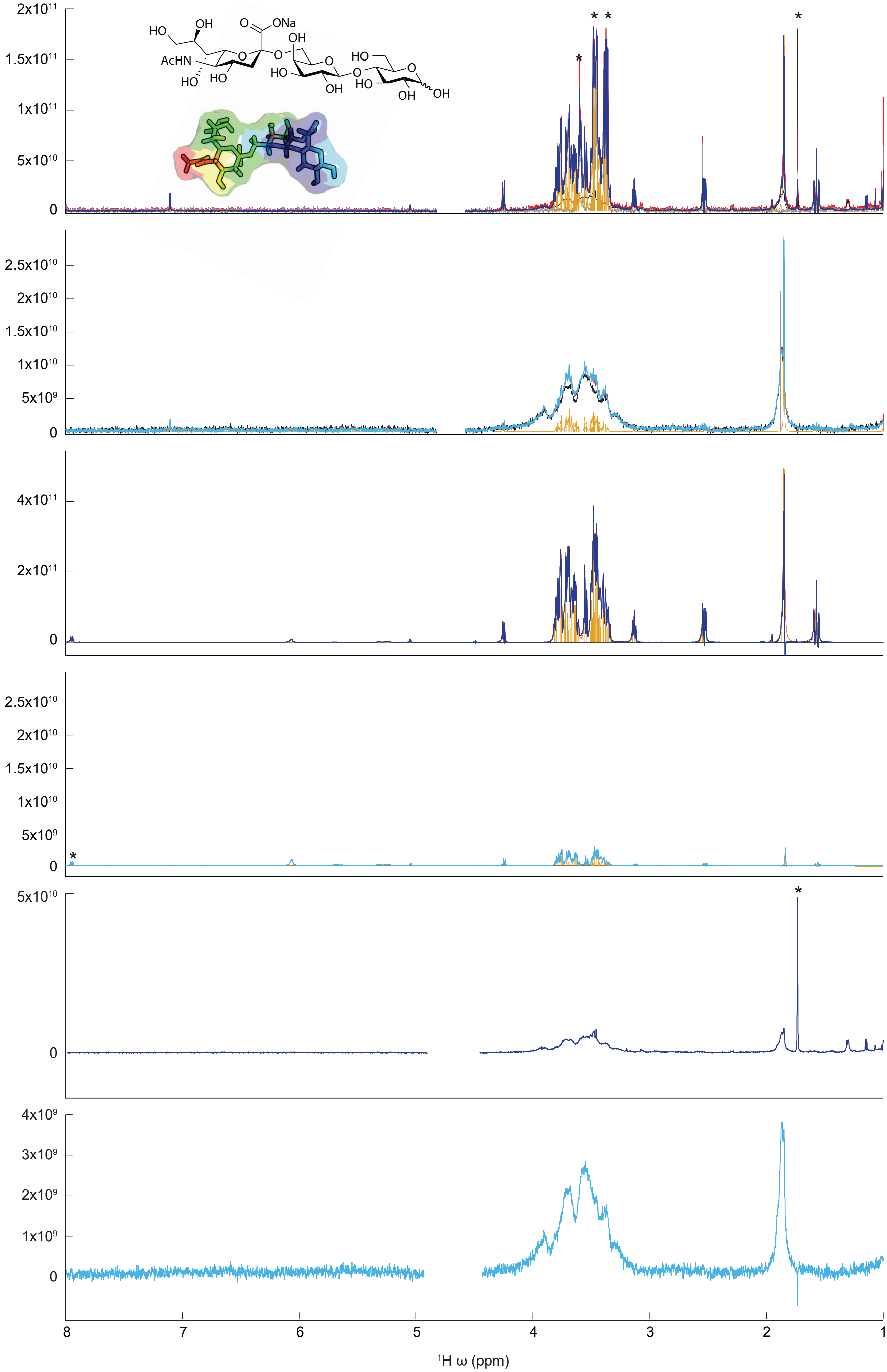

c) Sialic acid and SARS-CoV-2 Spike protein

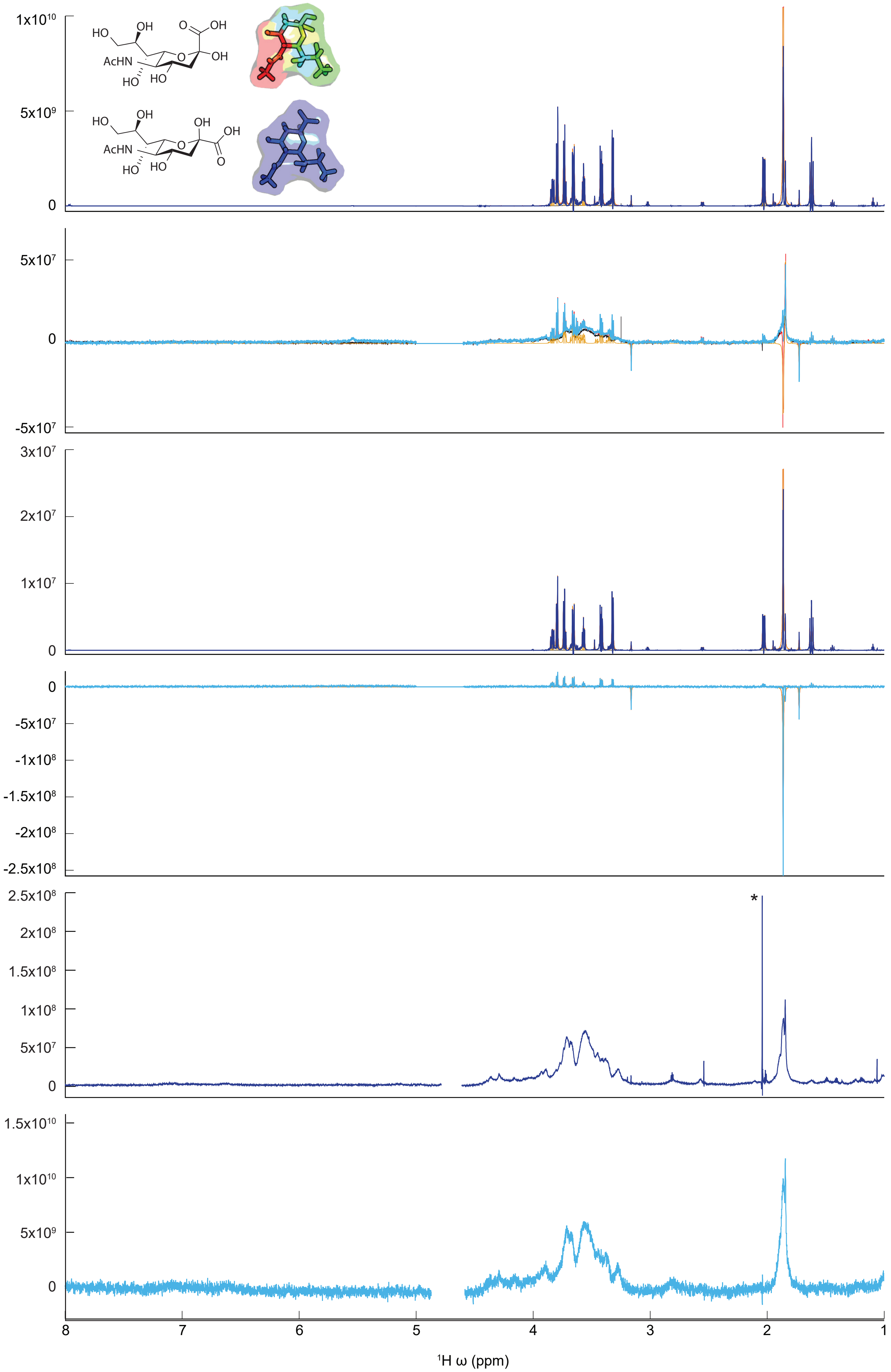

d) 9-azidosialic acid and SARS-CoV-2 Spike protein

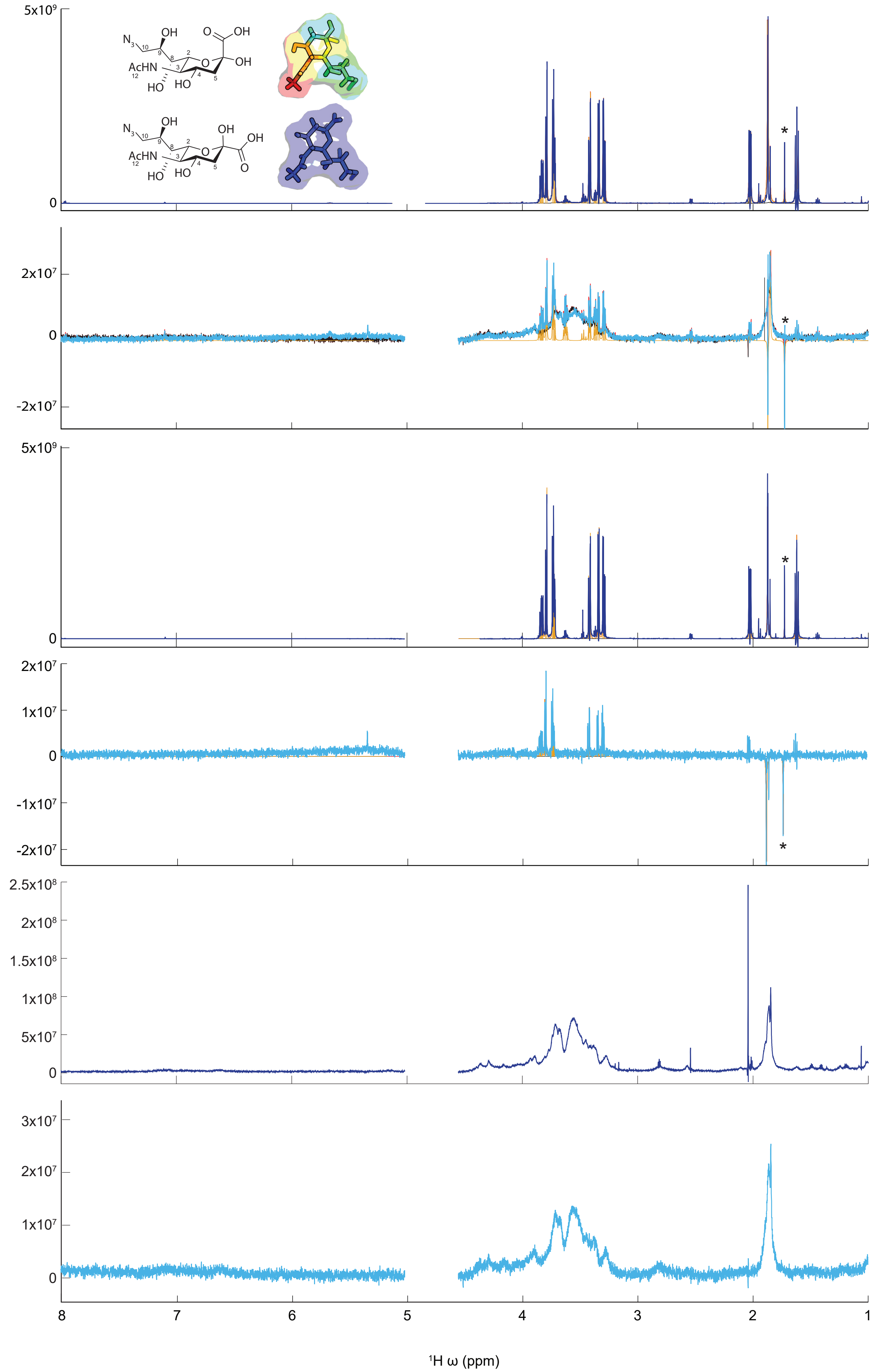

e) 9-BPC2,6-sialyllactose and SARS-CoV-2 Spike protein

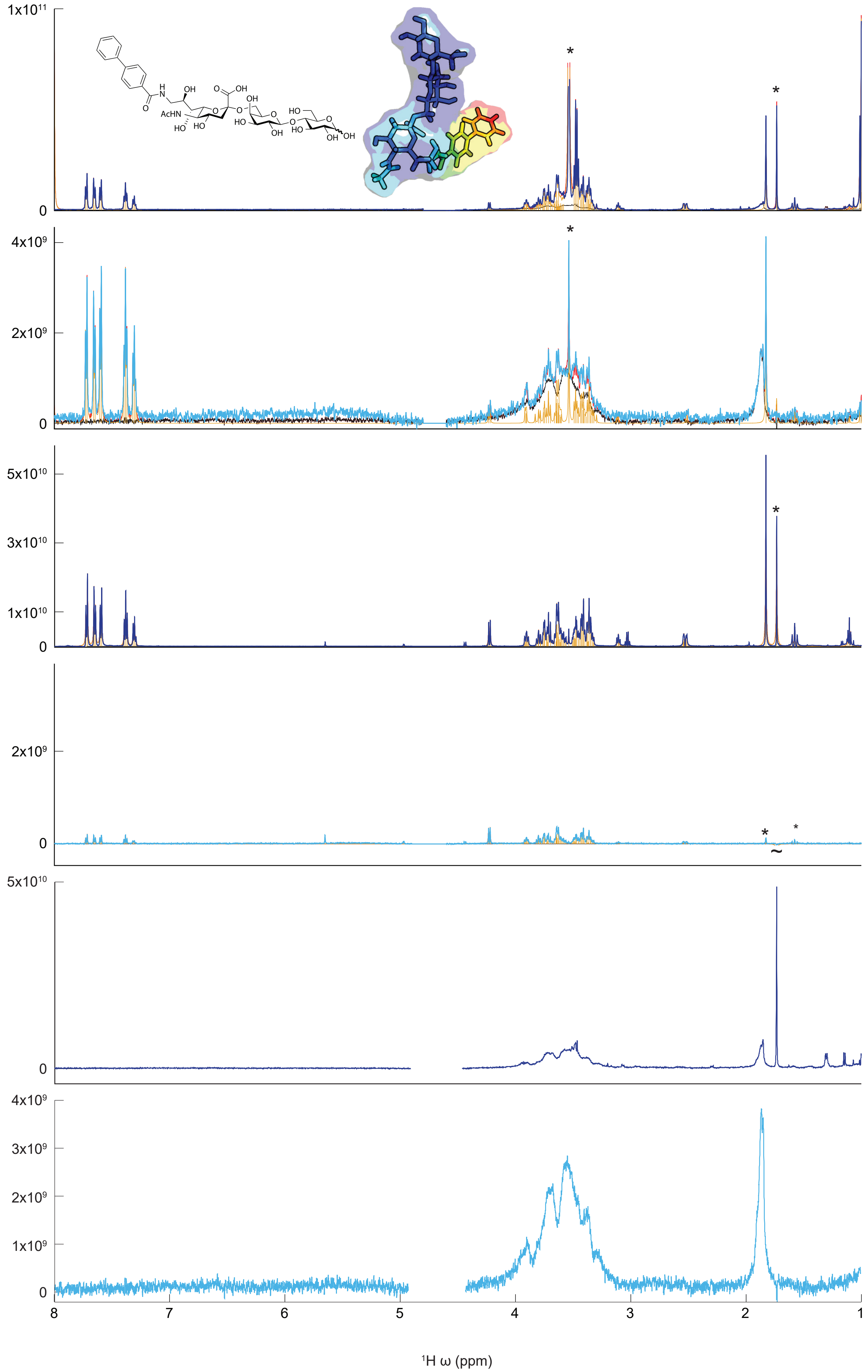

f) 9-BPCsialic acid and SARS-CoV-2 Spike protein

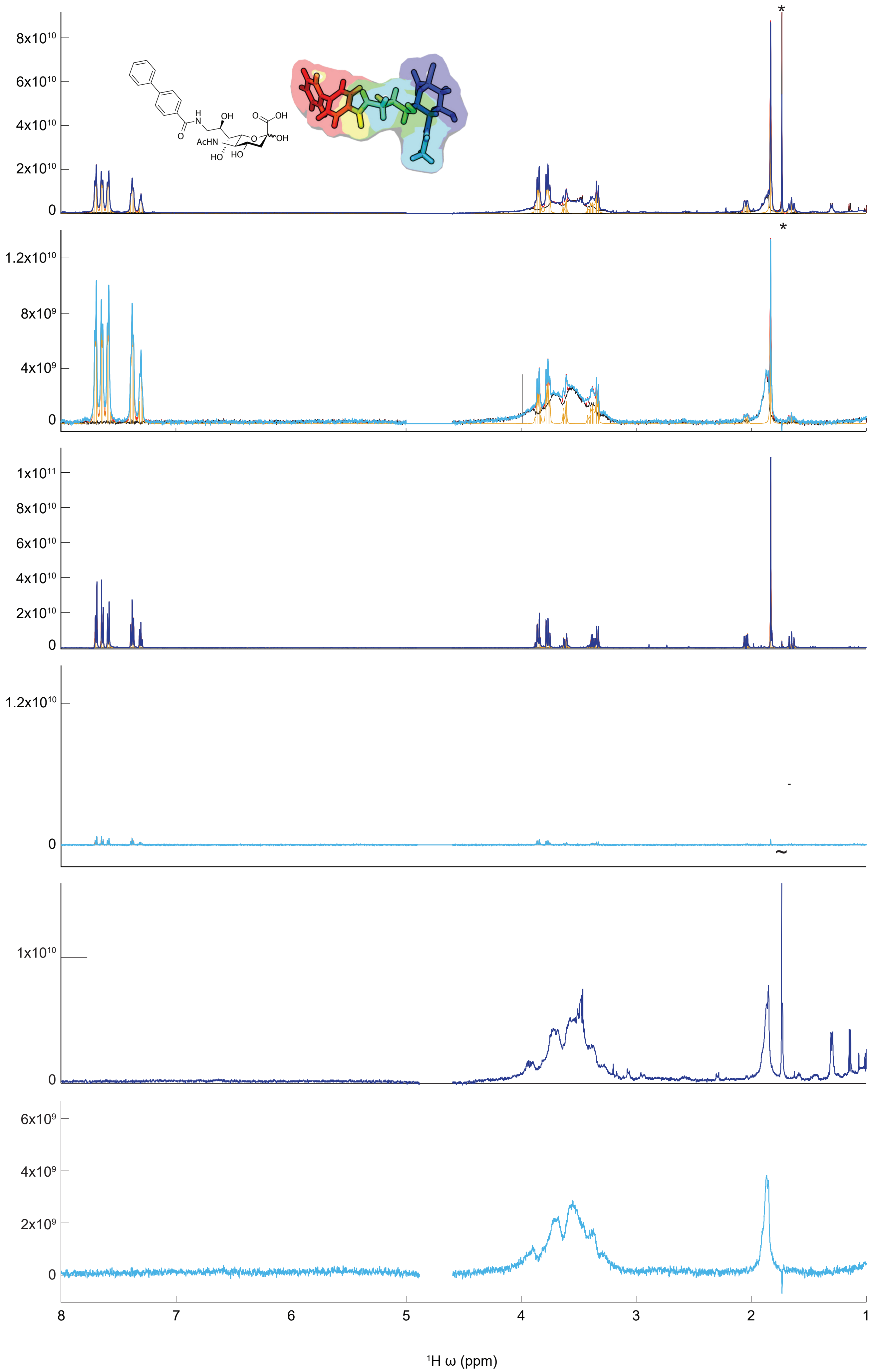

g) 40μM tryptophan and BSA

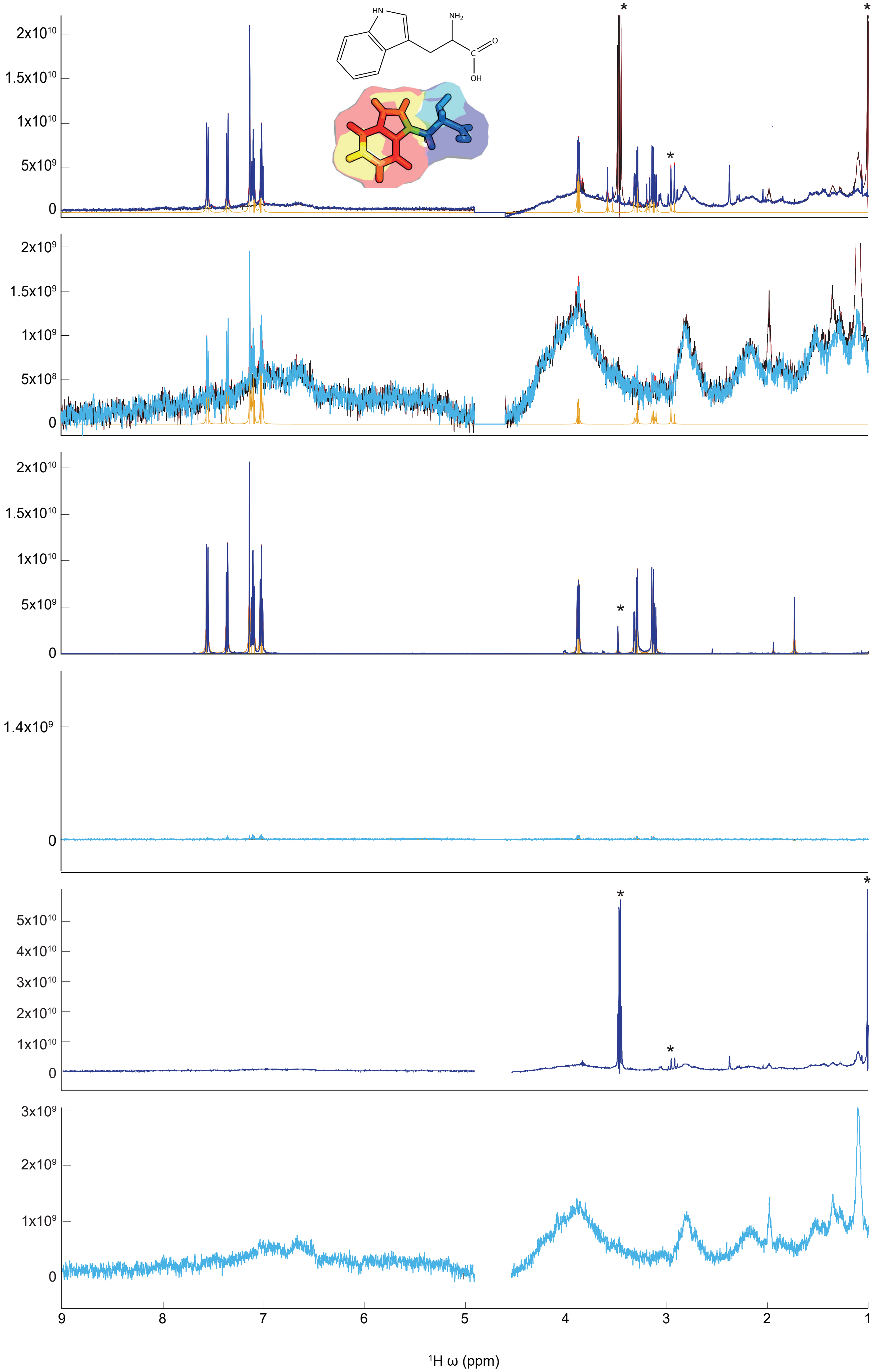

h) 200μM tryptophan and BSA

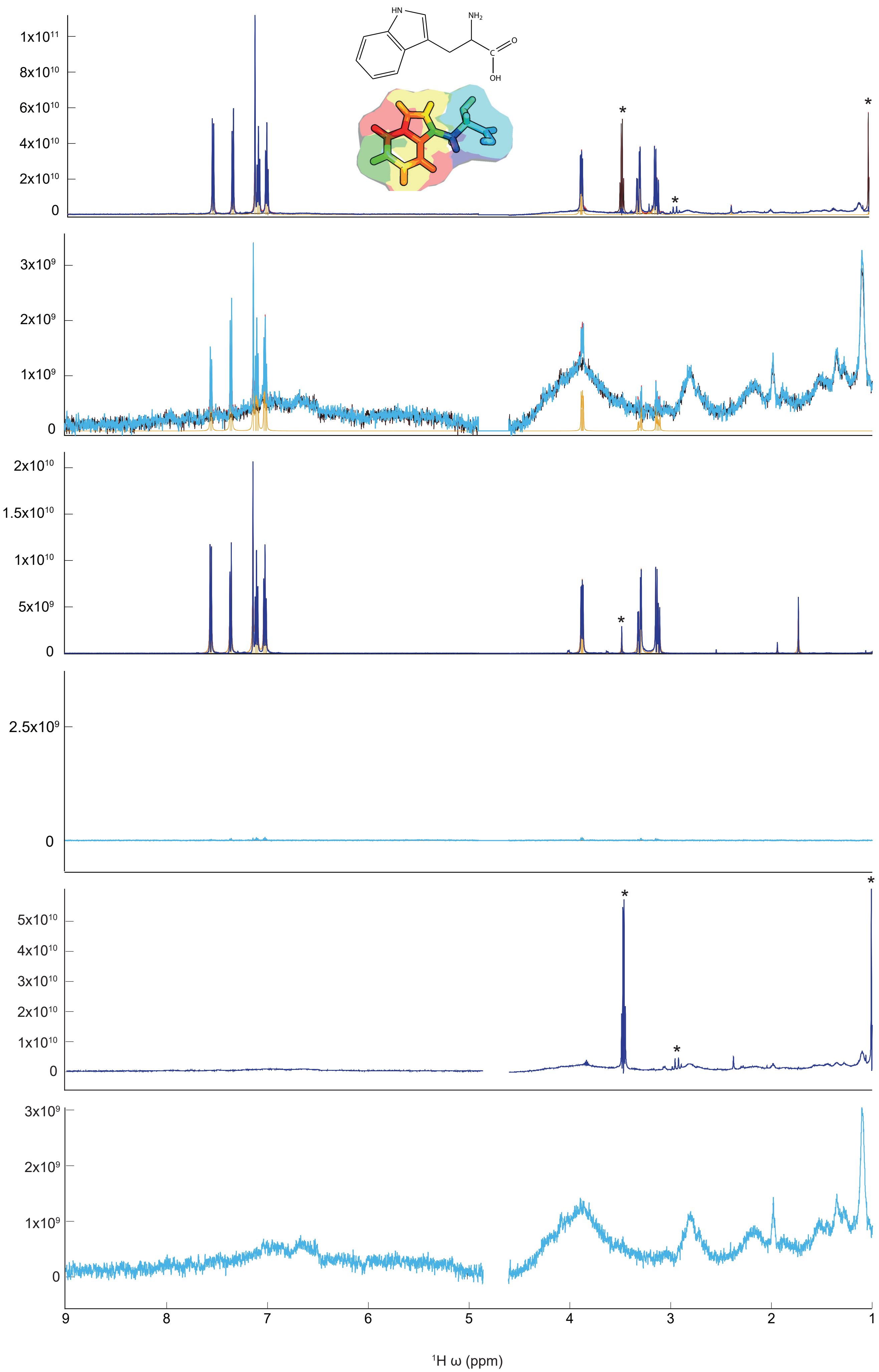

i) 1mM tryptophan and BSA

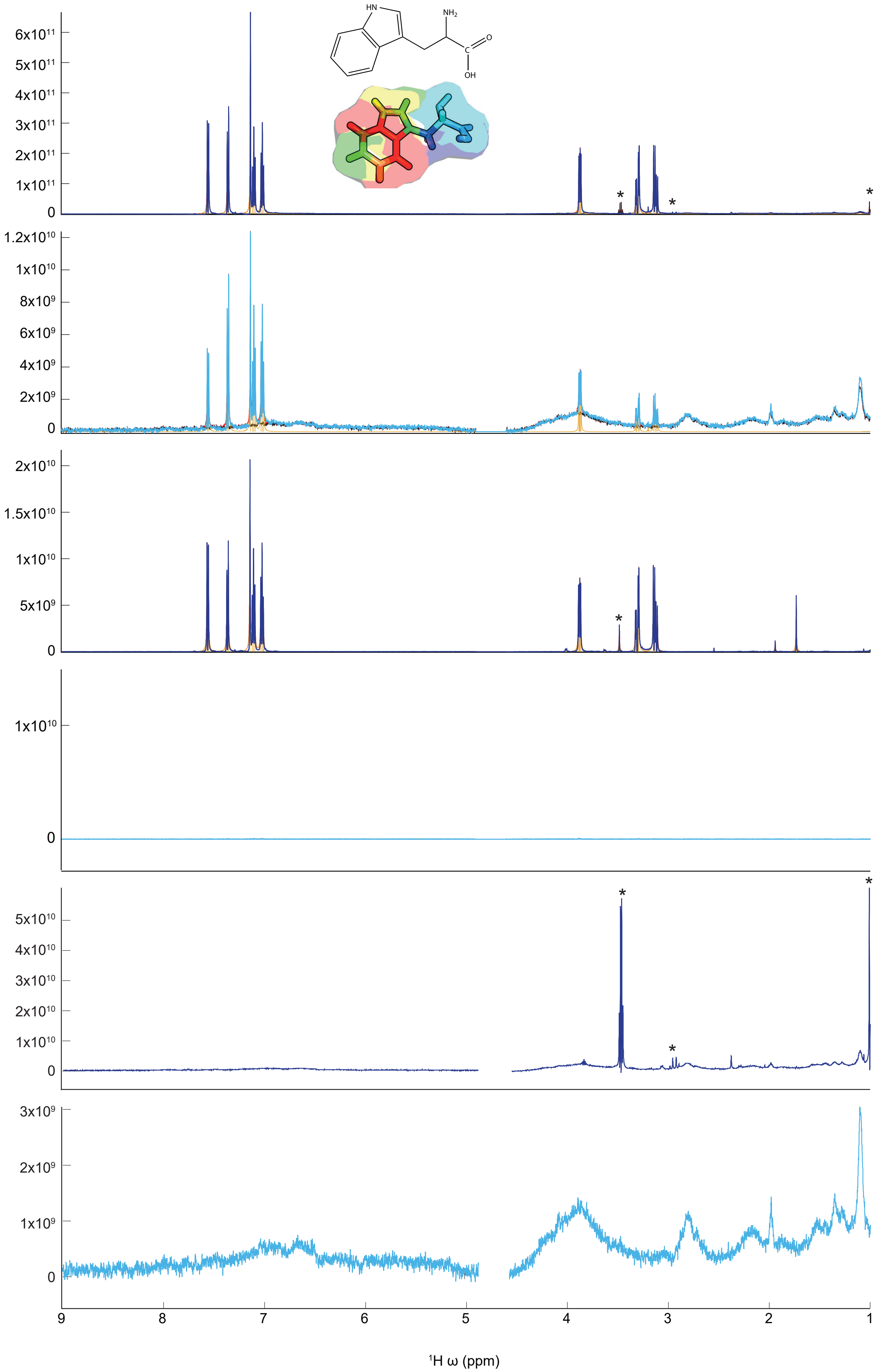
